# Bivariate causal mixture model quantifies polygenic overlap between complex traits beyond genetic correlation

**DOI:** 10.1101/240275

**Authors:** Oleksandr Frei, Dominic Holland, Olav B. Smeland, Alexey A. Shadrin, Chun Chieh Fan, Steffen Maeland, Kevin S. O’Connell, Yunpeng Wang, Srdjan Djurovic, Wesley K. Thompson, Ole A. Andreassen, Anders M. Dale

## Abstract

Accumulating evidence from genome wide association studies (GWAS) suggests an abundance of shared genetic influences among complex human traits and disorders, such as mental disorders. While current cross-trait analytical methods focus on genetic correlation between traits, we developed a novel statistical tool (MiXeR), which quantifies polygenic overlap independent of genetic correlation, using summary statistics from GWAS. MiXeR results can be presented as a Venn diagram of unique and shared polygenic components across traits. At 90% of SNP-heritability explained for each phenotype, MiXeR estimates that more than 9K variants causally influence schizophrenia, 7K influence bipolar disorder, and out of those variants 6.9K are shared between these two disorders, which have high genetic correlation. Further, MiXeR uncovers extensive polygenic overlap between schizophrenia and educational attainment. Despite a genetic correlation close to zero, these traits share more than 9K causal variants, while 3K additional variants only influence educational attainment. By considering the polygenicity, heritability and discoverability of complex phenotypes, MiXeR provides a more complete quantification of shared genetic architecture than offered by other available tools.

## INTRODUCTION

In recent years, genome-wide association studies (GWASs) have successfully detected genetic variants associated with multiple complex human traits or disorders, providing important insights into human biology^1^. Understanding the degree to which complex human phenotypes share genetic influences is critical for identifying the etiology of phenotypic relationships, which can inform disease nosology, diagnostic practice and improve drug development. Most human phenotypes are known to be influenced by multiple genetic variants, many of which are expected to influence more than one phenotype (i.e. exhibit allelic pleiotropy)^2,3^. This has led to cross-trait analyses, quantifying genetic overlap, becoming a widespread endeavor in genetic research, made possible by the public availability of most GWAS summary statistics (p-values and z-scores)^4,5^.

Currently, the prevailing measure to quantify genetic overlap is genetic correlation. The sign of the correlation indicates whether the shared genetic effects predominantly have the same or the opposite effect directions. Available methods can quantify genetic correlation using raw genotypes^6,7^or GWAS summary statistics^8-10^. However, these methods report overall positive, negative or no genetic correlation, but do not capture mixtures of effect directions across shared genetic variants. This scenario is exemplified by the genetic relationship between schizophrenia and educational attainment. Despite consistent estimates of a non-significant genetic correlation^11,12^, many genetic loci are found to be jointly associated with both phenotypes^13^. Among 25 shared loci^14^, 16 had effects in the opposite direction, while 9 had effects in the same direction. Thus, new statistical tools are needed to improve our understanding of the polygenic architecture of complex traits and their intricate relationships.

Here we developed a statistical tool (MiXeR), which quantifies polygenic overlap independent of genetic correlation, using summary statistics from GWAS. To evaluate polygenic overlap between two traits MiXeR estimates the total number of shared and trait-specific causal variants (i.e. variants with non-zero additive genetic effect on a trait). MiXeR bypasses the intrinsically difficult problem of detecting the exact location of causal variants, but rather aims at estimating their overall amount. MiXeR builds upon univariate causal mixture^15-18^, extending the model to four bivariate normal distributions as illustrated in Figure 1, with two causal components specific to each trait; one causal component of variants affecting both traits; and a null component of variants with no effect on either trait. From the prior distribution of genetic effects, we derive likelihood function of the observed signed test statistics (GWAS z-scores), incorporating effects of linkage disequilibrium (LD) structure, minor allele frequency, sample size, cryptic relationships, and sample overlap. The parameters of the mixture model are estimated from the summary statistics by direct optimization of the likelihood function.

**Figure 1.**
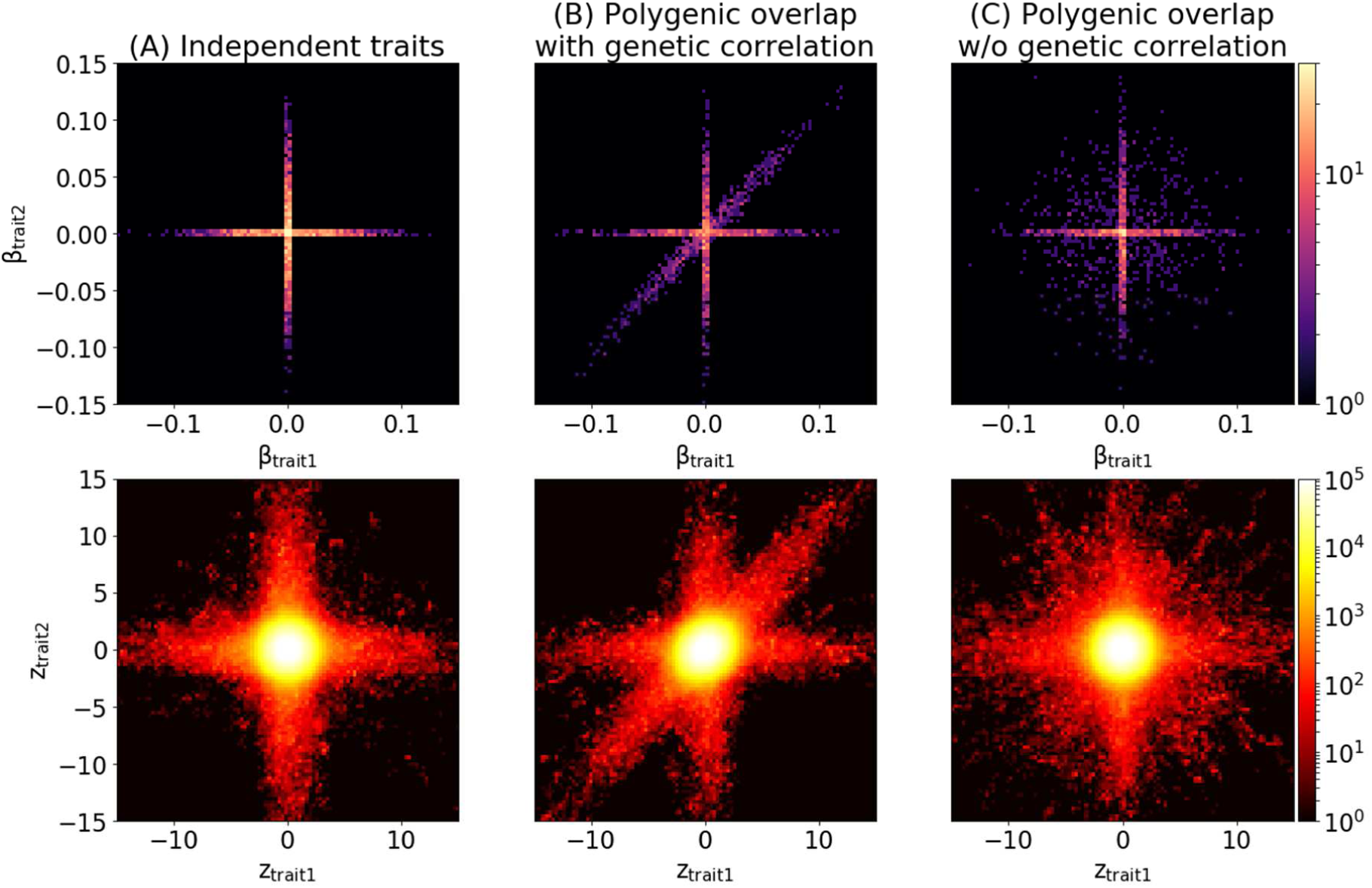
Components of the bivariate mixture in three scenarios of polygenic overlap. All figures are generated from synthetic data, where causal variants were drawn from MiXeR model, total polygenicity in each trait set to 0.01%, SNP heritability set to 0.4, GWAS N=100 000. First column shows two traits where causal variants do not overlap. Second column adds a component of causal variants affecting both traits in the same (concordant) direction. Third column shows scenario of polygenic overlap without genetic correlation. Top row shows simulated bivariate density of additive effects of allele substitution (β_1j_, β_2j_), bottom row shows bivariate density of GWAS signed test statistics (z_1j_, z_2j_) for GWAS SNPs (genotyped or imputed). Due to Linkage Disequilibrium, GWAS signed test statistic has substantially larger volume of SNPs associated with the phenotype. The aim of MiXeR model is to infer distribution of causal effects (top row), using GWAS data (bottom row) as an input. Figures are generated on a regular grid of 100 × 100 bins, color histogram indicates log10(N) where N is the number of SNPs projected into a bin.

We show in simulations that MiXeR provides accurate estimates of model parameters in the presence of realistic LD structure. Using GWAS summary data, we quantify polygenic overlap of several psychiatric disorders, including schizophrenia and bipolar disorder, with educational attainment and human height, with large implications for understanding how genetic factors overlap between complex human phenotypes.

## RESULTS

### Simulations studies

In our first set of simulations we generate synthetic GWAS data that follow model assumptions and check validity of MiXeR estimates (polygenic overlap, *π*_12_; correlation of effect sizes within the shared polygenic component, *p*_12_, and genetic correlation, rg) in the presence of realistic LD structure (Figure 2). We observe no bias in the estimates across a wide range of simulation scenarios (Supplementary Figure 1-3), except for a specific scenario with correlated effect sizes (*p*_12_=0.5) and high polygenicity 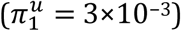. In this case polygenic overlap (*π*_12_) is underestimated, while correlation of effect sizes *p*_12_ is overestimated by the same factor, so that the estimated genetic correlation remains unbiased. This bias in *π*_12_ and *p*_12_ estimates is attributed to complete sample overlap. Additionally, the scenario with low heritability (*h*_2_ = 0.1) and high polygenicity 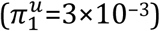 shows large variation among estimates, which is due to low GWAS signal. Standard errors estimated by the model are shown in Supplementary Table 1.

**Table 1.** Results of cross-trait analysis with MiXeR model for schizophrenia (SCZ), bipolar disorder (BIP), educational attainment (EDU) and height GWAS. Columns: *n*_12_ – estimated number of shared causal variants, reported in 1,000; *n*_1_ (*n*_2_)– estimated number of causal variants, unique to trait1 (trait2), expressed in 1,000; *p*_12_ – correlation of effect sizes in shared polygenic component; rg – genetic correlation (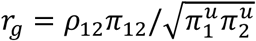, see Online Methods); rg_LDSR_ - estimate of genetic correlation from LD Score Regression. Number of variants (*n*_12_, *n*_1_ and *n*_2_) are adjusted to explain 90% of heritability in the corresponding component. Parameters are fitted using ca. 1.1M HapMap3 SNPs.

| trait1 | trait2 | $n_{12}$ (se) | $n_1$ (se) | $n_2$ (se) | $\rho_{12}$ (se) | rg (se) | rg<sub>LDSR</sub> (se) |
| --- | --- | --- | --- | --- | --- | --- | --- |
| SCZ | BIP | 6.88 (1.04) | 2.25 (1.34) | 0.36 (0.43) | 0.837 (0.019) | 0.708 (0.062) | 0.717 (0.024) |
| SCZ | EDU | 9.09 (0.92) | 0.08 (0.04) | 2.94 (1.12) | 0.056 (0.015) | 0.048 (0.014) | 0.079 (0.022) |
| SCZ | Height | 1.09 (0.12) | 8.12 (0.97) | 2.72 (0.14) | -0.007 (0.057) | -0.001 (0.011) | -0.008 (0.019) |
| BIP | EDU | 6.04 (1.34) | 1.07 (0.94) | 5.99 (1.49) | 0.308 (0.054) | 0.201 (0.030) | 0.176 (0.026) |
| BIP | Height | 1.08 (0.13) | 6.05 (1.24) | 2.71 (0.15) | 0.000 (0.064) | 0.000 (0.013) | -0.011 (0.023) |
| EDU | Height | 1.83 (0.11) | 10.19 (0.64) | 2.02 (0.11) | 0.442 (0.034) | 0.119 (0.009) | 0.141 (0.012) |

**Figure 2.**
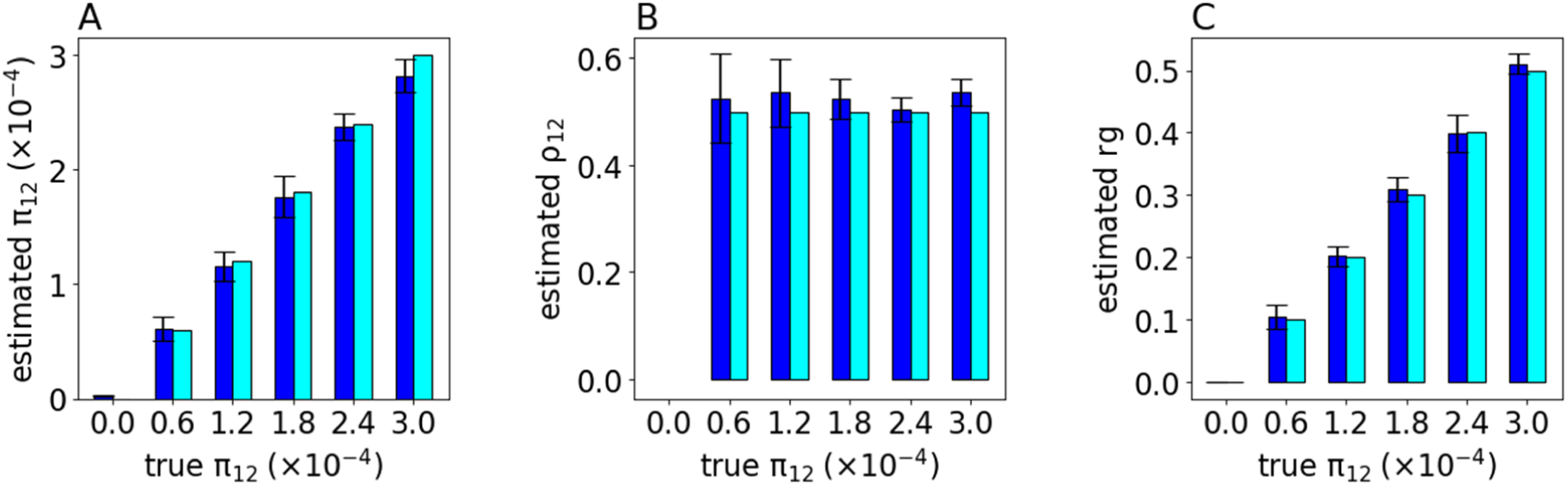
Selected simulations with bivariate model: (A) estimates of polygenic overlap; (B) estimates of correlation of the effect sizes in shared polygenic component; (C) estimates of genetic correlation. The bars in blue indicate an average value of model estimates across 10 simulation runs. The bars in cyan show true (simulated) parameters. Error bars represent standard deviation of the model estimate across 10 simulation runs. Different bars correspond to levels of polygenic overlap: from zero (no overlap) to complete polygenic overlap. Simulated heritability is 0.4, simulated fraction of causal variants is 0.03% in both traits.

Additionally we show that univariate estimates of polygenicity and heritability are correct in all scenarios except when heritability is low and polygenicity is high (Supplementary Figure 4, Supplementary Table 2). This case corresponds to insufficiently powered GWAS, yielding large variation among parameters, which leads to the bias from truncation, as the polygenicity parameter is bound to be non-negative.

**Figure 4.**
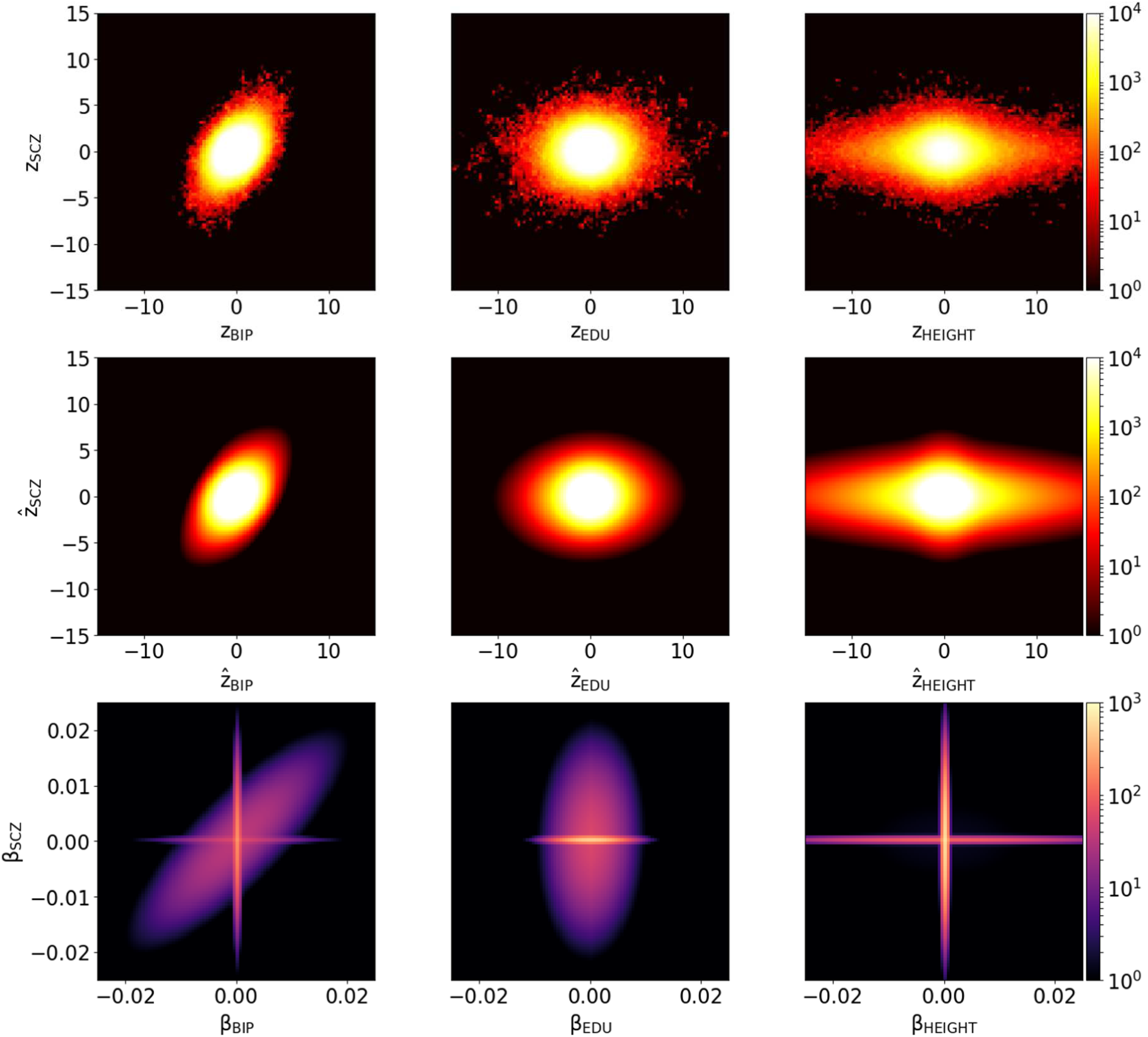
Top row shows bivariate density of the observed GWAS signed test statistics (*z*_1j_, *z*_2j_), middle row shows predicted density 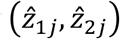 from MiXeR model. Bottom row shows estimated bivariate density of additive causal effects (*β*_1j_, *β*_2j_) that underlie model prediction. Three columns represent schizophrenia (SCZ) versus bipolar disorder (BIP), educational attainment (EDU) and height GWAS. Density is visualized using regular grid of 100 × 100 bins, color indicates log10(N) where N is the observed number (for the top row) or the expected number (for the middle and bottom rows) of SNPs projected into a bin.

Finally, we validate that the model accurately predicts GWAS quantile-quantile (Q-Q) plots (Supplementary Figure 5) and detailed Q-Q plots with SNPs partitioned into disjoint groups according to minor allele frequency (MAF) and LD score (Supplementary Figure 6a,b). Detailed Q-Q plots show a stronger GWAS signal for SNPs with higher MAF and higher LD score. The model’s prediction follows the same pattern, indicating that it correctly captures dependency of GWAS association statistics on MAF and LD score. Conditional Q-Q plots (Supplementary Figure 7) show observed versus expected -log10 p-values in the primary trait as a function of significance of association with a secondary trait at the level of p≤0.1, p≤0.01, p≤0.001, with data Q-Q plots being closely reproduced by the model’s predictions. Interestingly, scenarios without polygenic overlap are also showing a minor enrichment, arising because GWAS p-values depend on allele frequency and LD structure, though this effect is generally smaller than enrichment arising due to shared causal variants.

**Figure 5.**
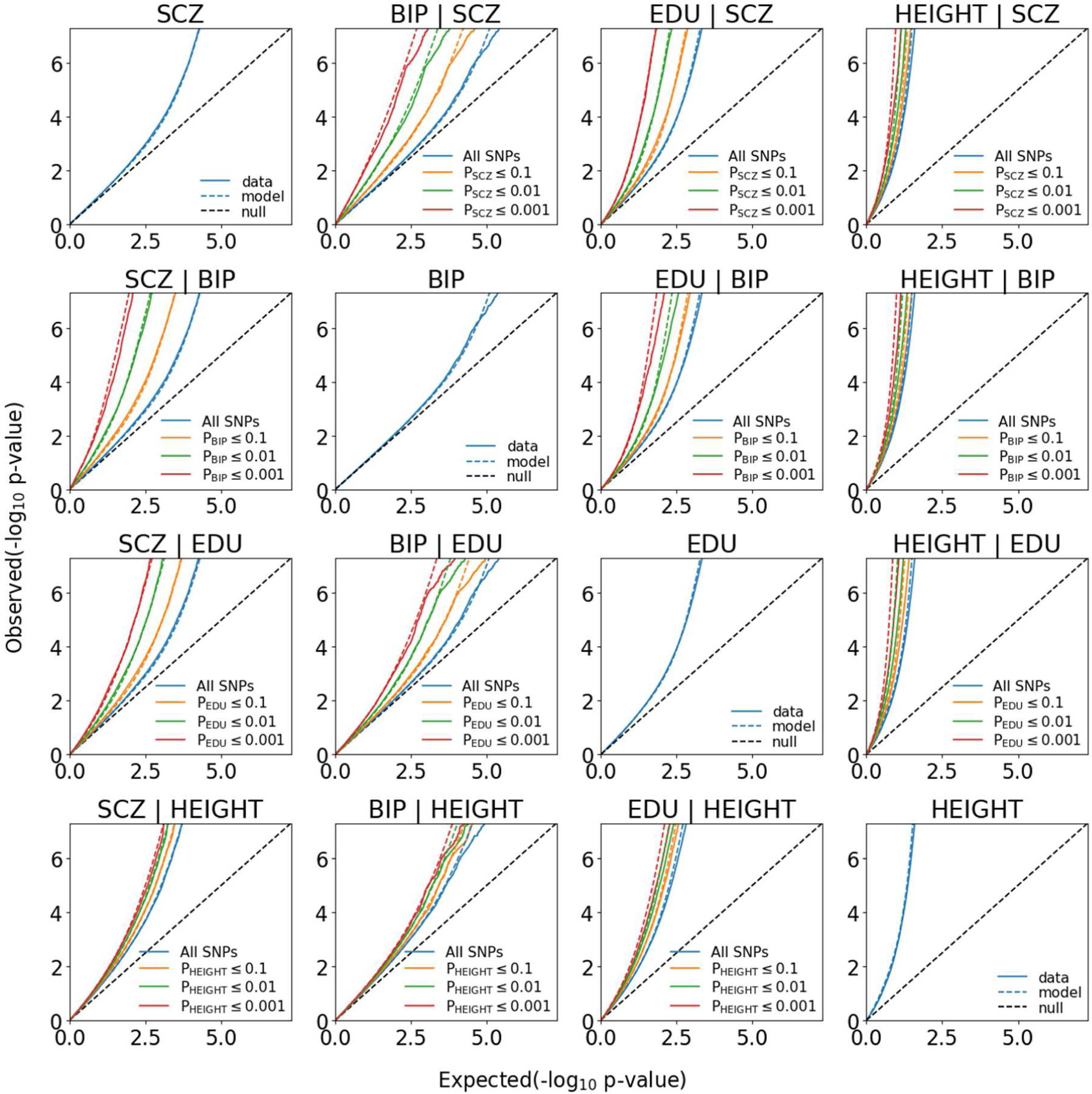
Conditional Q-Q plots of observed versus expected -log_10_ p-values in the primary trait as a function of significance of association with a secondary trait at the level of p≤0.1, p≤0.01, p≤0.001. Blue line indicates all SNPs. Dotted lines indicate model predictions for each stratum. Points on the QQ plot are weighted according to LD structure, using n=64 iterations of random pruning at LD threshold r2=0.1.

### Sensitivity analysis

For sensitivity analysis, we conducted simulations with traits that have a shared pattern of differential enrichment of heritability across genomic categories^19^, which is not accounted for by MiXeR model. Simulations were informed by the enrichment pattern of schizophrenia^20^, as estimated by stratified LD score regression^21^(Supplementary Figure 8a). In the univariate analysis, polygenicity was underestimated by about 20% (Supplementary Figure 9), indicating that the model may group adjacent causal variants together and interpret them as a single cluster. In the bivariate analysis we observe a small upwards bias in the estimate of polygenic overlap (Supplementary Table 3), but it did not exceed 10% of the polygenicity across all sufficiently powered scenarios.

Another assumption of MiXeR model is that effect sizes are independent of allele frequencies. We run simulations where, in addition to differential enrichment of genomic categories, all causal variants equally contribute to heritability regardless of their allele frequency (as modeled by stratified LDSR), as opposite to MiXeR assumption where causal variants contribute depending on their allele frequency. The results are showing that heritability is underestimated by 10% to 40% of its true value, while polygenicity is underestimated by a factor of 10 (Supplementary Figure 10), i.e. a larger bias than was observed in simulations with genomic annotations alone. The implications are less notable for the bivariate model: the estimated number of shared causal variants is consistent with the actual number arising by chance due to high polygenicity (Supplementary Table 4).

Finally, we run simulations with incomplete reference, and simulate phenotypes where causal variants are spread across our entire reference of N=11,015,833 variants, but only a fraction (50%, 25% or 12.5%) of the variants enter LD structure estimation and fit procedure. The results (Supplementary Table 5) show that the total number of causal SNPs, as well as the heritability, are estimated correctly, while polygenicity parameter is different from simulated value, because it reflects the fraction of all tagged causal variants with respect to the reference that went into LD structure estimation.

### GWAS Summary Statistics

We apply MiXeR to summary statistics from GWAS representing 7 phenotypes^11,20,22-26^(see Supplementary Table 6 for metadata about the studies).

MiXeR estimates of genetic correlation (Table 1, Supplementary Table 7) were generally consistent with those of cross-trait LD Score Regression^8^, with the highest genetic correlation observed between schizophrenia and bipolar disorder. Naturally, these disorders also exhibit substantial polygenic overlap, sharing 6.9K out of 9.5K causal variants involved. Here and below the numbers of causal variants are reported as 22.6% of their total estimate, which jointly accounts for 90% of SNP heritability in each phenotype, to avoid extrapolating model parameters into the area of infinitesimally small effects (Supplementary Figure 11).

Further, MiXeR reveals important differences among traits with low genetic correlation, represented as Venn diagrams of shared and unique polygenic components (Figure 3, Supplementary Figures 12, 13a-g). For example, schizophrenia and educational attainment exhibit substantial polygenic overlap, sharing 9.1K out of 12.1K of causal variants involved. On the contrary, schizophrenia and height share only about 1.1K out of 11.9K causal variants. Intriguingly, educational attainment and height also show low polygenic overlap, sharing 1.8K out of 14.0K causal variants. Nevertheless, these traits have relatively high correlation of effect sizes within the shared component, *p*_12_ = 0.44 (0.03), which at genome-wide level is observed as genetic correlation of rg=0.12 (0.01) according to MiXeR, or rg=0.14 (0.01) according to LDSR.

**Figure 3.**
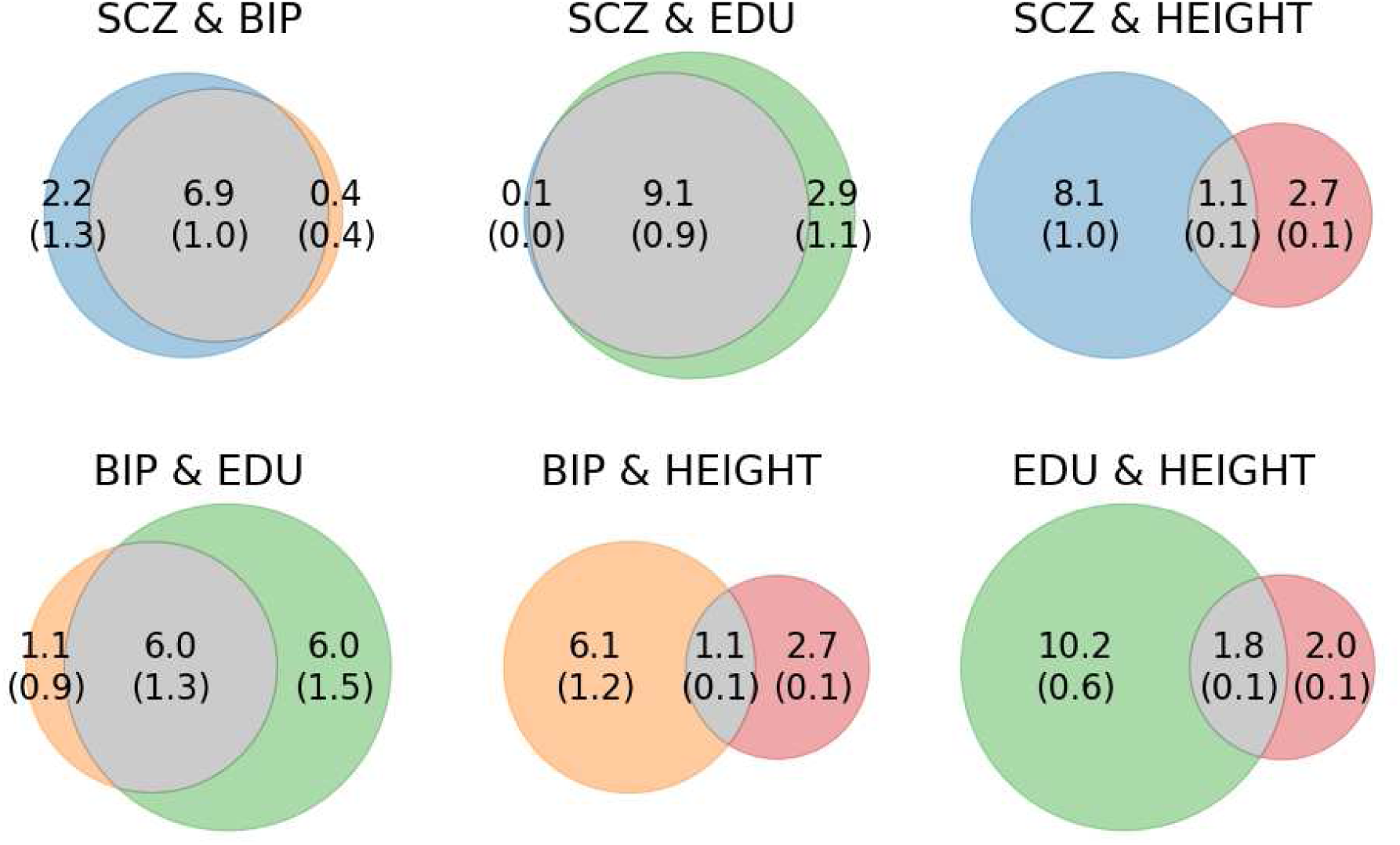
Venn diagrams of unique and shared polygenic component at the causal level, showing polygenic overlap (grey) between schizophrenia (SCZ, blue), bipolar disorder (BIP, orange), educational attainment (EDU, green) and height (red). The numbers indicate estimated quantity of causal variants (in 1,000) per component, explaining 90% of SNP heritability in each phenotype, followed by the standard error. The size of circles reflects the polygenicity.

MiXeR estimates of the unique polygenic components provide insight into the trait-specific genomic architecture. For example, while schizophrenia has 2.2K causal variants not shared with bipolar disorder, only 0.1K are not shared with educational attainment, and as many as 8.1K are non-overlapping with height. Also, for the other phenotypes the number of trait-specific causal variants varies across different pairs of traits (Figure 3).

Figure 4 and Supplementary Figures 14a-g visualize bivariate density of the observed GWAS signed test statistics (*z*_1j_, *z*_2j_), the predicted density 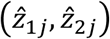 from MiXeR model, and estimated bivariate density of additive causal effects (*β*_1j_, *β*_2j_) that underlie model prediction. Figure 4 gives real examples for the three different scenarios of polygenic overlap (genetically independent traits, polygenic overlap with and without genetic correlation, as previously shown by Figure 1). Finally, we use conditional Q-Q plots^27,28^, where a consecutive deflection of the curves indicates polygenic overlap, and shows that MiXeR-based prediction provides accurate estimates of the data Q-Q plots (Figure 5).

## DISCUSSION

MiXeR is a novel method for cross-trait analysis of GWAS summary statistics, which enables a more complete quantification of polygenic overlap than provided by other existing tools^8,9,29,30^. In addition to genetic correlation, MiXeR estimates the total number of shared and trait-specific causal variants, providing new information into the genetic relationships between complex traits and disorders.

MiXeR extends cross-trait LD score regression^8^by incorporating a causal mixture model^15-18^, thus relying on a biologically more plausible prior distribution of genetic effect sizes compared to the “infinitesimal” model^31,32^. We show that polygenicity, measured as a total number of causal variants, and discoverability^33^, measured as variance of individual causal variants, has major implications on the future of GWAS discoveries (Supplementary Figure 15).

Applying MiXeR to real phenotype data, we provide new insights into the genetic relationships between schizophrenia, bipolar disorder, educational attainment and height. In line with the strong clinical relationship^34^between schizophrenia and bipolar disorder, and prior genetic studies^25,35^, we find substantial polygenic overlap between these two disorders. Intriguingly, both schizophrenia and bipolar disorder are estimated to have a small fraction of causal variants conferring individual risk of a specific disorder (Figure 3). Identifying such genetic variants could provide critical knowledge about the distinct genetic risk architectures underlying these psychiatric disorders. Moreover, we find that nearly all causal variants influencing schizophrenia risk also appear to influence educational attainment, despite a genetic correlation close to zero (Table 1). This is in line with recent studies demonstrating shared genetic loci between schizophrenia and educational attainment^14^and a strong genetic dependence between the phenotypes possibly related to different subtypes of schizophrenia^13^. In contrast, 85% of genetic variants influencing bipolar disorder also appear to influence educational attainment, but there is a significant positive genome-wide correlation of 0.20(0.03) in compliance with the cross-trait LD score regression estimate of 0.18(0.03) (Table 1, Supplementary Table 7).

We show that polygenicity is best expressed as a total number of causal variants (Supplementary Table 5). Previous studies presented it as a fraction, which is highly dependent on the used reference (1.1M hapmap in ref^17^, or 484K Affymetrix SNPs in ref^18^). When expressed as a total number, or estimates of polygenicity for schizophrenia, bipolar disorder, educational attainment and height are consistent with previously reported results. In addition, we estimate that just 5% of causal variants are needed to explain 50% of heritability, and 22.6% of causal variants are needed to explain 90% of heritability (Supplementary Figure 11). These numbers are expected to be less dependent on modeling assumptions, because with finite GWAS sample it is not possible to distinguish small effects from truly null effects. The actual number of causal variants is, potentially, even higher, as our model tends to clump together variants if they are located too close to each other (Supplementary Tables 3, 4).

Some existing methods can already uncover polygenic overlap in the absence of genetic correlation. For example, conjFDR analysis^27,28^is a non-parametric model-free approach, which detects shared genetic loci regardless of their allelic effect directions, by prioritizing variants with strong associations across more than one GWAS^36^. Other methods, including gwas-pw^37^and HESS^29^, also aim at detecting genomic loci jointly associated with two traits. MiXeR complements these methods by providing an easily interpretable high-level overview of the shared and unique genomic architectures underlying complex phenotypes.

MiXeR has some notable advantages compared to the existing methods that implement causal mixture. First, our mathematical model for the likelihood term p(*z*_j_l*β*_j_) is conceptually simpler and more flexible, resulting in unbiased estimates of model parameters across a wide range of simulation scenarios (Supplementary Figure 1-3) and providing accurate prediction of GWAS z-scores across varying ranges of MAF and LD (Supplementary Figure 6a,b). Second, MiXeR implementation works well with a large reference of 10M variants, while other methods have reduced it to 1.1M HapMap SNPs (ref^17^) or 484K Affymetrix SNPs (ref ^18^). Finally, our model individually processes all SNPs, without grouping them into bins (ref ^15^).

MiXeR models causal effects as a single gaussian component, while recent work^17,38^suggests that certain phenotypes, including height, require at least two causal components of small and large effects. We note that MiXeR model still provides good fit for SNPs not reaching GWAS threshold (Supplementary Figure 16) and shows deviations only towards the tail of the distribution. To further investigate the effects of model misspecification we implemented right-censoring of genome-wide significant SNPs (see Online methods). Results (Supplementary Tables 7, 8) are consistent with our main analysis, except for height which received a lower estimate of heritability (65% instead of 70%), a slight increase in polygenicity, and increased polygenic overlap with other traits. We advocate that for a better estimate of height’s polygenicity it would be beneficial to run MiXeR on a residualized GWAS, after covarying association statistics for genotypes of all genome-wide significant SNPs.

Recent work suggests the importance of MAF- and LD-dependent genetic architectures^18,39^, which are not directly modeled by MiXeR. Our simulations with an extreme case of a different MAF model shows 10-40% underestimation of heritability (Supplementary Figure 10), but less noticeable effect on the relative proportion of shared causal variants (Supplementary Table 4). On real data we observe effects of MAF-dependent architectures by drawing Q-Q plots for subsets of SNPs (Supplementary Figures 17a-g) partitioned into 9 groups according to minor allele frequency (MAF) and LD score, where the model tends to underestimate z-scores in low MAF bins. This effect, however, is quite subtle, and does not manifest itself on the overall Q-Q plots (Supplementary Figure 16).

The MiXeR method requires large GWAS studies. Our recommendation is to apply MiXeR to studies with at least N=50 000 participants, and inspect standard errors reported by MiXeR. Polygenicity estimation requires more GWAS power than heritability estimation, which can be visually explained by GWAS Q-Q plots (Supplementary Figure 16): heritability is determined by the overall departure of the GWAS curve from the null line, while polygenicity is determined by its curvature, i.e. the point where the GWAS curve begins to bend upwards from the null line, which is harder to estimate when GWAS signal is weak. This is captured by MiXeR standard errors, which show that individual parameters of the mixture model have lower estimation accuracy than their combinations – for example, relative errors for *π*_1_ and 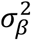 are larger than for the heritability estimate 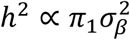, due to inversely-correlated errors (Supplementary Table 2). Despite these limitations, there is still a clear minimum on the energy landscape of cost function (Supplementary Figures 18, 19, showing log-likelihood as a function of model parameters around the optimum).

In our future work we are planning to incorporate an additional gaussian component to model small and large effects^17^, and explicitly account for MAF-dependent architectures^39^. Further extensions may account for differential enrichment for true associations across genomic annotations^19^. Another limitation to address is that MiXeR model assumes similar LD structure among studies and is not currently applicable for analysis across different ethnicities. We aim to extend the MiXeR modelling framework to be used to improve power for SNP discovery by estimating the posterior effect size of SNPs associated with one trait given the test statistics in another trait, as well as for improving predictive power of polygenic risk scores.

In conclusion, MiXeR represents a useful addition to the tool-box for cross-trait GWAS analysis. By taking into account the intricate polygenic architectures of complex phenotypes MiXeR allows for measures of polygenic overlap beyond genetic correlation. We expect this to lead to new insights into the pleiotropic nature of human genetic etiology.

## Supporting information

## ACKNOWLEDGMENTS

This work was supported by the Research Council of Norway (#223273, #225989, #248778) South-East Norway Health Authority (#2016-064, #2017-004), KG Jebsen Stiftelsen (#SKGJ-Med-008), and National Institutes of Health (R01MH100351, R01GM104400). The simulations were performed on resources provided by UNINETT Sigma2 - the National Infrastructure for High Performance Computing and Data Storage in Norway.

## AUTHOR CONTRIBUTIONS

Conceived and designed the study – AMD, OAA, OF, Method development – AMD, OF, DH, AAS, Analysis and interpretation of results – OF, DH, AAS, OBS, CCF, YW, AW, SD, Drafting manuscript – OF, OAA, OBS, Revision and approval of final manuscript – ALL AUTHORS

## URLS

https://github.com/precimed/ - MiXeR code (MATLAB/Octave), will be released after publication; https://data.broadinstitute.org/alkesgroup/LDSCORE/ - LD scores and reference panel derived from 1000 Genomes phase 3.

## CORRESPONDING AUTHORS

Oleksandr Frei Anders M. Dale

## ONLINE METHODS

This article is accompanied by a Supplementary Note with further details.

### Bivariate causal mixture model

Consider simple additive model of genetic effects, ignoring gene-environment interactions, epistasis and dominance effects. Under these assumptions, the contribution of the genotype to the phenotype is modelled as a sum of individual contributions from genetic variants: *y*_k_ = ∑_j_ *g*_jk_ *β*_j_, where *y*_k_ is a quantitative phenotype or disease liability of k-th individual, *g*_jk_ is 0,1,2-coded number of reference alleles for j-th variant, and *β*_j_ is additive genetic effect of allele substitution. We say that genetic variant is causal for a trait if it has non-zero effect on that trait (*β*_j_ ≠ 0).

MiXeR builds upon univariate causal mixture model^15^, 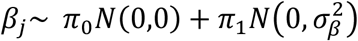, which assumes that only a small fraction (*π*_1_) of variants has an effect on the trait, while the effect of the remaining variants is zero. In a joint analysis of two traits we expect some variants to affect both traits; some variants to affect one trait but not the other; and most variants to have no effect on either trait. Based on these assumptions, MiXeR models additive genetic effects *β*_1*j*_, *β*_2*j*_ of variant *j* on the two traits as a mixture of four bivariate Gaussian components (Figure 1):

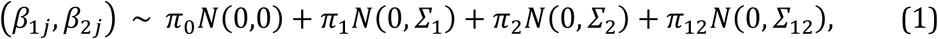

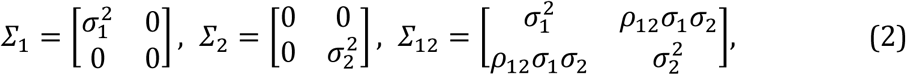

where *π*_1_ and *π*_2_ are weights of the unique components (variants with an effect on the first only, and on the second trait only); *π*_12_ is a weighting of the component affecting both traits; and *π*_0_ is a fraction of variants that are non-causal for both traits, *π*_0_ + *π*_1_ + *π*_2_ + *π*_12_ = 1; 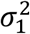 and 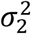 control expected magnitudes of per-variant effect sizes; and *?*_12_ is the correlation coefficient of the effect sizes in the shared component. All parameters are assumed to be the same for all genetic variants.

The effects 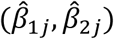 estimated by a GWAS, represent only proxies of the true causal effects (*β*_1j_, *β*_2j_), which are distorted by limited sample size (poor statistical power), cryptic relatedness within a GWAS sample as well as LD between variants. To disentangle these effects we derive the likelihood term for observed GWAS signed test statistics (*z*_1j_, *z*_2j_), incorporating effects of LD structure (allelic correlation *r*_*ij*_ between variants *i* and *j*); heterozygosity *H*_j_ = 2*p*_j_(1 - *p*_j_) where *p*_*j*_ is the minor allele frequency of the *j*-th variant; number of subjects genotyped per variant (*N*_1*j*_ and *N*_2*j*_); and variance distortion parameters 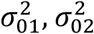 and *ρ*0. Specifically (see Supplementary Note),

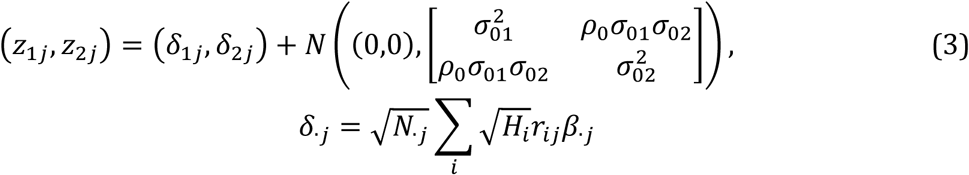

The nine parameters of the model 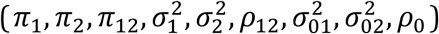 are fit by direct optimization of the weighted log likelihood, with standard errors estimated from the Observed Fishers Information matrix.

Forcing *π*_12_ = 1 (so that *π*_0_ = *π*_1_ = *π*_2_ = 0) reduces our model to an “infinitesimal” assumption that underlie cross-trait LD score regression^8^. Under this constraint our model predicts that GWAS signed test statistics follow bivariate Gaussian distribution with zero mean and variance-covariance matrix

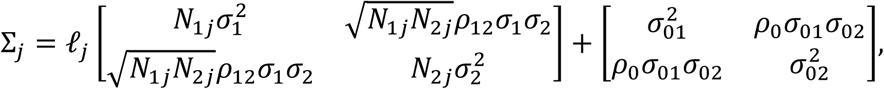

i.e., (*z*_1*j*_,*z*_2*j*_) ∼ N(0,Σ*j*_*j*_), where 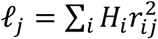 is the LD score (adjusted for heterozygosity). This model is consistent with cross-trait LD score regression, with expected chi square statistics 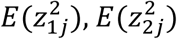 and cross-trait covariance E(*z*_1j_*z*_2j_) being proportional to the LD score of j-th SNP, and parameters *p*_0_, σ_01_, σ_02_ playing the role of LD score regression intercepts^40^. The only distinction here is that we choose to model effect sizes that are independent of allele frequency, leading to the incorporation of *H*_*i*_ in our model; this factor is absent from the LD score regression model due to the assumption there of effect sizes that are inversely proportional to *H*_*i*_. Thus, MiXeR is a direct extension to cross-trait LD score regression, which relaxes the “infinitesimal” assumption.

### Model for bivariate distribution of GWAS z-scores

We derive two models for GWAS z-scores, which we call “fast model” and “full model”. “fast model” is quicker to run, and we use it to perform initial search in the space of model’s parameters. “full” model is slower but more accurate, and we use it for a final tuning of model estimates.

“full” model for GWAS z-scores approximates (*z*_1j_, *z*_2j_) distribution of a given GWAS SNP as a mixture of K=20000 bivariate normal distributions, all having equal weight in the mixture. For each k = 1,…, K we randomly draw the location of causal variants (*π*_1_N causal variants specific to the first trait, *π*_2_N specific to the second trait, and *π*_12_N shared causal variants, where N denotes the total number of variants in the reference panel), and calculate the variance-covariance matrix 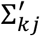 from (3), using estimated LD r^2^correlations between the assumed causal variants and the GWAS SNP. Then

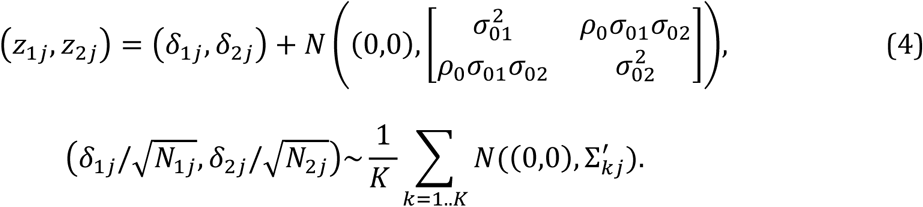

The “fast” model is derived from the method of moments (see Supplementary note):

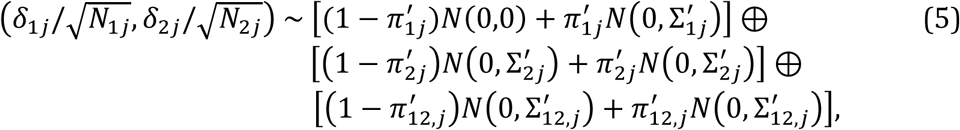

where ⊕ denotes convolution of probabilistic distribution functions (so that right-hand size evaluates to a mixture of 8 components), 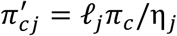 is adjusted weight of mixture component 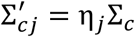 is adjusted variance-covariance matrix; 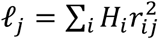 is the LD score, adjusted for heterozygosity^41^; and 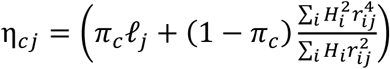 can be interpreted as shape parameter that affects fourth and higher moments of the distribution. This model explains second moments 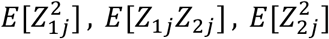, and fourth moments 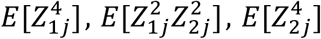, of z score distribution, and forms a theoretical basis for the mixture model of sparse and ubiquitous effects^42,43^. Of interest is that the “fast” model involves the forth power of allelic correlation 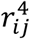, which is directly proportional to kurtosis (measure of heavy tails) of z-score distribution.

### LD structure estimation

To estimate LD structure we use 489 individuals of 1000 Genome project^44^(phase 3 data), obtained from LD score regression website^8,21,45^(see URLs). 14 individuals were excluded due to relatedness^46^. For simulations LD scores were estimated from the genotypes that we use to produce synthetic GWAS data. LD r^2^coefficients were calculated using PLINK^47^with LD r2 cutoff of 0.05 and fixed window size of 50 000 SNPs, corresponding on average to a window of 16 centimorgans. We deliberately choose a larger LD window compared to LDSR-recommended window of 1 centimorgan because the later appears to truncate a noticeable part of LD structure. At the same time, we did not observe an effect of using unbiased estimate^48^of r^2^, thus fall back to the standard Pearson correlation coefficient. In simulations LD structure was reestimated from the actual genotypes used to generate synthetic summary statistics. We employ small integer compression^49^for efficient storage of the LD matrix.

### Fit procedure

We fit the model by direct optimization of weighted log likelihood

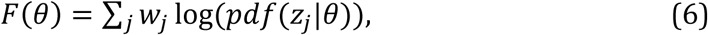

where 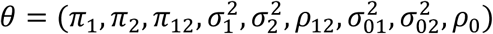 is a vector of all parameters being optimized, and weights *w*_j_ chosen by random pruning (64 iterations at LD r2 0.1). Optimization is done by Nelder-Mead Simplex Method^50^as implemented in MATLAB’s fminsearch. First, we fit univariate parameters separately for each trait, i.e. 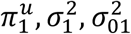 for the first trait, and similarly for the second trait. Univariate fit employs a sequence of optimizations to ensure robust convergence: first, we use “fast” model under constraint 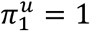 to find 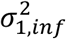 and to initialize 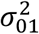 second, we use constraint 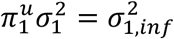 to find initial values of 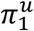 and 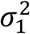, again with “fast” model. Finally, we use full model and unconstrained optimization to jointly fit 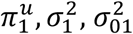 parameters. The same procedure is repeated for the second trait, to find 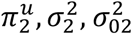. To improve convergence, we parametrize univariate log-likelihood as a function of 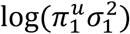 and 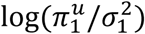, which represent almost independent dimensions of the energy landscape. In bivariate optimization use “fast” model and constraint *π*_12_ = 1 to initialize *p*_12_ and *p*_0_. Then proceed with “full” model optimization of the parameters specific to bivariate model (*π*_12_, *p*_12_, *p*_0_), constraining all other parameters to their univariate estimates. The additional analysis (Supplementary Tables 7, 8) uses right-censoring^51^of z-scores exceeding *z*_t_ = 5.45, by using cumulative distribution function^52^in the log likelihood:

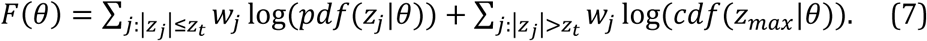

### Standard error estimation

We estimate standard errors of all parameters from the observed Fisher’s information, based on the “fast” model. It is known from the likelihood optimization theory that the observed Fisher’s information may not be suitable for a parameter near its boundary, which is applicable to the mixture weights *π*_1_, *π*_2_, *π*_12_ and the correlation of effect sizes *p*_12_. To mitigate this problem we apply transformations **—** MATLAB’s logit() for *π*_1_, *π*_2_, *π*_12_, exp() for 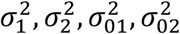, and erf()for *p*_0_, *p*_12_ and estimated variance-covariance matrix of errors in the transformed parameter space. We validated that our estimates based on the observed Fisher’s information are in good agreement with block jack-knife estimates. To estimate standard errors for a function of the parameters, such as *r*_g_ or h^2^, we incorporate linear correlation among parameter errors in the transformed space. We sample N=1000 realizations of the parameter vector, calculating the function (e.g., *r*_g_ or h^2^) on each of them, and report the standard deviations. In cases when joint hessian was not positive definite, we estimate marginal errors of fitted parameters.

### Large LD blocks

The log-likelihood cost function and the Q-Q plots apply a weighting scheme to SNPs to avoid overcounting evidence from large LD blocks. As an alternative to weighting by inverse LD score, we choose to infer the weights by random pruning. This technique is a stochastic procedure which averages log likelihood function across repeatedly selected subsets of variants, such that for each pair of variants i, j in a subset J the squared allelic correlation 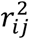 falls below certain threshold. Given T iterations of random pruning the log-likelihood function can be calculated as follows:

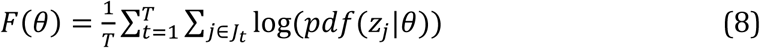

which is equivalent to weighted log-likelihood F(0) = ∑_j_ *w*_j_ log(pdf(*z*_j_|0)) with weights *w*_j_ = |{t: j ∈ *J*_t_}|/T, |S| denotes cardinality of set S. Random pruning with stringent threshold r2=0.1 justify independent modeling of the residuals in (3) across SNPs, which otherwise would be correlated.

### Heritability estimates

In additive model, SNP heritability is defined as a sum across all causal variants: 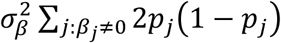, which we approximate from an average heterozygosity of all variants in the reference: 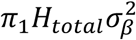, where 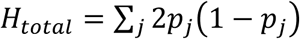. To estimate the proportion of causal variants that explain certain fraction of heritability (Supplementary Figure 11) we randomly sample N=10000 causal effects from the reference, draw their effects *β*_j_ from normal distribution, sort according to 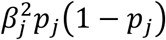, and report fraction of variants that cumulatively account 90% of heritability.

### Genetic correlation

Parameter *p*_12_ in MiXeR defines correlation of effect sizes within shared polygenic component. Genome-wide genetic correlation, calculated across all SNPs, is related to *p*_12_ by the following formula that involves polygenicity 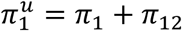 and 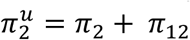 of the traits, and polygenic overlap *π*_12_:

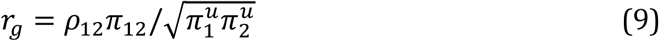

For traits with K-fold difference in polygenicity 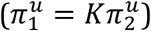 the formula predicts an upper bound on genome-wide genetic correlation: 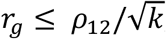, where equality holds if causal variants of the less polygenic trait form a subset of the higher-polygenic trait.

### Quantile-quantile plots

Univariate Q-Q plots and stratified Q-Q plots for the model were constructed from pdf_j_(z) density as defined by (3), given fitted parameters of the model and LD structure of j-th SNP, calculated across a fine grid of z-scores ranging from 0 to 38 with 0.05 step. We average pdf_j_(z) across 1% of randomly sampled SNPs, and numerically integrate the resulting probability density function to convert it into cumulated distribution function. Error bars on data Q-Q plots represent the 95% binomial confidence interval 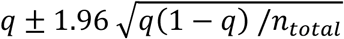, where q is portability of observing a p-value as extreme as, or more extreme then the chosen p-value, and *n*_total_ is the effective number of SNPs after controlling for LD structure, which in our case was calculated as a sum of random pruning weights across all SNPs.

### GWAS power curves

Causal mixture model can project the future of GWAS discoveries, by estimating proportion S(N) of narrow-sense heritability captured by genome-wide significant SNPs at a given sample size N. The S(N) is defined as follows:

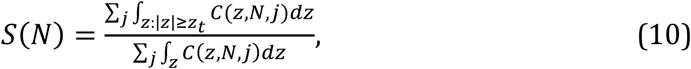

where *z*_t_ = 5.45 gives z-score corresponding to the standard genome-wide significance threshold 5·

### SNPs in the analysis

To enable direct comparison of our model with LD score regression we use the same set of SNPs in our log likelihood optimization, which consist of approx. 1.1 million variants, subset of 1000 Genomes and HapMap3^53^, with MAF above 0.05, ambiguous SNPs excluded, imputation INFO above 0.9, MHC and other long-range LD regions excluded. Calculation of the LD structure, LD scores *l*_j_ and shape parameter η_j_ are based on the approx. 10 million SNPs from 1000 Genomes Phase 3 data, downloaded from LD score regression website (see URLs). In simulations we generate GWAS and estimate LD structure on a subset of 11,015,833 SNPs from 1000 Genomes Phase 3, with MAF above 0.002, call rate above 90%, excluding duplicated RS numbers; the fit procedure was constrained to approximately 130K GWAS SNPs, keeping only HapMap3 SNPs, and pruning SNPs at LD r2 threshold of 0.1.

### LD score regression estimates

For dichotomous phenotypes we used an effective sample size of *N*_eff_ = 4/(1/*n*_case_ + 1/*n*_cont_) to account for imbalanced number of cases and controls, both in MiXeR and in LD score regression. Additionally, we run LDSR using MiXeR MAF model (using –per-allele flags in LD score estimation), and show the results alongside with original LDSR estimates (Supplementary Tables 7, 8). For case/control phenotypes heritability is reported on the observed scale.

### Simulations

In our simulations we use a panel of *N* = 100,000 samples, generated by HapGen2^54^using 1000 Genomes^44^data to approximate the LD structure for European ancestry. For each simulation run we use PLINK to obtain GWAS summary statistics of two synthesized quantitative phenotypes, with complete sample overlap between GWAS samples. Quantitative phenotype *y*_k_ of k-th sample is calculated via simple additive genetic model, *y*_k_ = ∑_j_ *g*_kj_*β*_j_ + *E*_k_, where *g*_kj_ is the number of reference alleles for j-th SNP on k-th sample, *β*_j_ is causal effect size, and E is the residual vector drawn from normal distribution with zero mean and variance chosen in a way that sets heritability h^2^= var(Gβ)/var(y) to a predefined level.

For the simulations shown in Figure *2*, we draw effect sizes (*β*_1j_, *β*_2j_) from the four-component mixture model (1), varying polygenicity of each phenotype (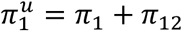 and 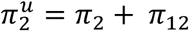), and polygenic overlap (*π*_12_). We choose total polygenicity in both traits to be 3×10^−3^or 3×10^−4^and include an additional scenario of uneven polygenicity 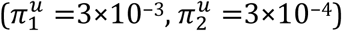. For each combination 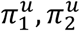 we set polygenic overlap to be a fraction of total polygenicity 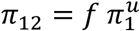, choosing the fraction f from six equally spaced values (0.0 to 1.0 with a step of 0.2). Correlation of effect sizes *p*_12_ set to 0.0 or 0.5. Heritability was set to 0.1, 0.4 or 0.7, which let us keep GWAS sample size constant (N=100,000) because the distribution of GWAS z-scores depends on N and *h*_2_ only through their product, *h*_2_ × N (thus, simulations with N=700,000 and *h*_2_ = 0.1 would be equivalent to our scenario with N=100,000 and *h*_2_=0.7). Finally, for each combination of heritability, polygenicity, polygenic overlap and correlation of effect sizes, we repeat simulations 10 times.

For the simulations with differential enrichment, we simulate three levels of polygenicity (3K, 30K and 300K causal variants), three levels of heritability (0.1, 0.4 and 0.7), and for each combination generate 20 pairs of genetically independent traits (except for having shared pattern of enrichment). To simulate the enrichment, we keep a constant variance of effect sizes across all SNPs but modulate the probability of having causal variant proportionally to LDSR regression coefficient. We cover two scenarios: first, LDSR with MiXeR MAF model (using –per-allele flags in LD score estimation), and secondly with the original LDSR MAF model.

## SUPPLEMENTARY NOTE

Separate PDF document.

## SUPPLEMENTARY FIGURES

1. Simulations: polygenic overlap estimates in bivariate analysis
2. Simulations: estimates of the correlation of effect sizes in bivariate analysis
3. Simulations: estimates of genetic correlation in bivariate analysis
4. Simulations: polygenicity and heritability estimates in univariate analysis
5. Simulations: Q-Q plots with simulated data
6. Simulations: Q-Q plots of SNPs partitioned into a grid of MAF and LD score
7. Simulations: stratified QQ plots for traits with and without polygenic overlap
8. Sensitivity analysis: differential enrichment profile of synthesized data
9. Sensitivity analysis: polygenicity and heritability estimates under differential genomic enrichment
10. Sensitivity analysis: polygenicity and heritability estimates under differential genomic enrichment and mis-specified MAF architecture
11. Cumulated fraction of explained heritability versus proportion of causal variants
12. Venn diagrams across all traits.
13a-f. Venn diagrams and conditional cross-trait QQ plots
14a-f. Observed and predicted bivariate density of GWAS association statistics
15. Projected power plot for future GWAS sample sizes
16. Univariate Q-Q plots
17a-f. Q-Q plots of SNPs partitioned by MAF and LD score
18. Univariate likelihood as a function of polygenicity parameter
19. Bivariate likelihood as a function of polygenic overlap parameter

## SUPPLEMENTARY TABLES

1. Simulations: parameter estimates in bivariate analysis
2. Simulations: polygenicity and heritability estimates in univariate analysis
3. Sensitivity analysis: polygenic overlap estimates under differential genomic enrichment
4. Sensitivity analysis: polygenic overlap estimates under genomic enrichment and misspecified MAF architecture
5. Sensitivity analysis: simulations with incomplete reference
6. Summary statistics Metadata
7. Results of bivariate analysis with MiXeR, with and without right-censoring
8. Results of univariate analysis with MiXeR, with and without right-censoring

## EQUATIONS

1. MiXeR (beta1, beta) prior distribution
2. MiXeR Sigma1, Sigma2, Sigma12
3. MiXeR (z1, z2) exact formula
4. MiXeR (z1, z2) full model (sampling)
5. MiXeR (z1, z2) fast model (two gaussian)
6. MiXeR log likelihood function
7. Right censoring
8. MiXeR random pruning
9. MiXeR genetic correlation
10. MiXeR power curves S(N)

