## Supplementary material for "Bivariate causal mixture model quantifies polygenic overlap between complex traits beyond genetic correlation"

#### 1a. Simulations: polygenic overlap estimates in bivariate analysis, without genetic correlation

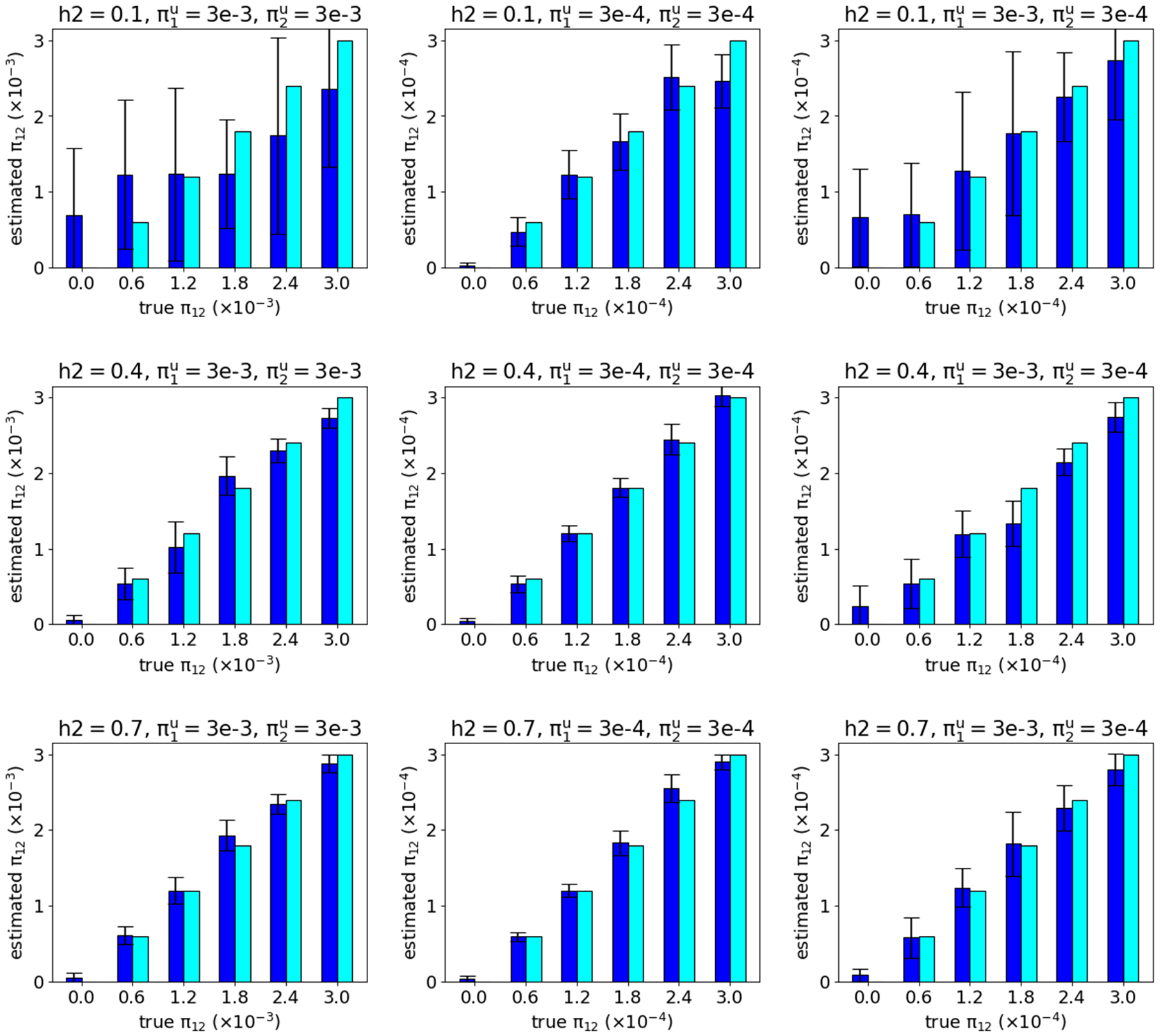


Simulation with synthetic GWAS summary statistics, generated according to MiXeR model, showing accuracy of $\pi_{12}$ parameter in scenarios without genetic correlation (simulated rho12=0.0). The bars in blue indicate an average value of model estimates across 10 simulation runs. The bars in cyan show true (simulated) parameters. Error bars represent standard deviation of the model estimate across 10 simulation runs.

#### 1b. Simulations: polygenic overlap estimates in bivariate analysis, with genetic correlation

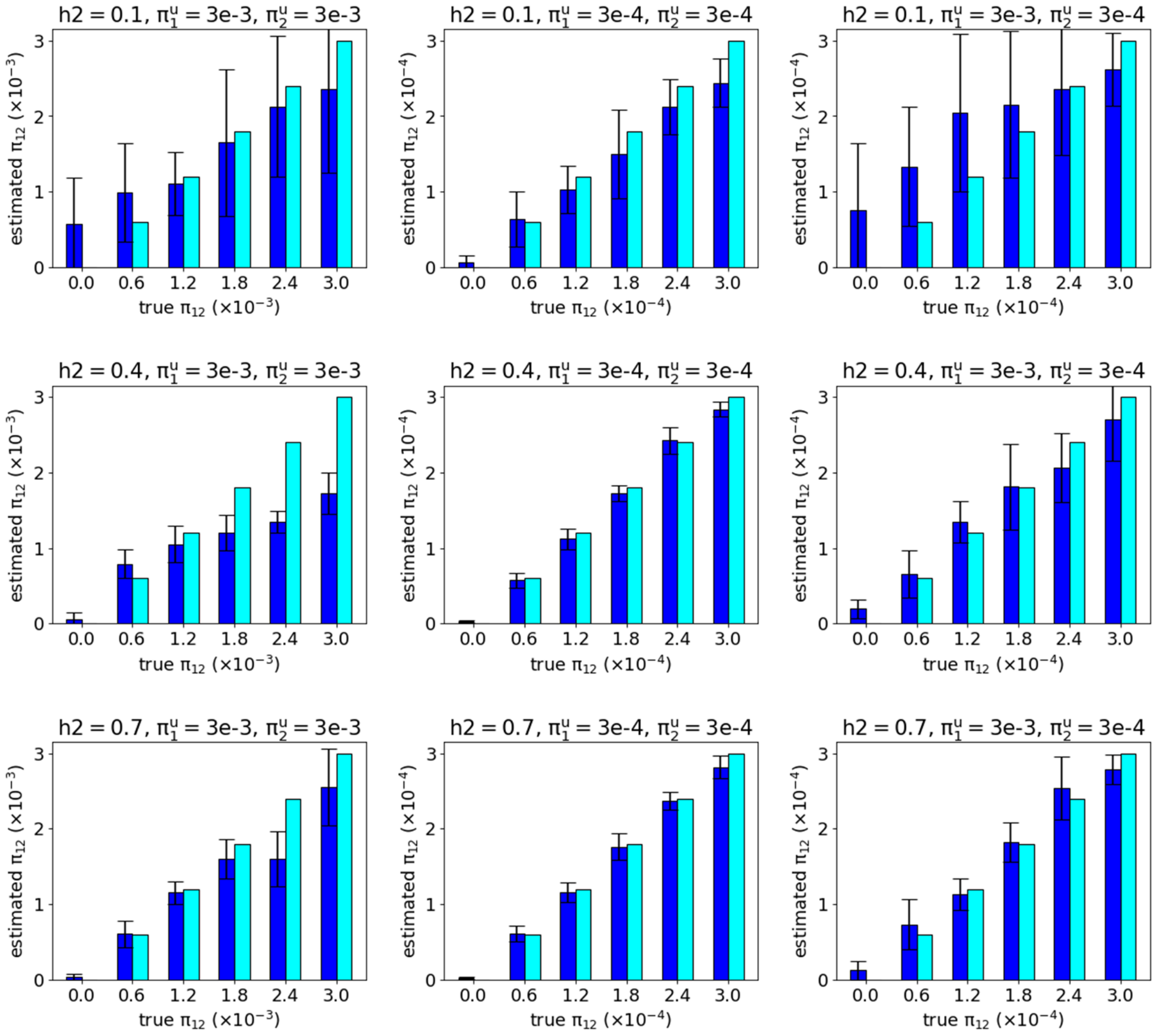


Simulation with synthetic GWAS summary statistics, generated according to MiXeR model, showing accuracy of $\pi_{12}$ parameter in scenarios with genetic correlation (simulated rho12=0.5). Appearance of the data bars and error bars is the same as on the previous figure.

#### 2a. Simulations: estimates of the correlation of effect sizes in bivariate analysis, simulated rho12=0

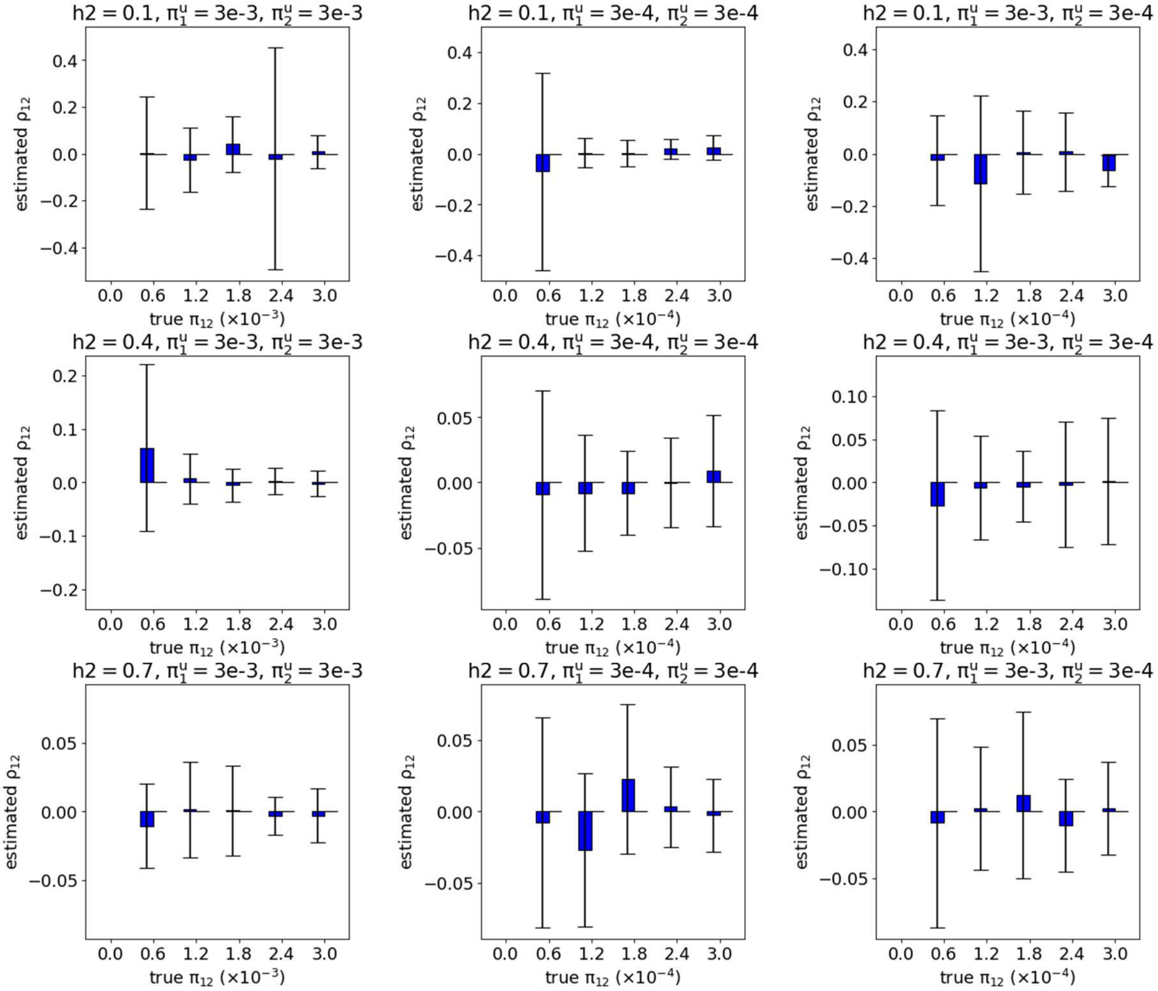


Simulation with synthetic GWAS summary statistics, generated according to MiXeR model, showing accuracy of rho12 parameter in scenarios without genetic correlation (simulated rho12=0). Appearance of the data bars and error bars is the same as on the previous figure.

#### 2b. Simulations: estimates of the correlation of effect sizes in bivariate analysis, simulated rho12=0.5

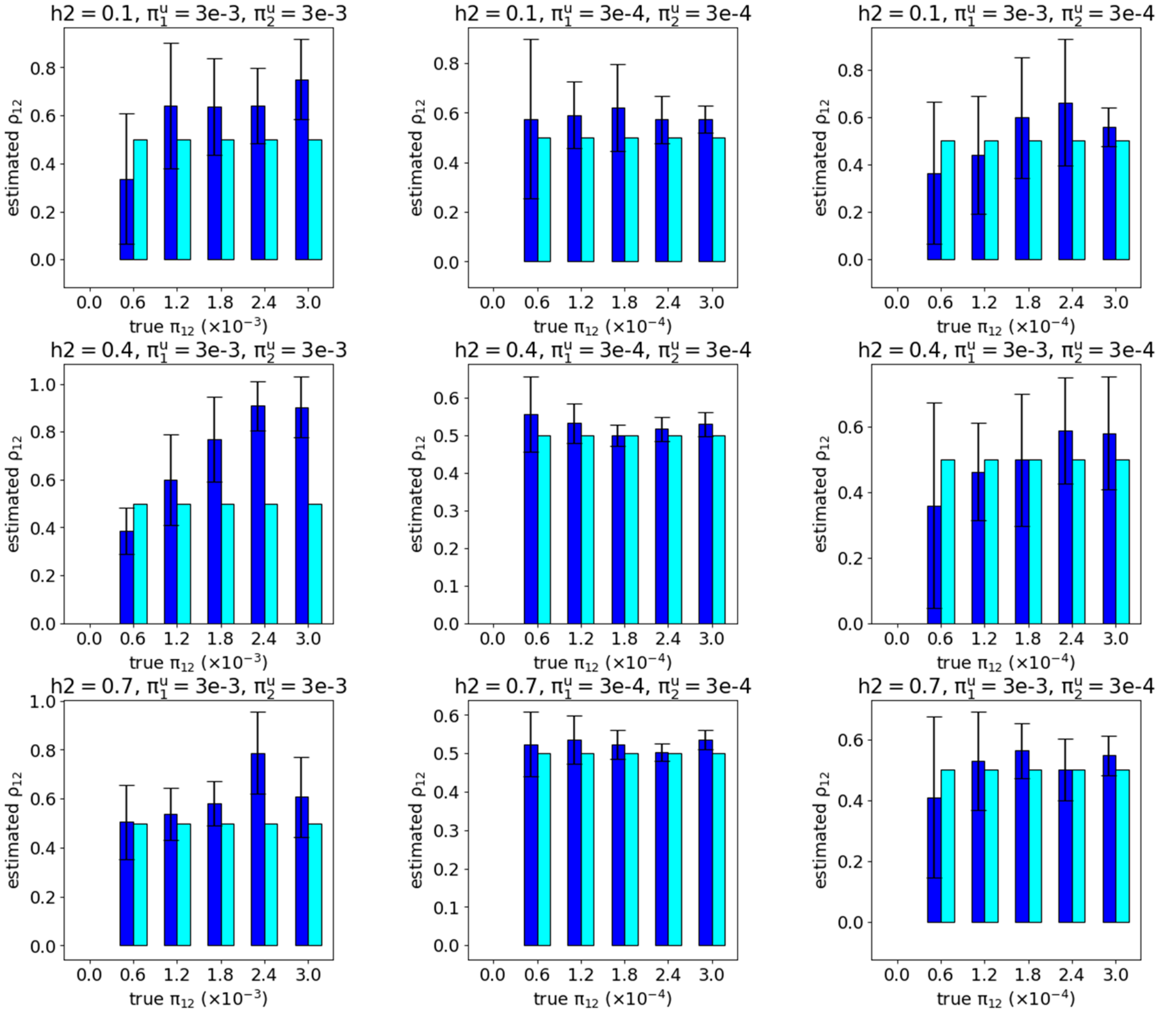


Simulation with synthetic GWAS summary statistics, generated according to MiXeR model, showing accuracy of rho12 parameter in scenarios with genetic correlation (simulated rho12=0.5). Appearance of the data bars and error bars is the same as on the previous figure.

#### 3a. Simulations: estimates of genetic correlation in bivariate analysis

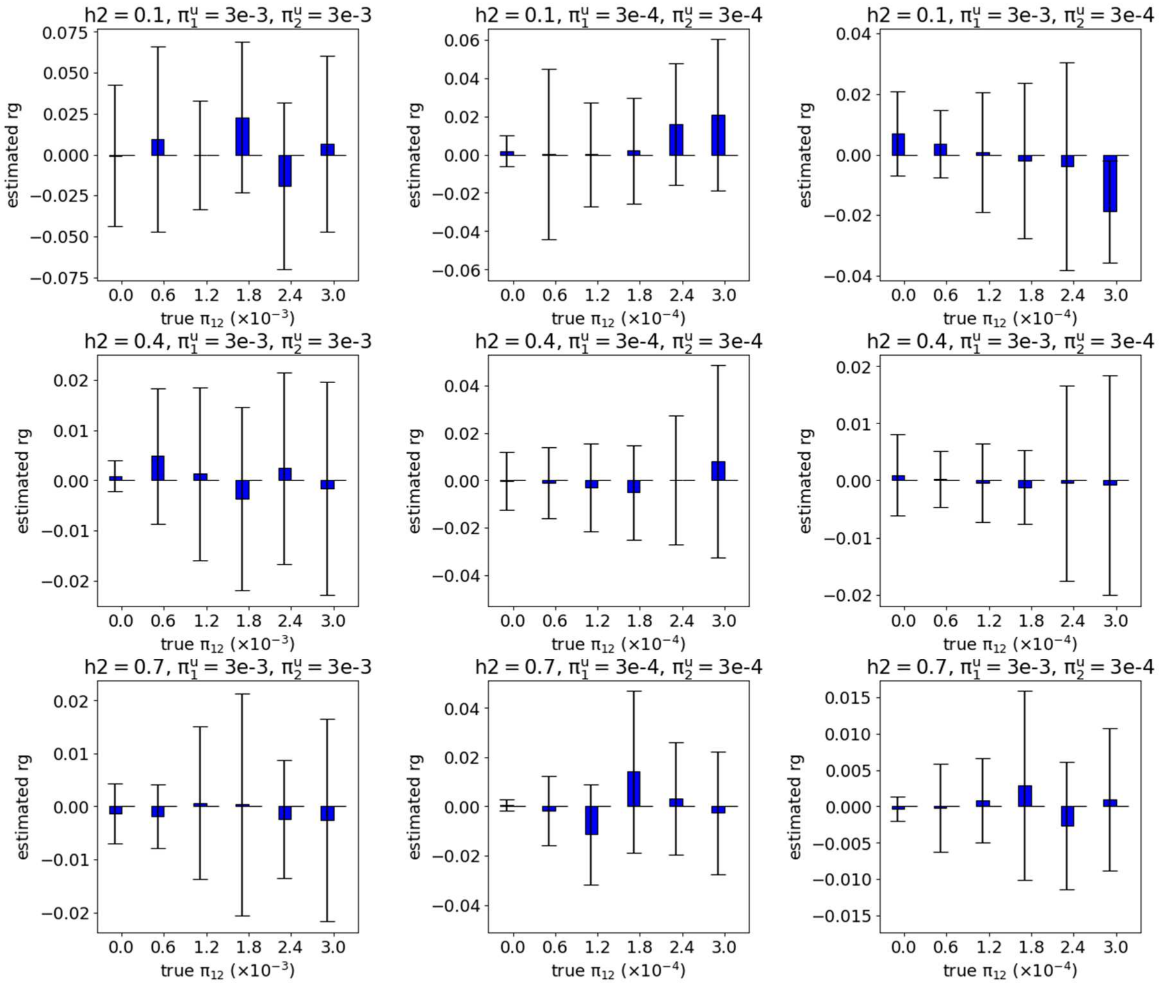


Simulation with synthetic GWAS summary statistics, generated according to MiXeR model, showing accuracy of $r_g = \rho_{12}\pi_{12}/\sqrt{\pi_1^u \pi_2^u}$ parameter in scenarios without genetic correlation (simulated rho12=0).

#### 3b. Simulations: estimates of genetic correlation in bivariate analysis

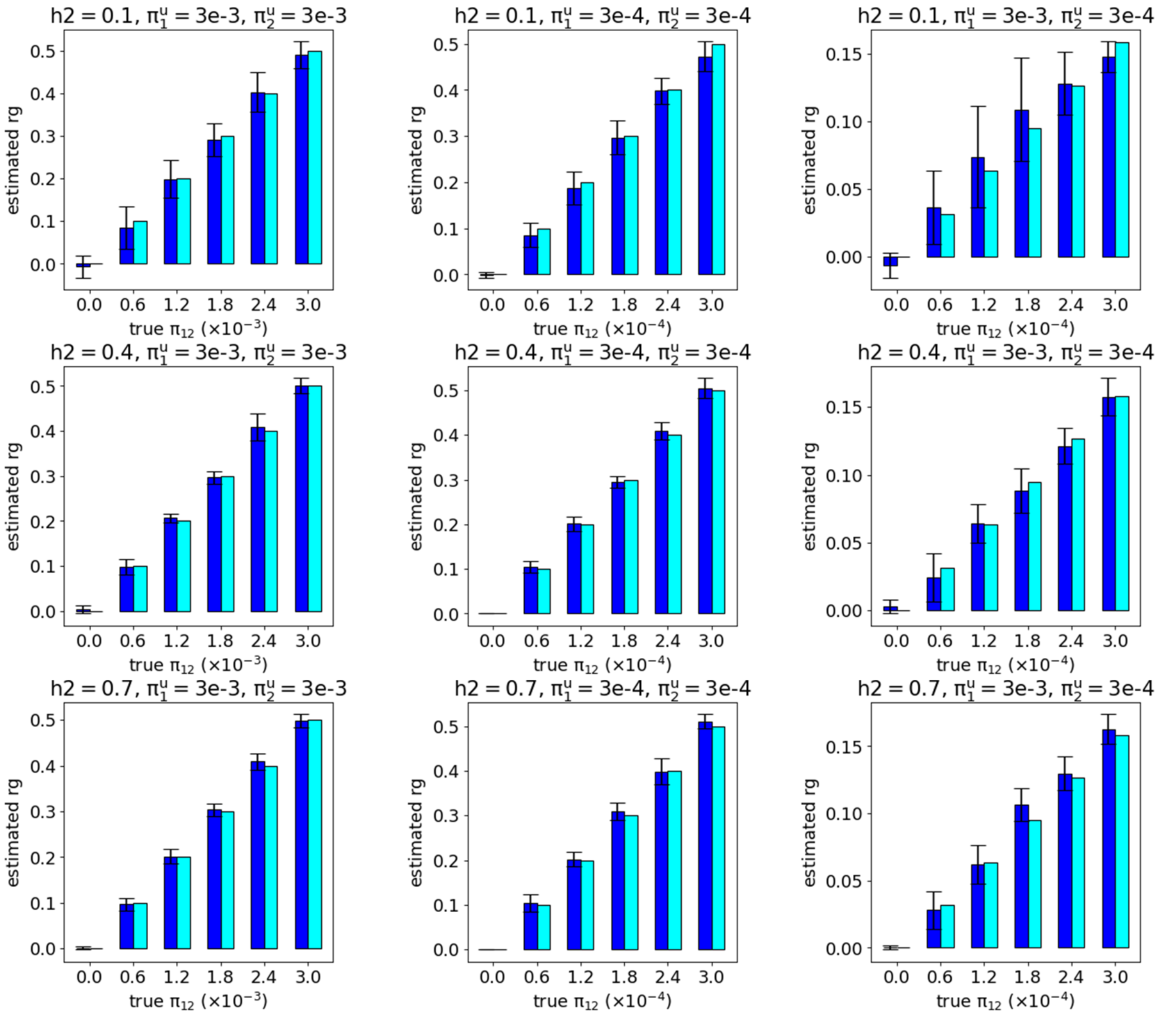


Simulation with synthetic GWAS summary statistics, generated according to MiXeR model, showing accuracy of $r_g = \rho_{12}\pi_{12}/\sqrt{\pi_1^u \pi_2^u}$ parameter in scenarios with genetic correlation (simulated rho12=0.5).

#### 4. Simulations: polygenicity and heritability estimates in univariate analysis

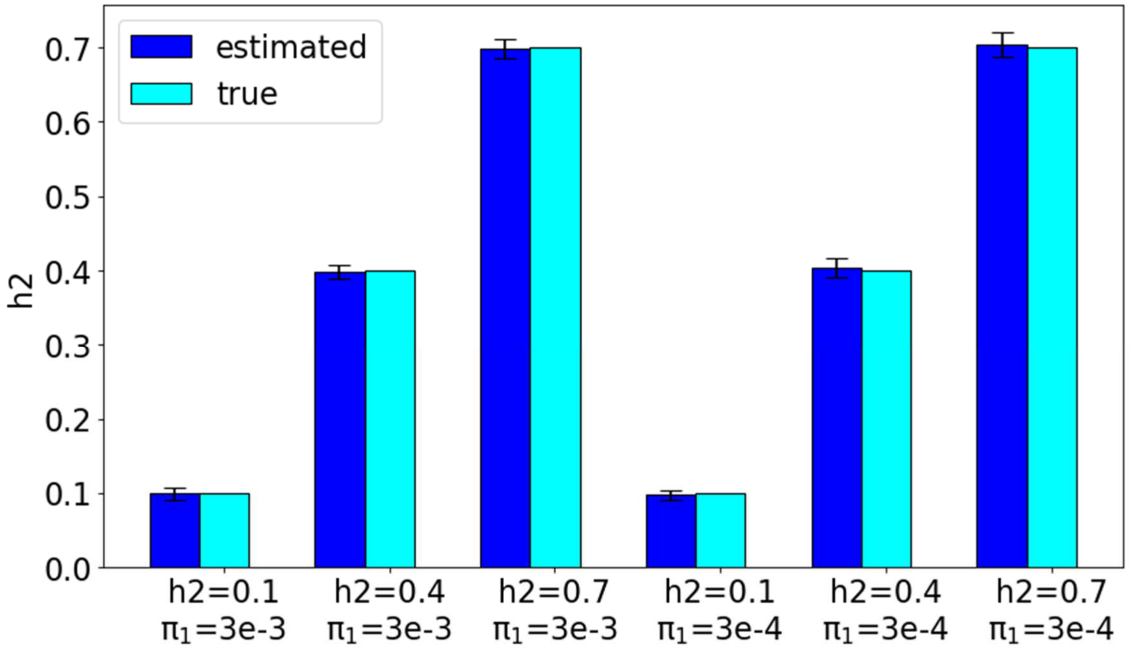


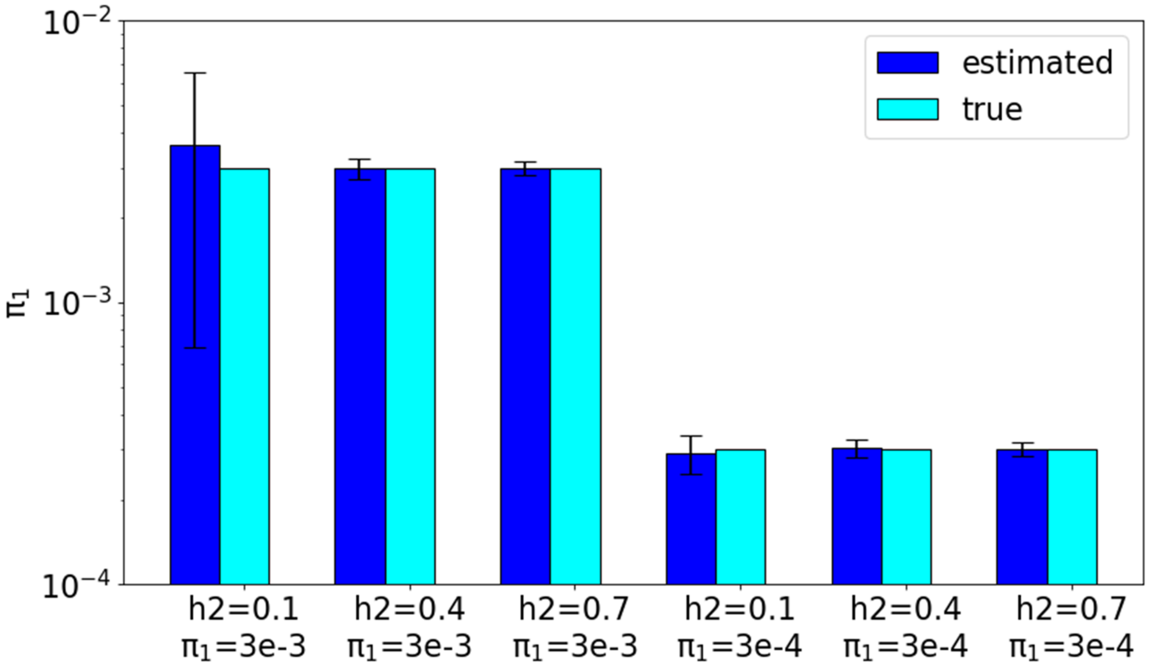


Simulations with univariate model. Top figure: validation of the heritability estimation. Bottom figure: validation of the polygenicity estimation. In total six scenarios are tested, with simulated heritability h2 set to 0.1, 0.4, 0.7 and polygenicity $\pi_1$ set to 3e-3, 3e-4. The bars in blue indicate an average value of model estimates across 120 simulation runs. The bars in cyan show true (simulated) parameters. Error bars represent standard deviation of the model estimate across 120 simulation runs.

#### 5. Simulations: Q-Q plots with simulated data

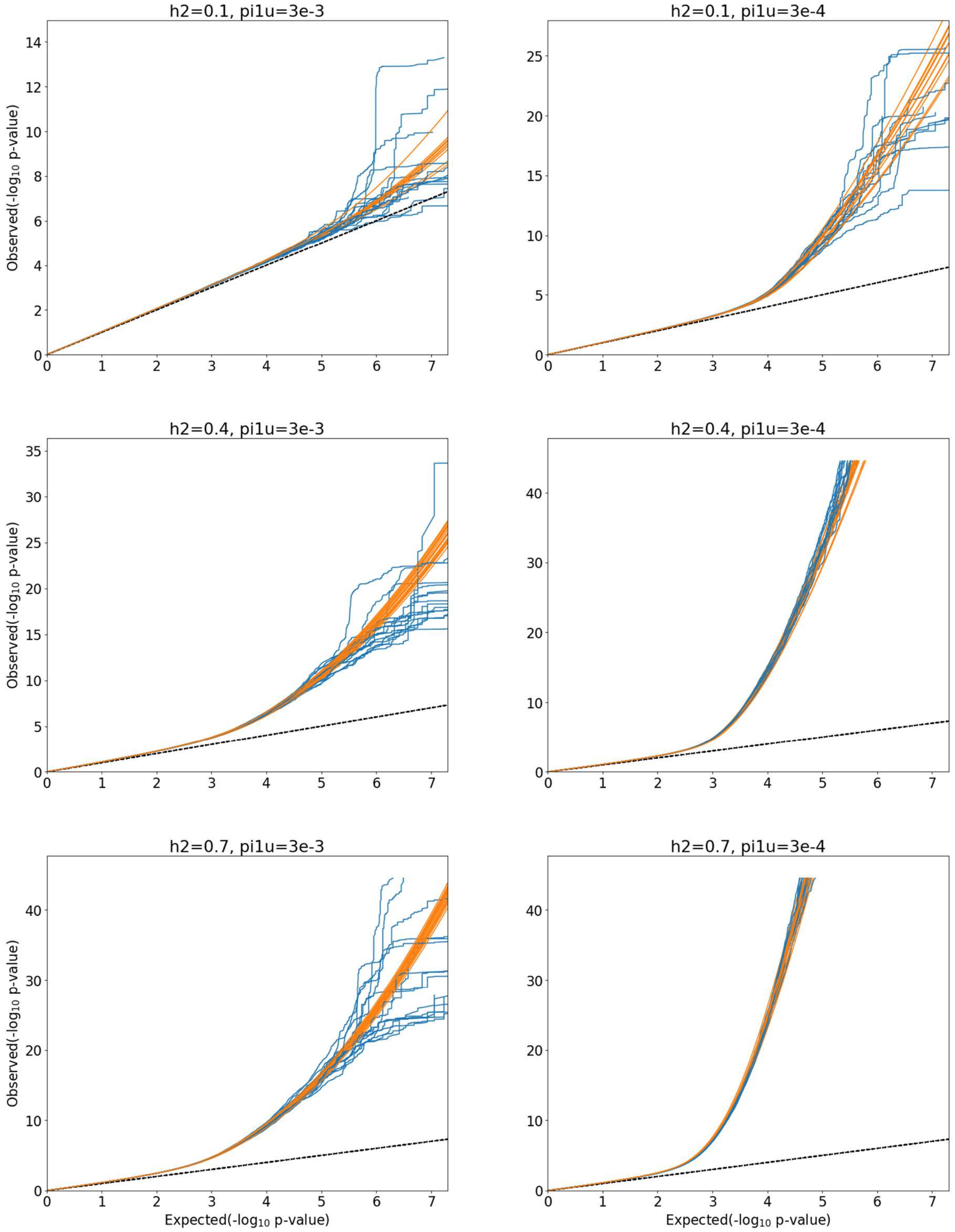


Simulated QQ plots and model prediction across six scenarios (two levels of polygenicity: 3e-3, 3e-4; three levels of heritability: 0.1, 0.4, 0.7). Each simulation was repeated 10 times with random instantiation of causal variants and their effects. Points on the QQ plot are weighted according to LD structure, using n=64 iterations of random pruning at LD threshold r2=0.1.

#### 6a. Simulations: Q-Q plots of SNPs partitioned into a grid of MAF and LD score, high polygenicity

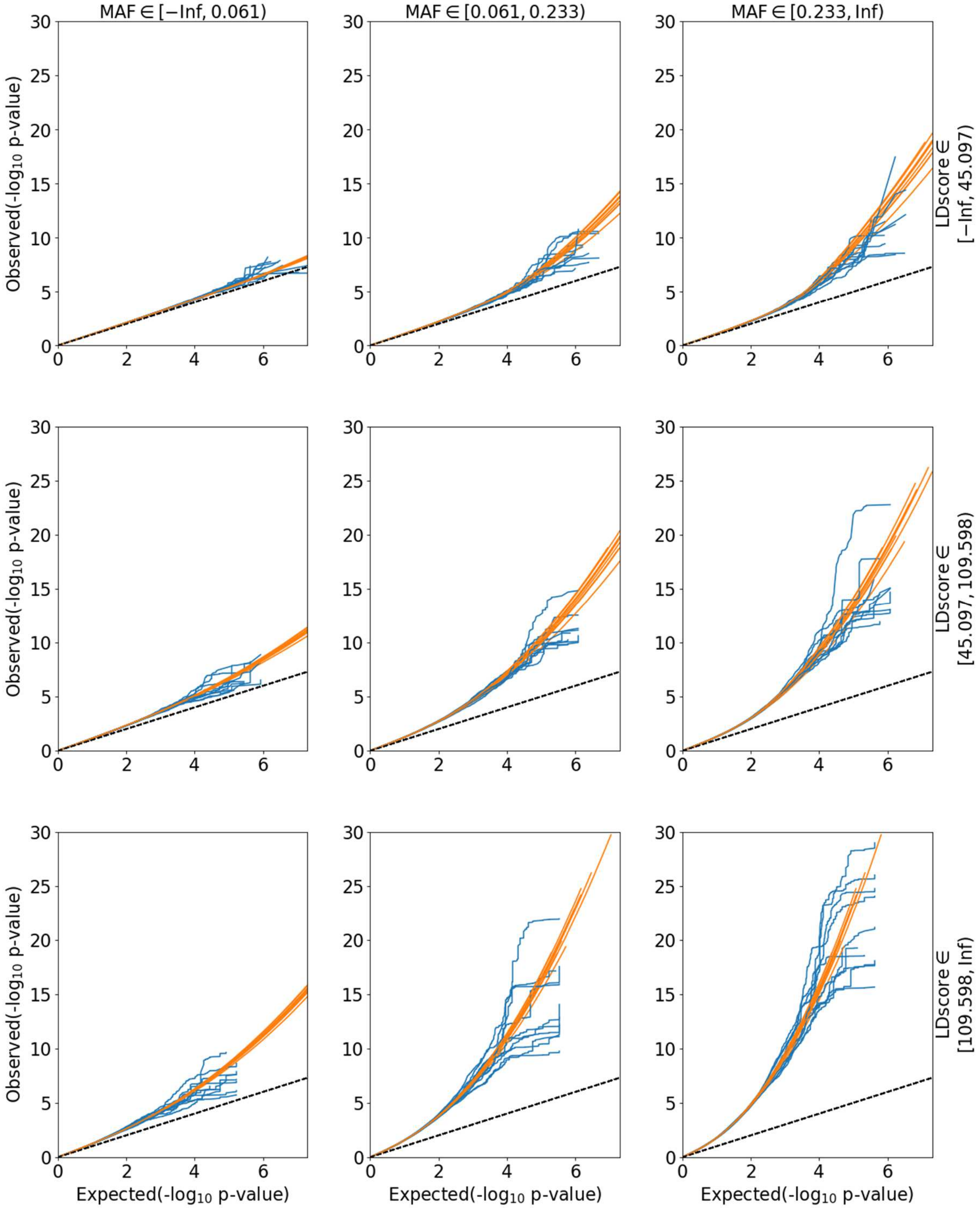


Simulated data. QQ plots for subsets of SNPs, partitioned into 9 groups according to minor allele frequency (MAF) and total LD score. Scenario with high polygenicity (pi1u=3e-03), heritability h2=0.4. Points on the QQ plot are weighted according to LD structure, using n=64 iterations of random pruning at LD threshold r2=0.1.

#### 6b. Simulations: Q-Q plots of SNPs partitioned into a grid of MAF and LD score, low polygenicity

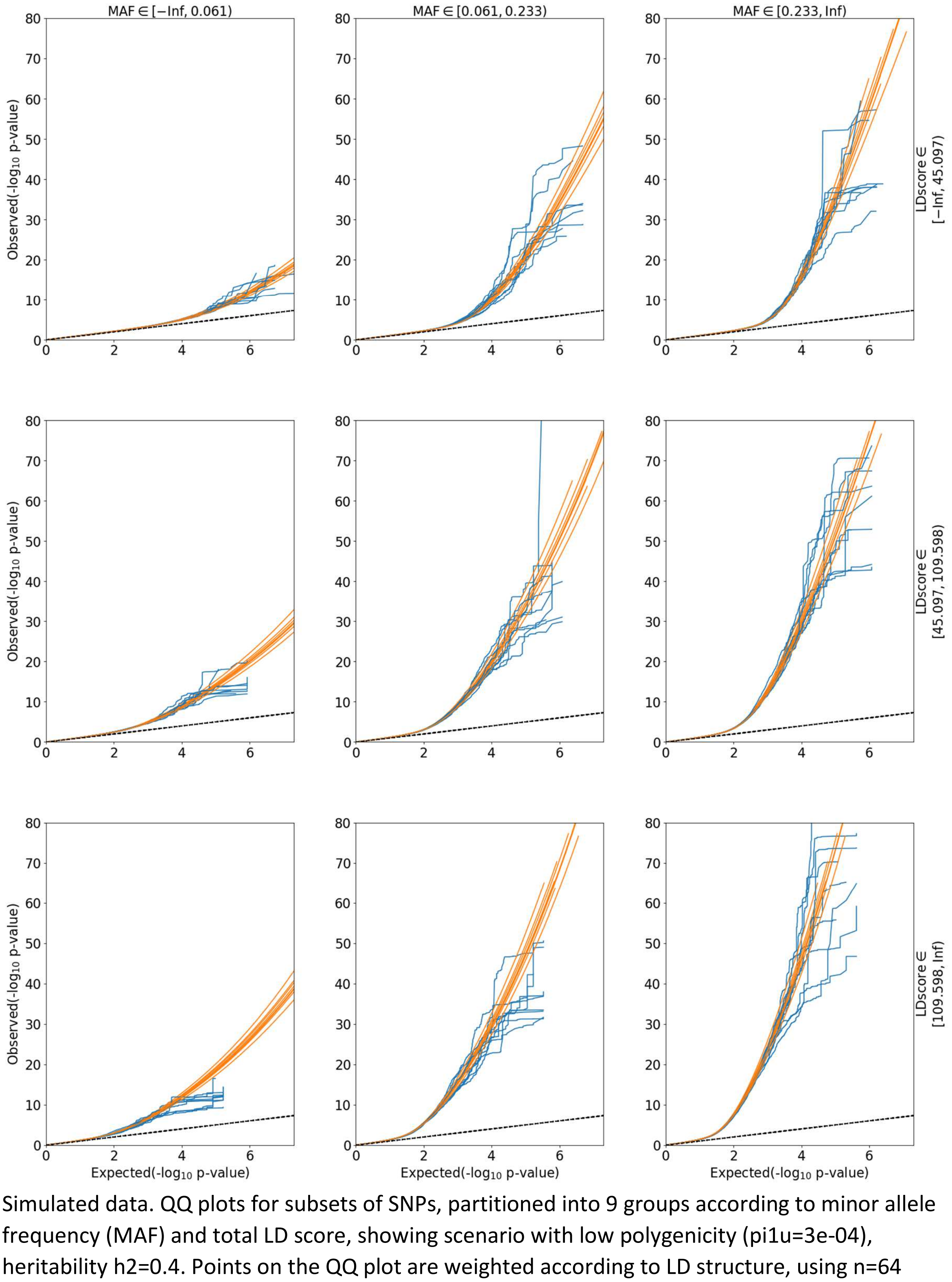


Simulated data. QQ plots for subsets of SNPs, partitioned into 9 groups according to minor allele frequency (MAF) and total LD score, showing scenario with low polygenicity (pi1u=3e-04), heritability h2=0.4. Points on the QQ plot are weighted according to LD structure, using n=64 iterations of random pruning at LD threshold r2=0.1.

#### 7. Simulations: stratified QQ plots for traits with and without polygenic overlap

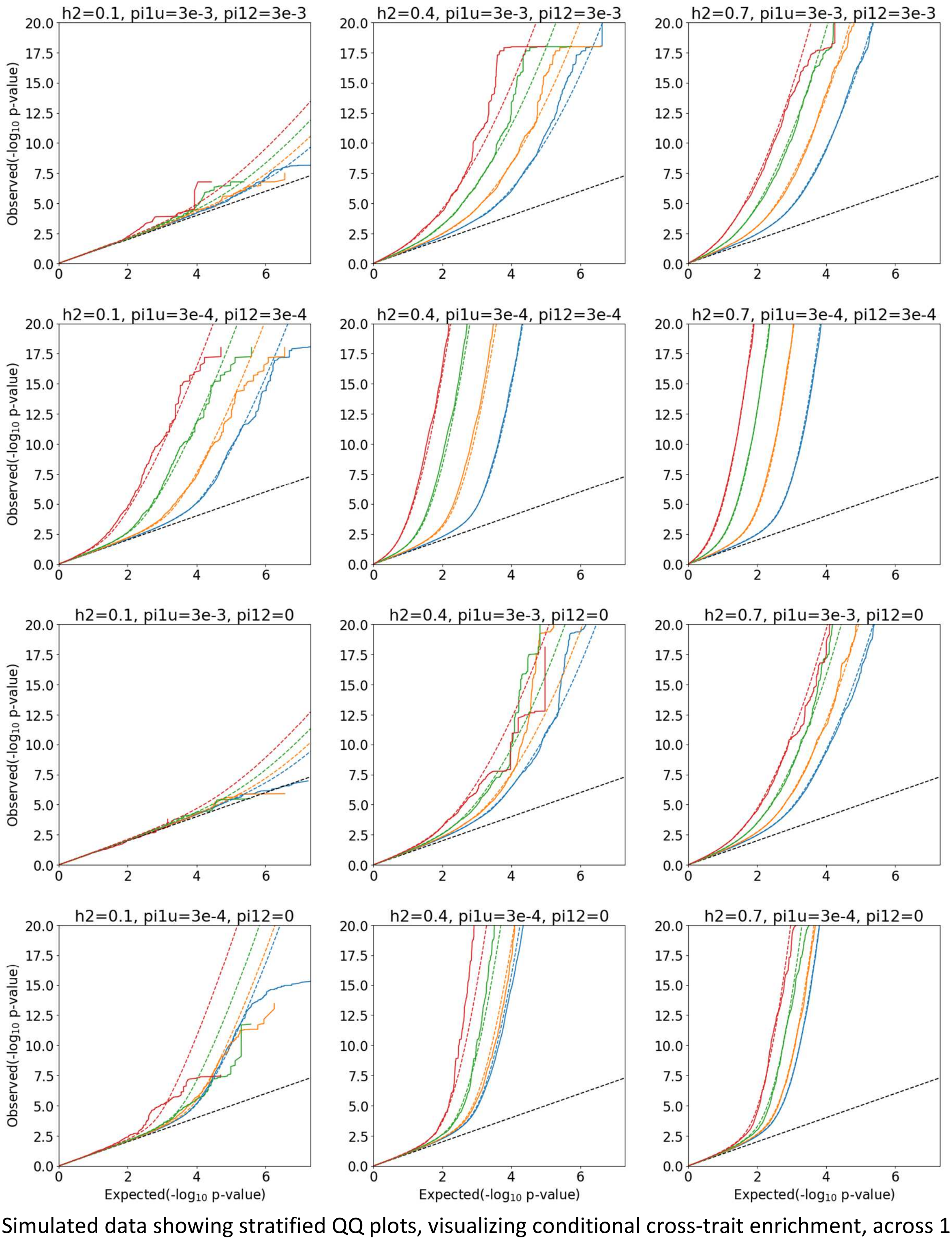


Simulated data showing stratified QQ plots, visualizing conditional cross-trait enrichment, across 12 scenarios (two levels of polygenicity: 3e-3, 3e-4; three levels of heritability: 0.1, 0.4, 0.7, with and without polygenic overlap). Top six figures represent scenarios with polygenic overlap at causal level. Bottom six figures are showing scenarios without polygenic overlap. Points on the QQ plot are weighted according to LD structure, using n=64 iterations of random pruning at LD threshold r2=0.1.

#### 8a. Sensitivity analysis: differential enrichment profile of synthesized data, simulated with MAF architecture following MiXeR assumptions

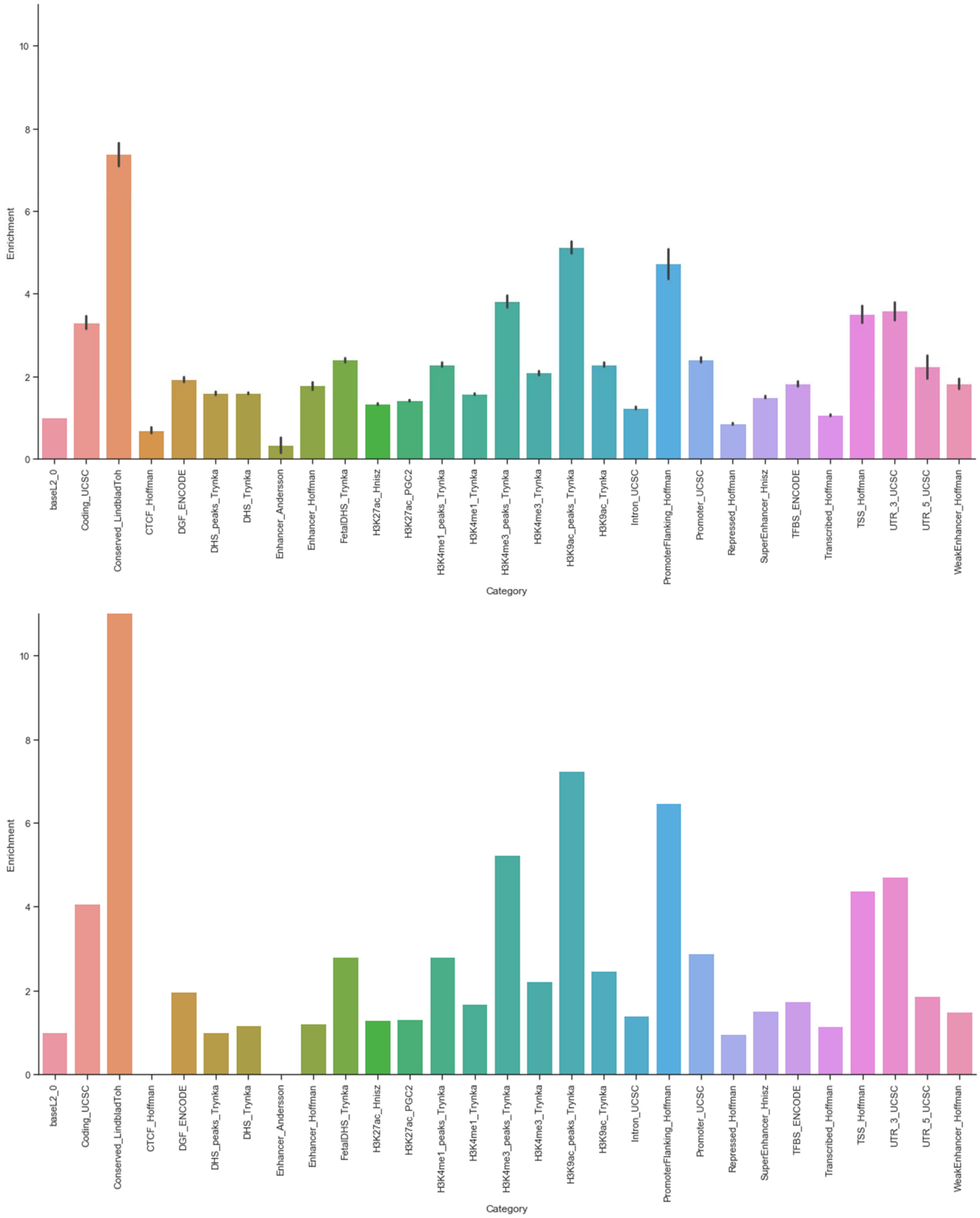


Comparison of enrichment between simulated data (top) and enrichment of Schizophrenia (bottom), estimated with stratified LDSR using `--per-allele` option for LD score estimation

#### 8b. Sensitivity analysis: differential enrichment profile of synthesized data, simulated with MAF architecture following LDSR assumptions

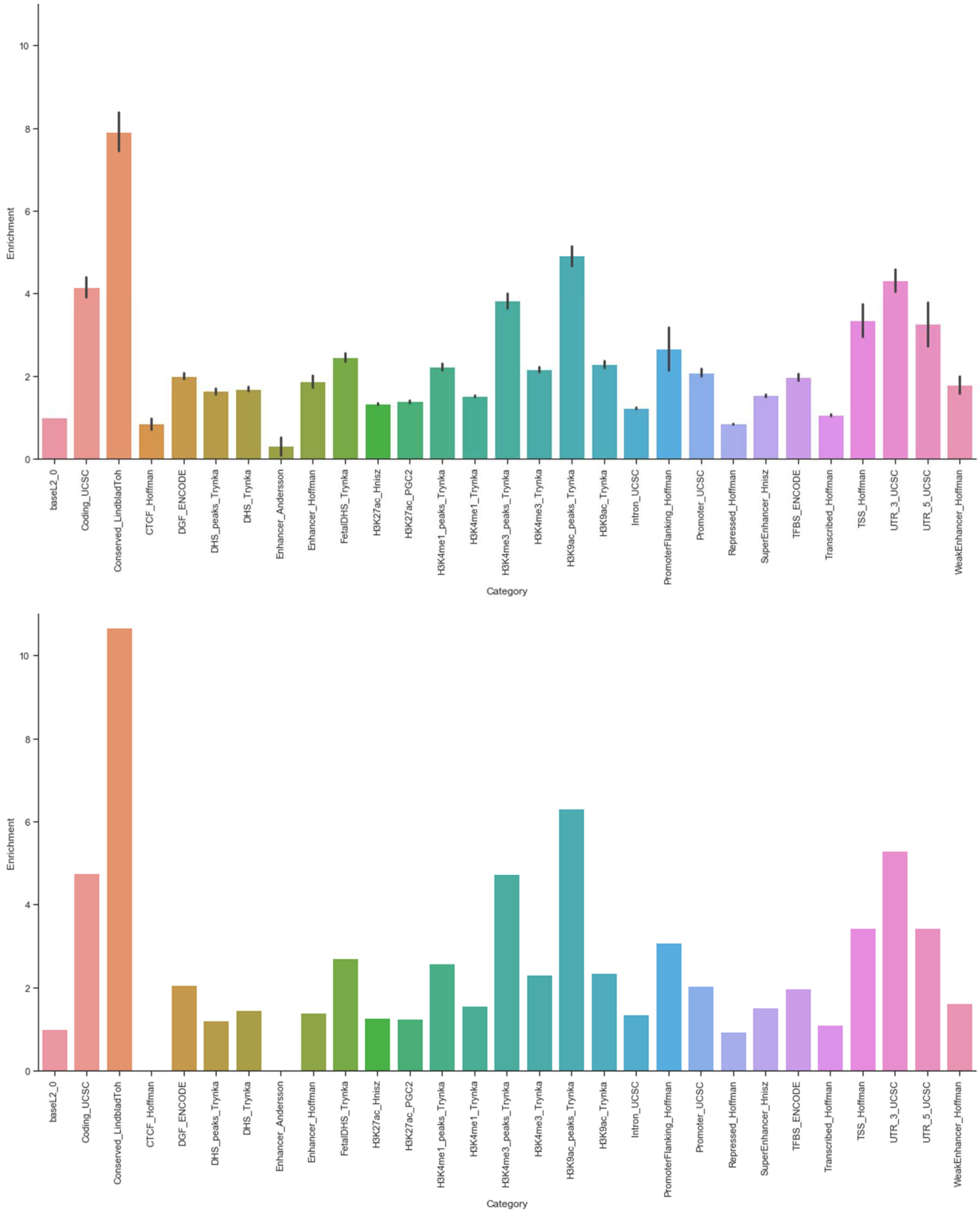


Comparison of enrichment between simulated data (top) and enrichment of Schizophrenia (bottom), estimated with stratified LDSR.

#### 9. Sensitivity analysis: polygenicity and heritability estimates under differential genomic enrichment, with MAF architecture simulated following MiXeR assumptions

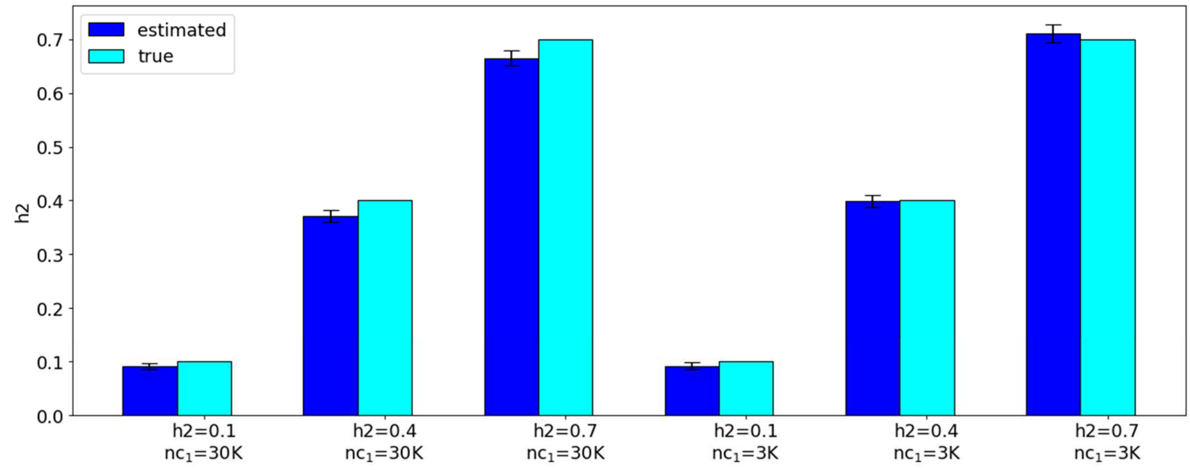


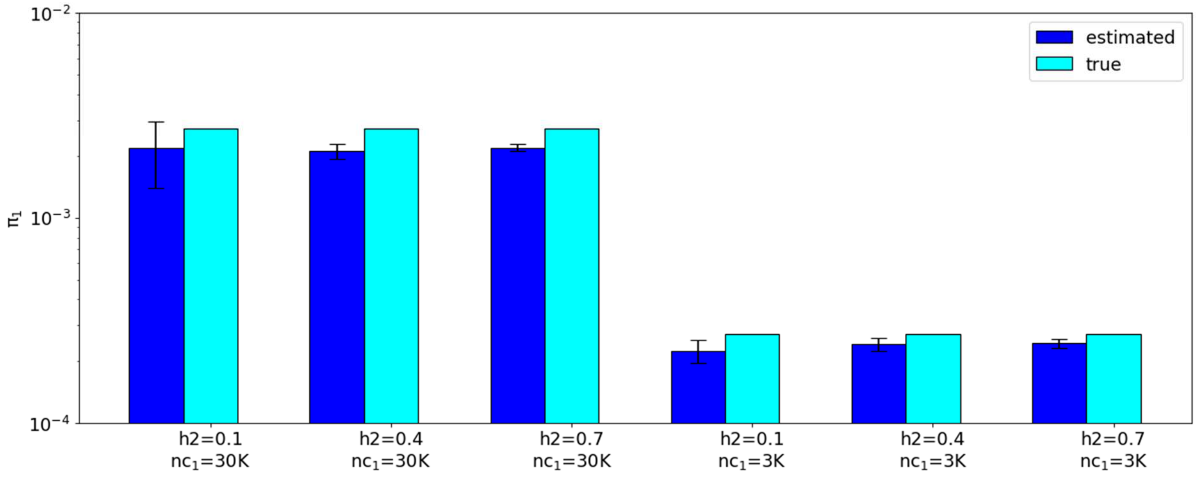


Simulations with model misspecification showing a minor bias in univariate estimates in the presence of genomic annotations. Top figure: validation of the heritability estimation. Bottom figure: validation of the polygenicity estimation. The results of bivariate analysis are presented in a separate table.

#### 10. Sensitivity analysis: polygenicity and heritability estimates under differential genomic enrichment, with MAF architecture simulated following LDSR assumptions

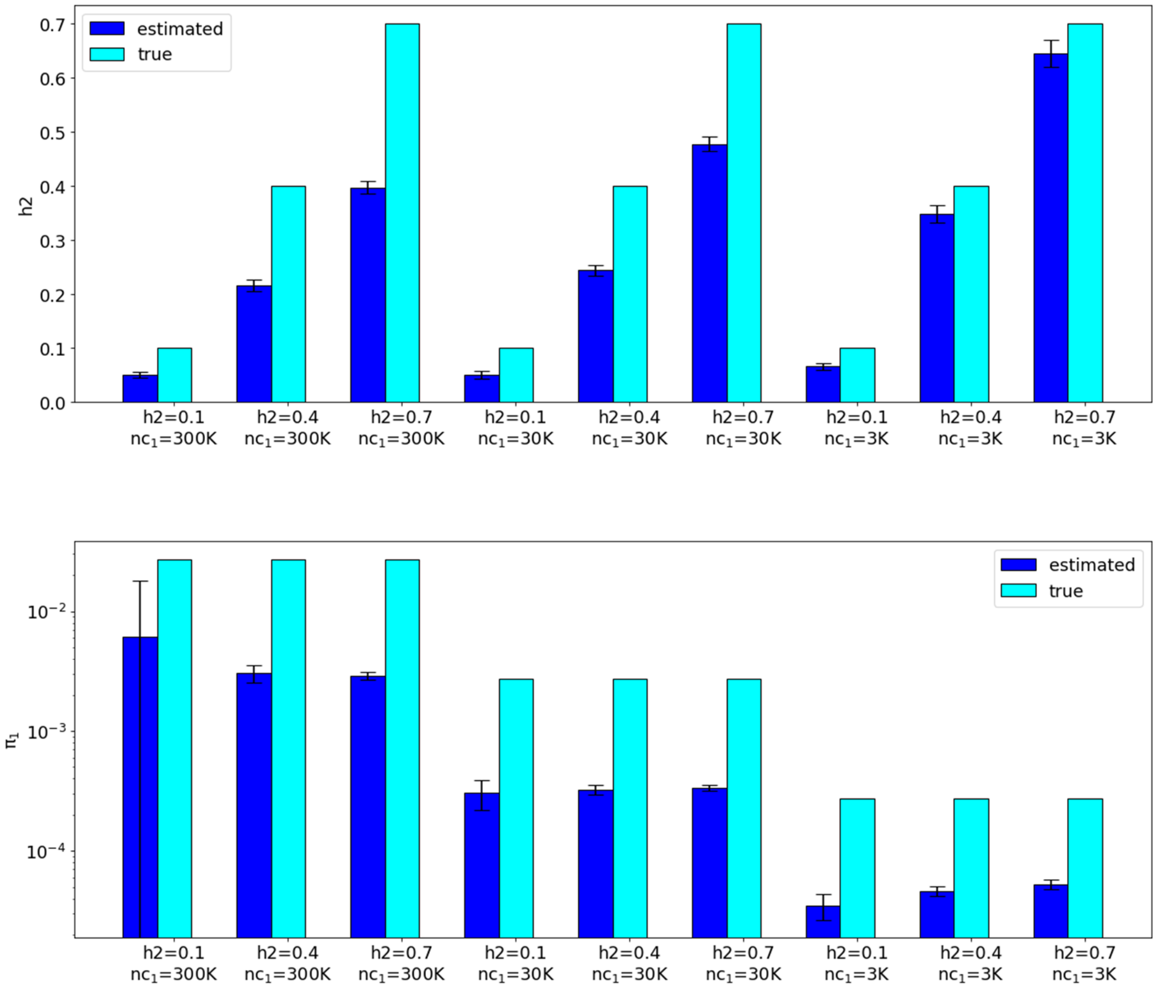


Simulations with model misspecification showing significant bias in univariate estimates in the presence of genomic annotations and extreme misspecification of MAF-dependent architecture. Top figure: validation of the heritability estimation. Bottom figure: validation of the polygenicity estimation. The results of bivariate analysis are presented in a separate table.

#### 11. Cumulated fraction of explained heritability versus proportion of causal variants

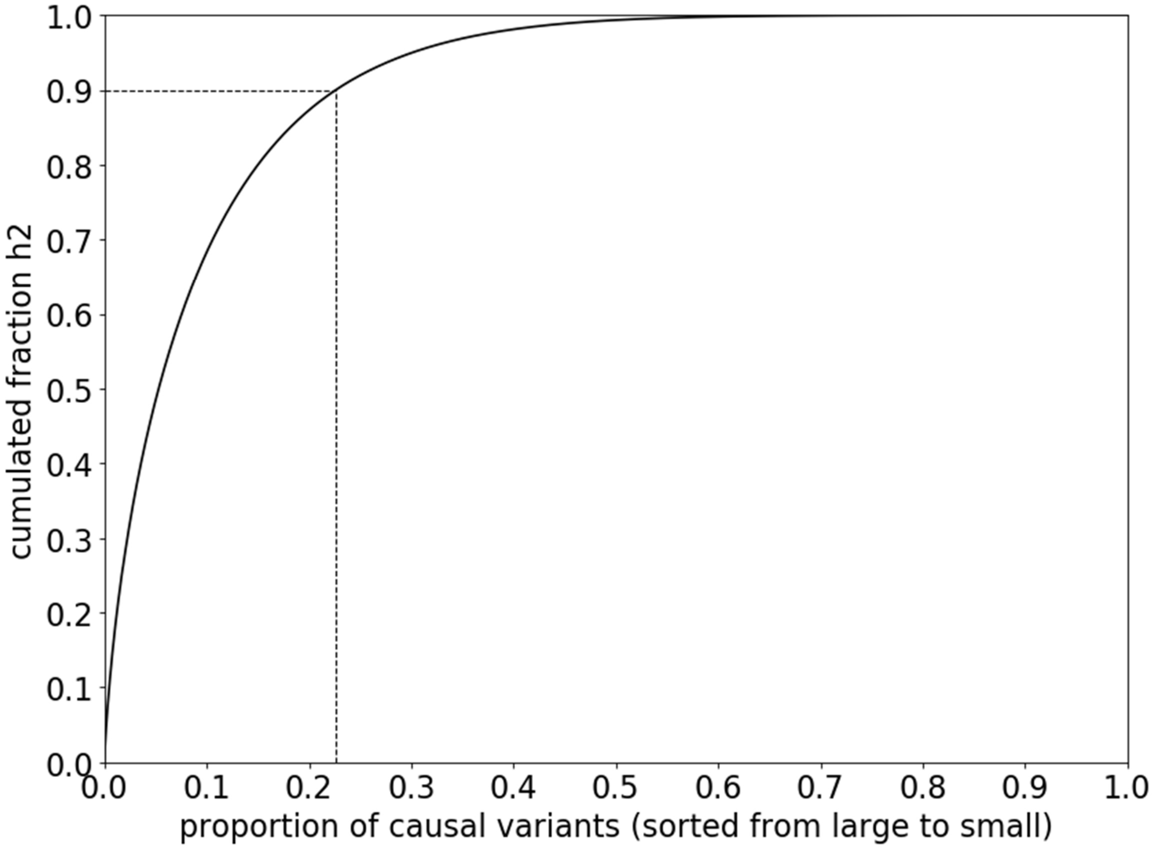


In MiXeR model the contribution of causal variant to heritability depends on the effect of the variant, and on its minor allele frequency: $h_2 = \beta_j^2 \times 2p_j(1 - p_j)$. Since many SNPs have low allele frequency, only a small fraction (22.6%) of causal variants with relatively large effects explains 90% of the total heritability.

#### 12. Venn diagrams across all traits

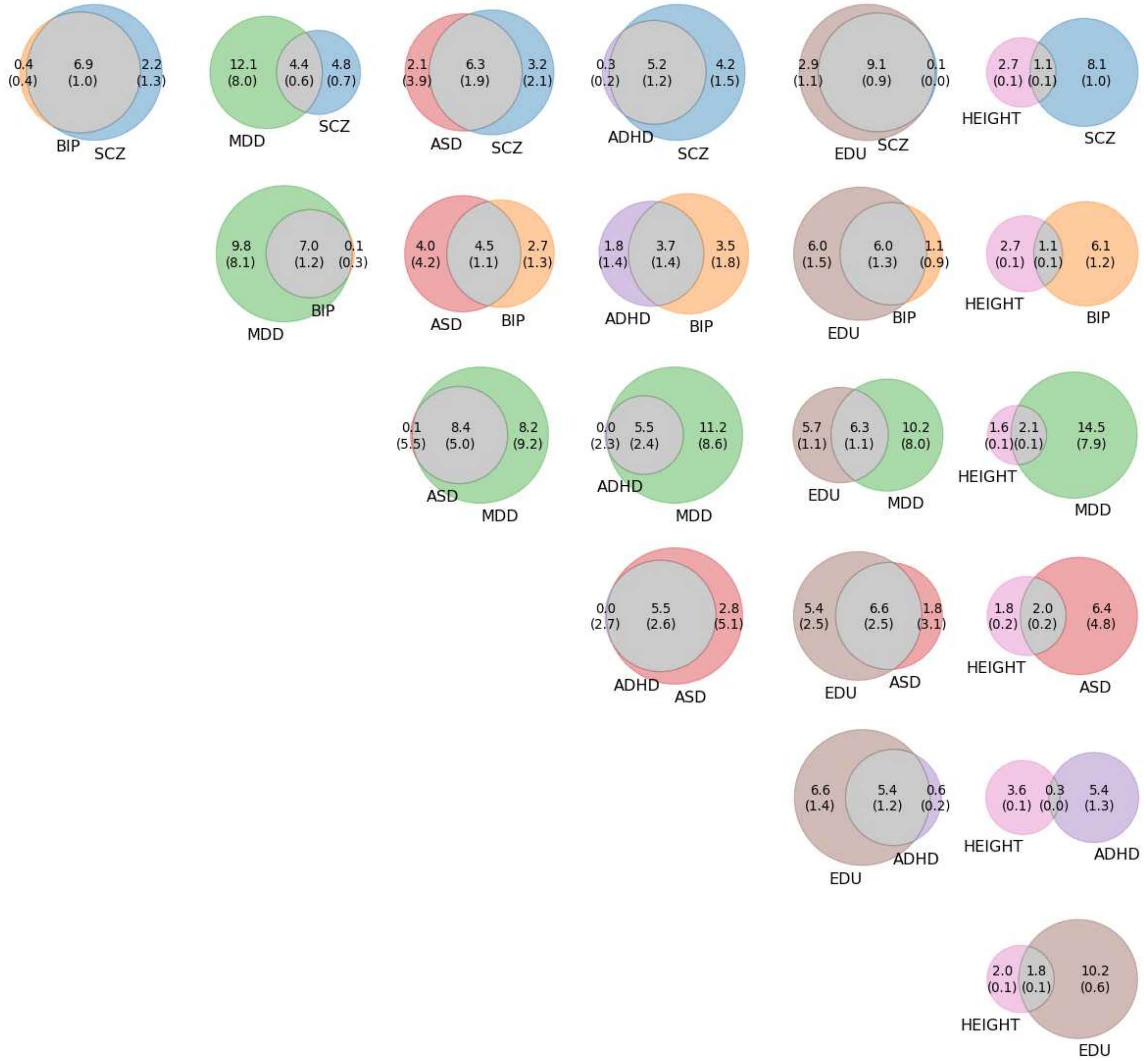


Venn diagrams of unique and shared polygenic component at the causal level. SCZ: schizophrenia; BIP: bipolar disorder; MDD: major depressive disorder; ASD: autism spectrum disorder; ADHD: attention deficit / hyperactivity disorder; EDU: educational attainment. The numbers indicate estimated quantity of causal variants (in 1,000) per component, explaining 90% of SNP heritability in each phenotype, followed by the standard error. The size of circles reflects the polygenicity (scaled individually for each of the Venn diagrams).

#### 13a. Venn diagrams and conditional cross-trait QQ plots for schizophrenia

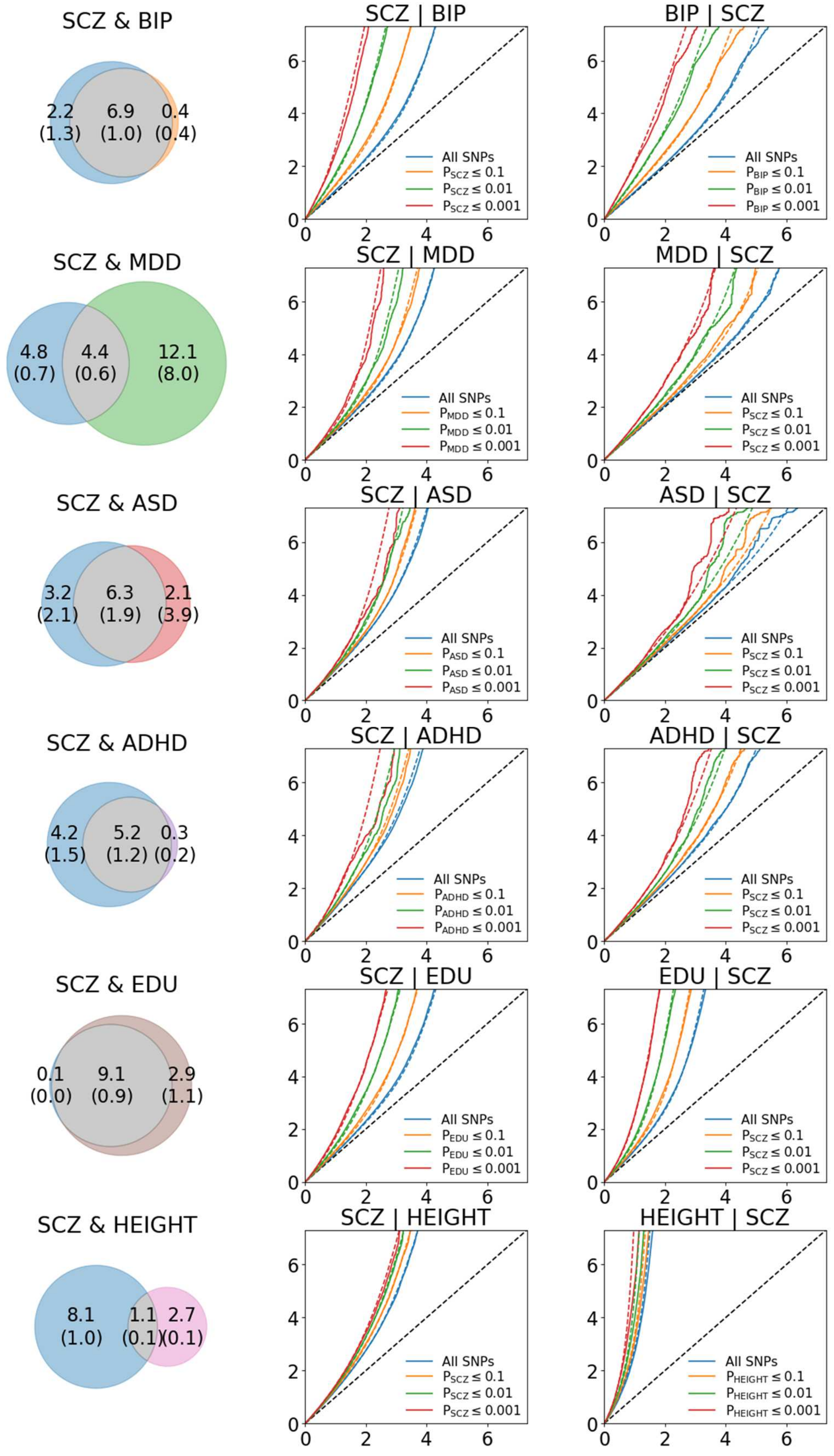


Venn Diagrams and stratified QQ plots, visualizing conditional cross-trait enrichment.

#### 13b. Venn diagrams and conditional cross-trait QQ plots for bipolar disorder

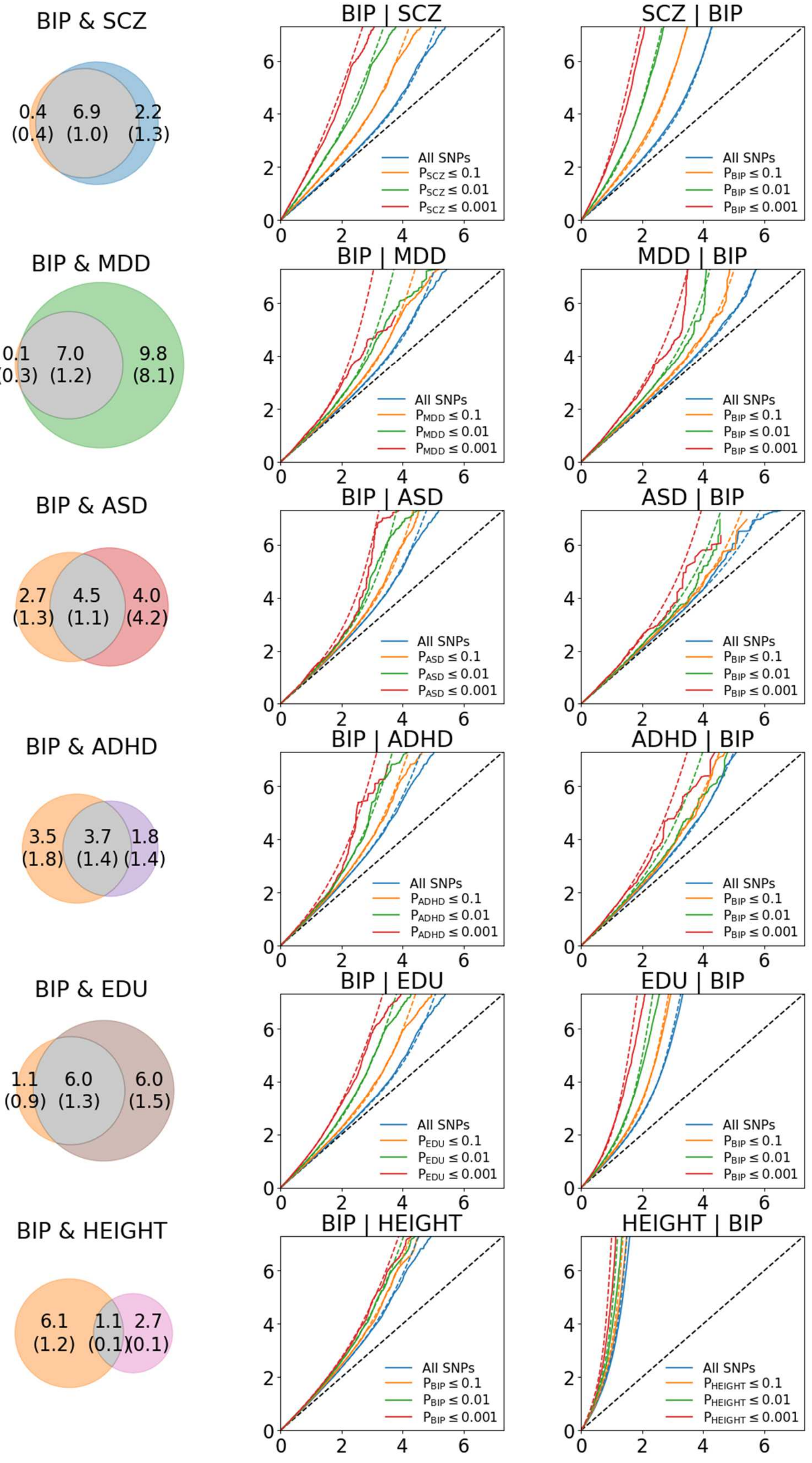


Venn Diagrams and stratified QQ plots, visualizing conditional cross-trait enrichment.

#### 13c. Venn diagrams and conditional cross-trait QQ plots for major depressive disorder

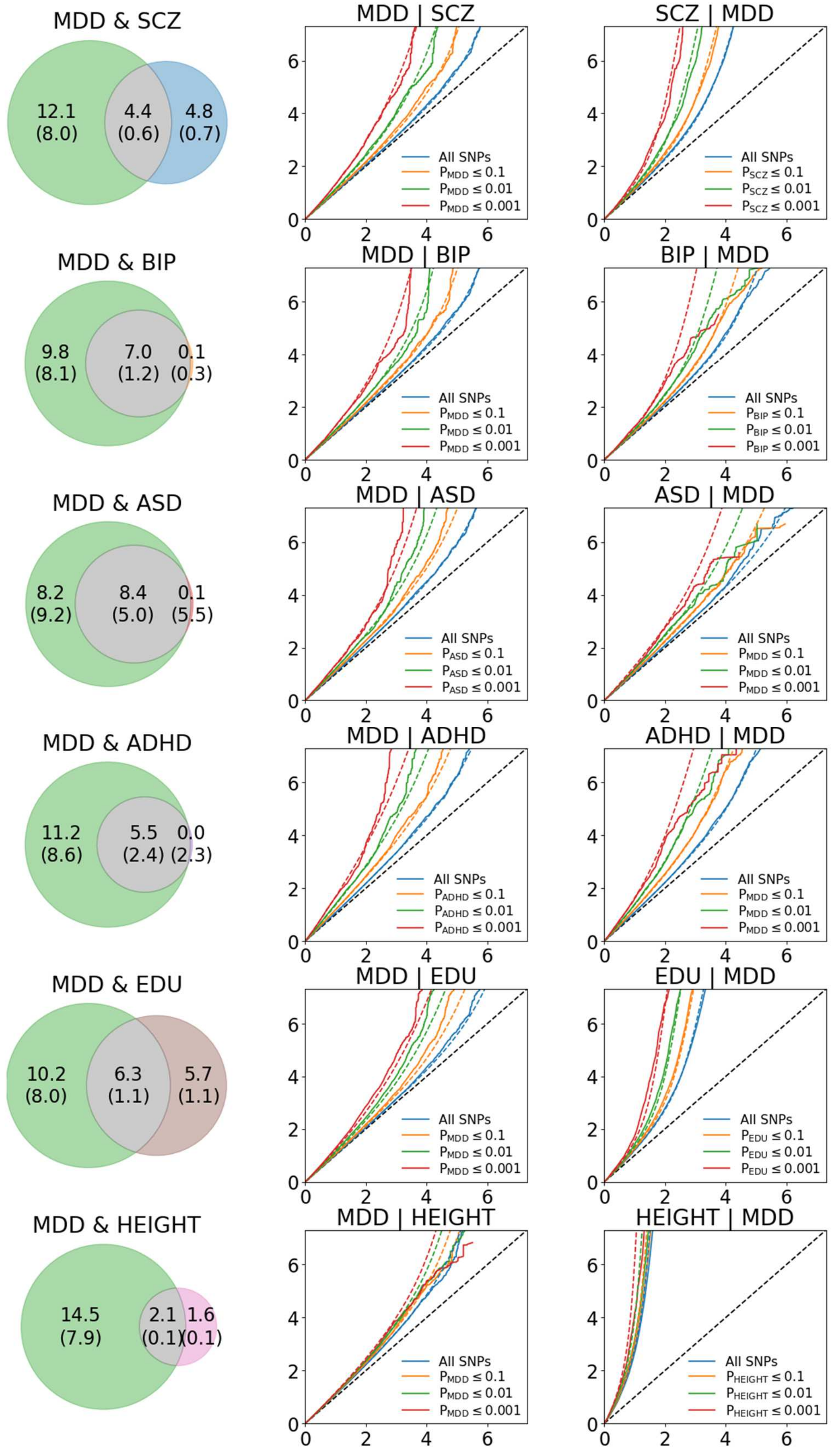


Venn Diagrams and stratified QQ plots, visualizing conditional cross-trait enrichment.

#### 13d. Venn diagrams and conditional cross-trait QQ plots for autism spectrum disorder

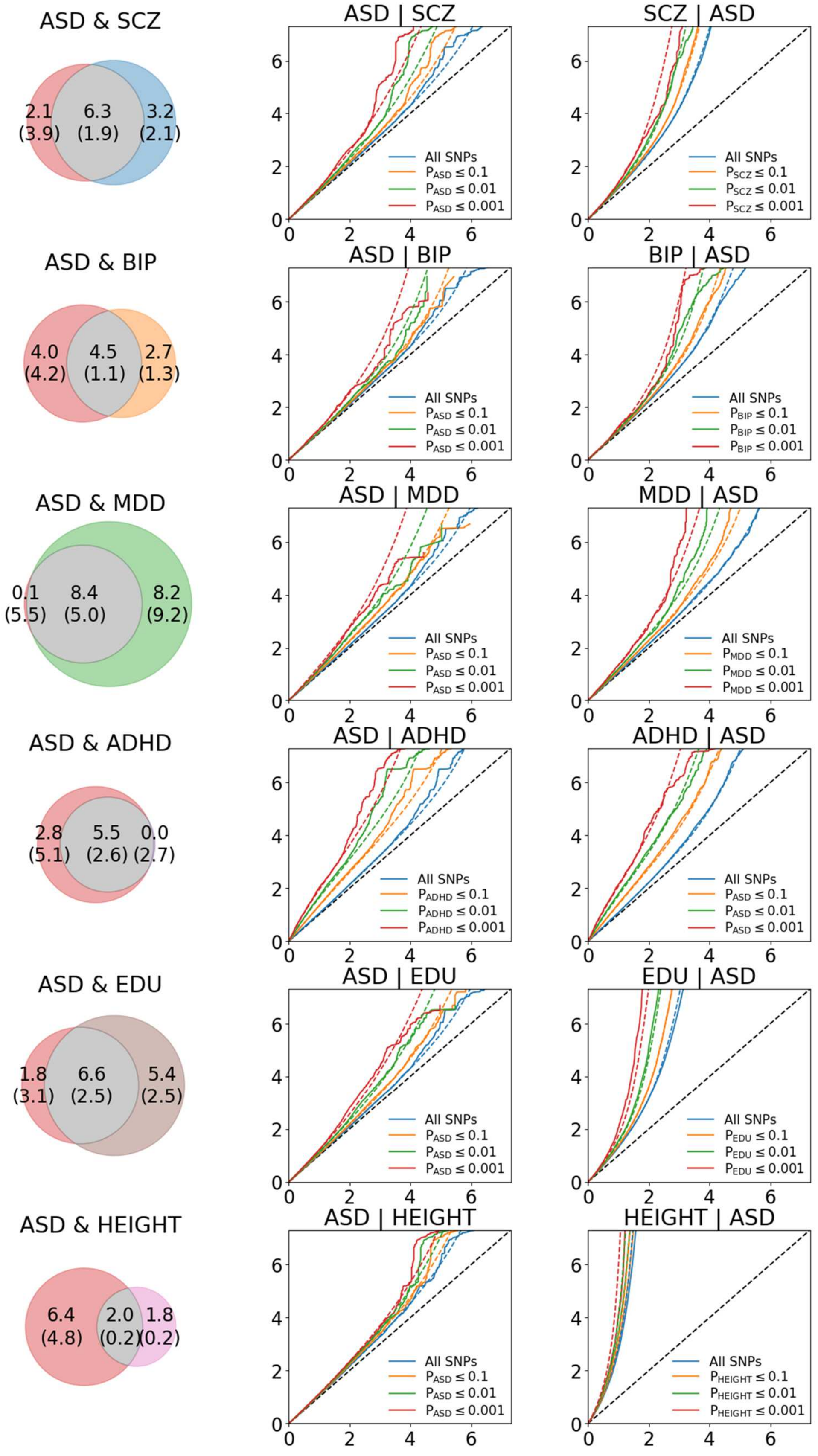


Venn Diagrams and stratified QQ plots, visualizing conditional cross-trait enrichment.

#### 13e. Venn diagrams and conditional cross-trait QQ plots for ADHD

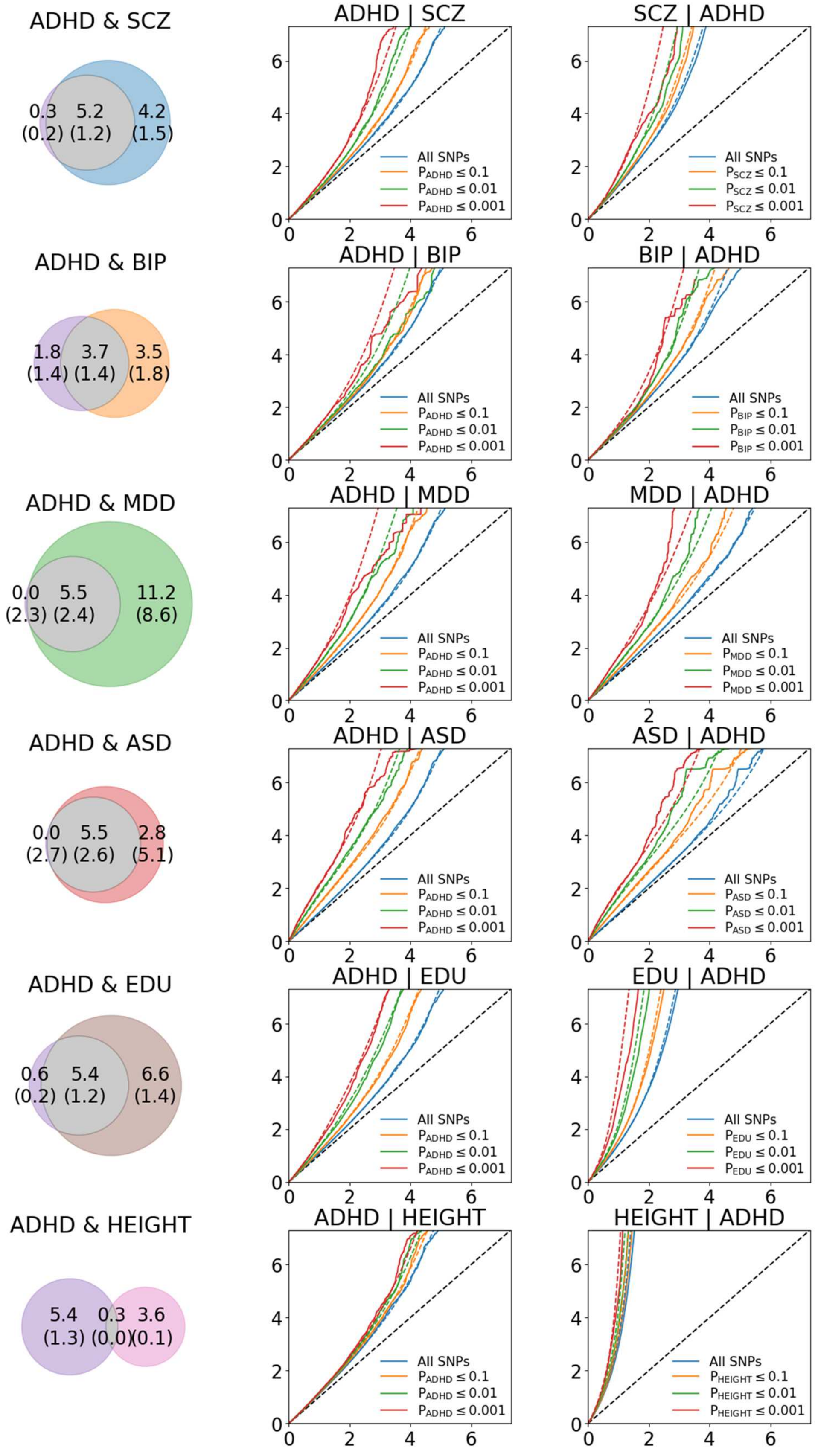


Venn Diagrams and stratified QQ plots, visualizing conditional cross-trait enrichment.

#### 13f. Venn diagrams and conditional cross-trait QQ plots for educational attainment

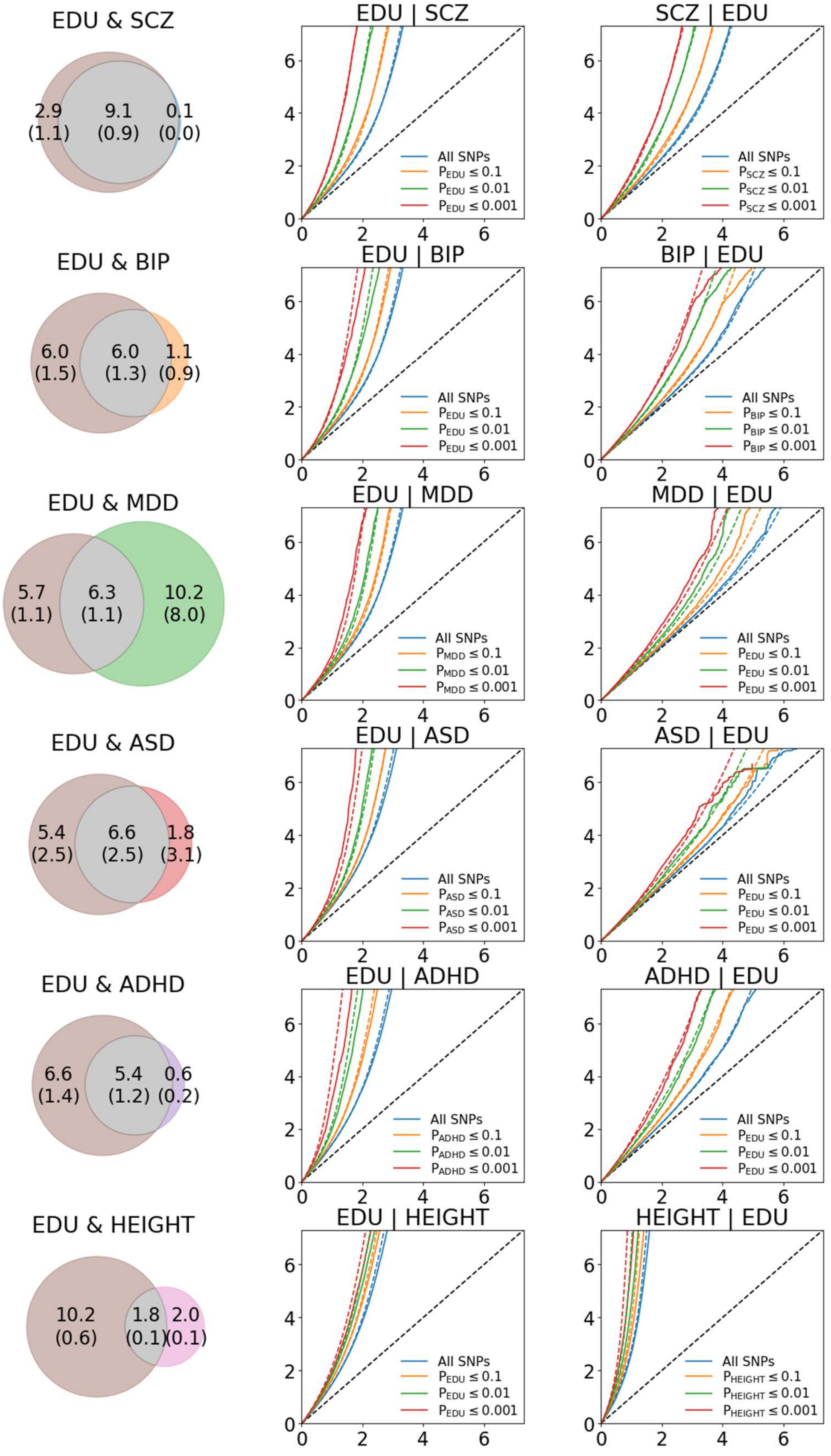


Venn Diagrams and stratified QQ plots, visualizing conditional cross-trait enrichment.

#### 13g. Venn diagrams and conditional cross-trait QQ plots for height

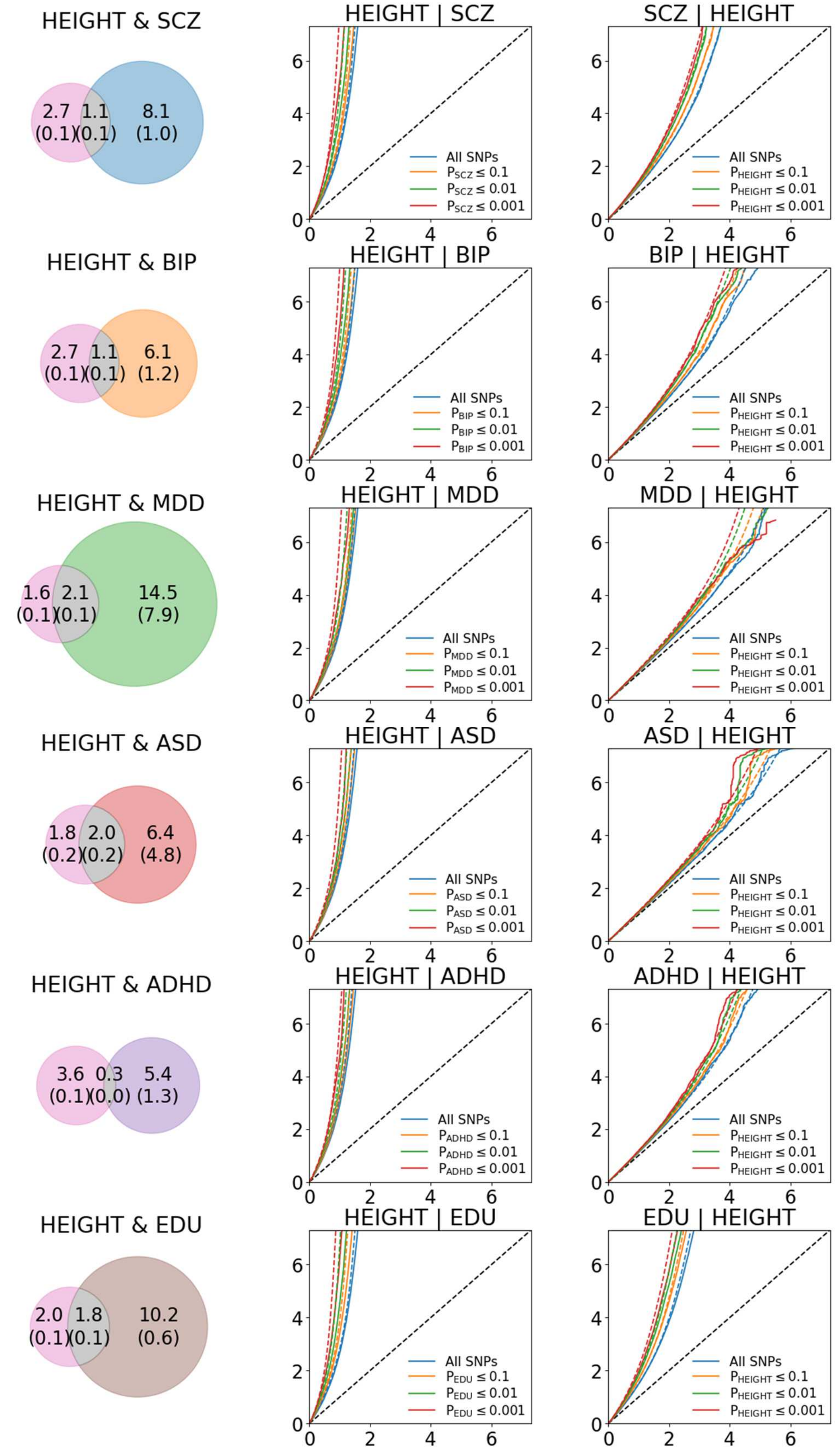


Venn Diagrams and stratified QQ plots, visualizing conditional cross-trait enrichment.

#### 14a. Observed and predicted bivariate density of GWAS association statistics, schizophrenia

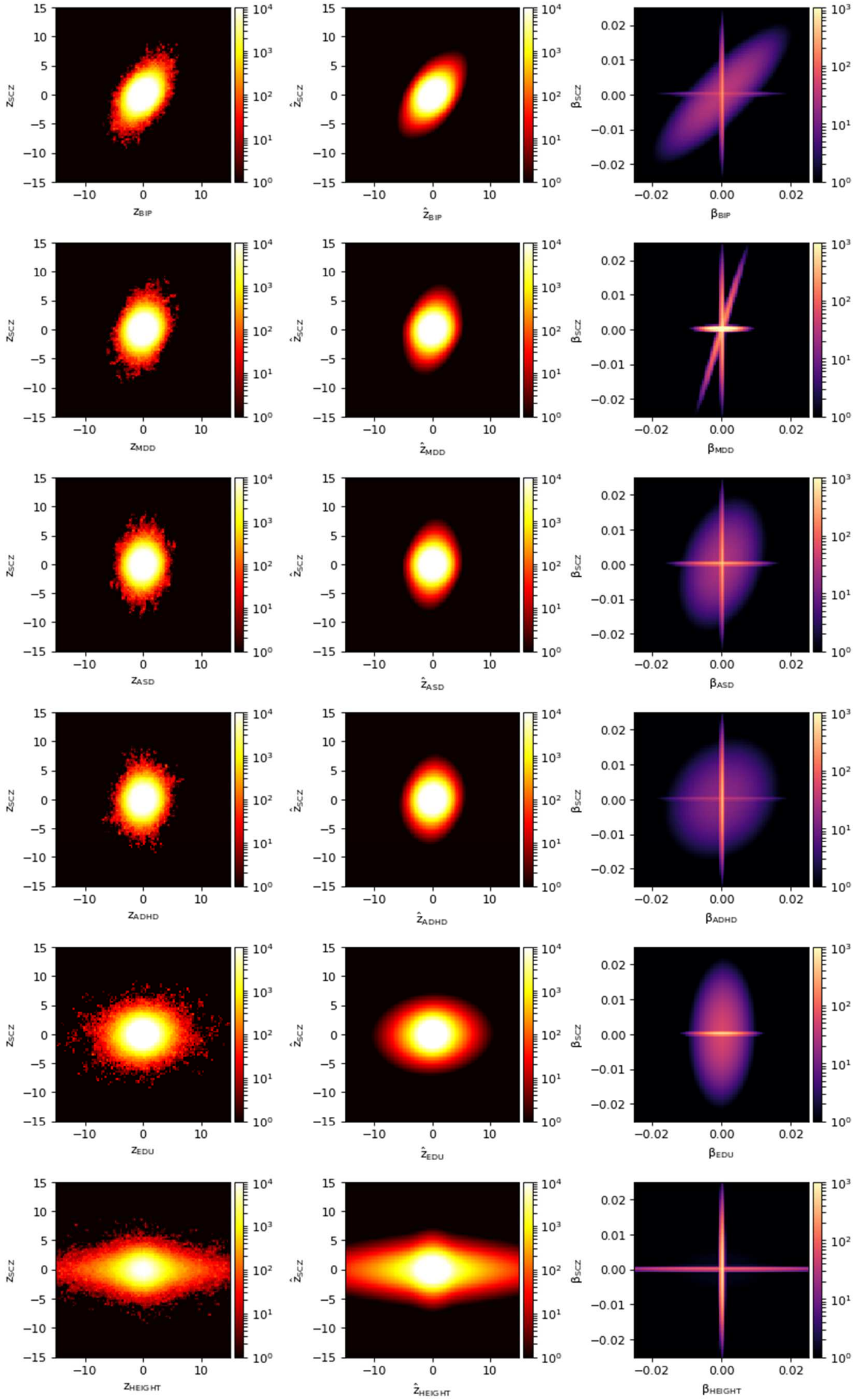

#### 14b. Observed and predicted bivariate density of GWAS association statistics, bipolar disorder

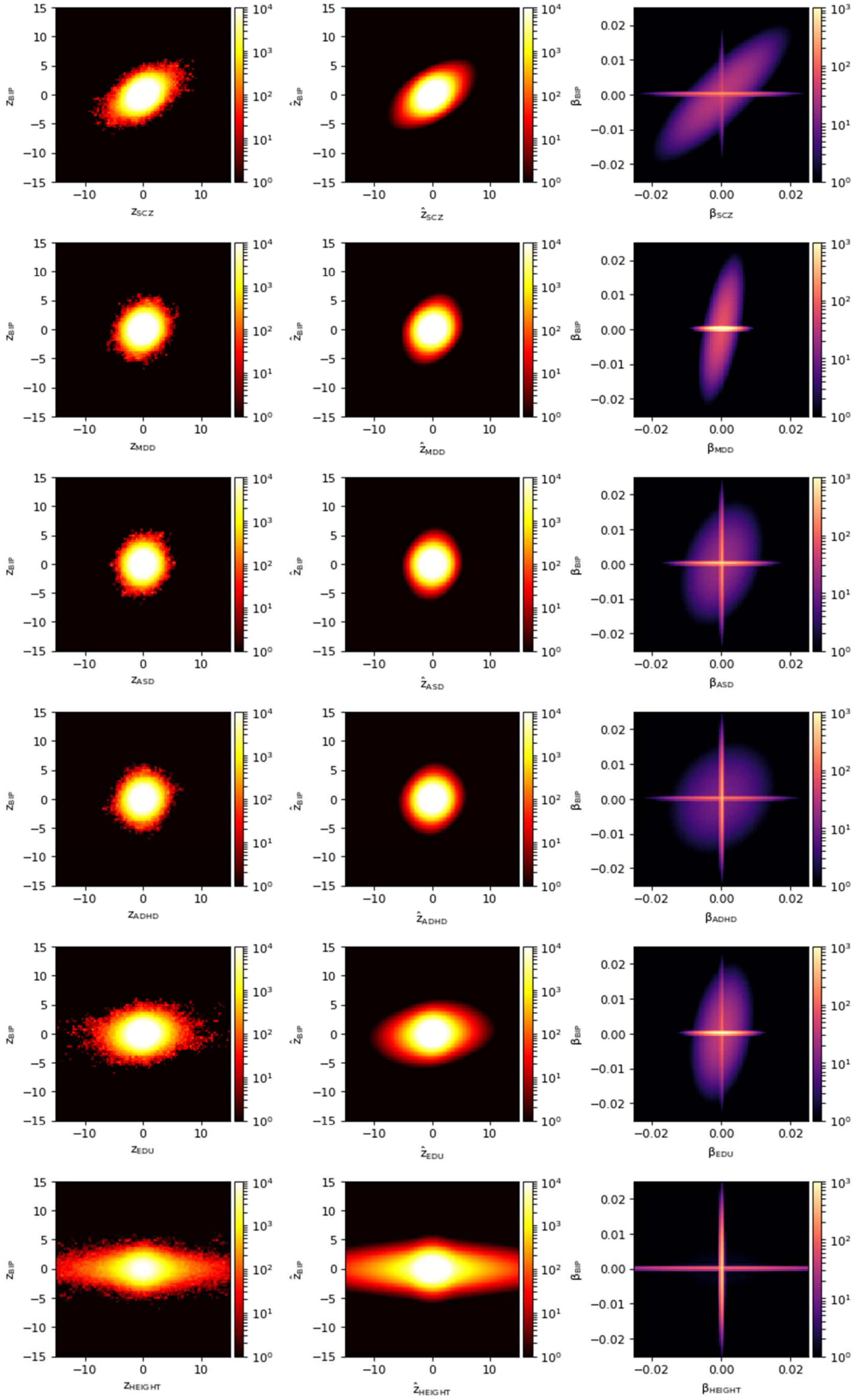

#### 14c. Observed and predicted bivariate density of GWAS association statistics, major depressive disorder

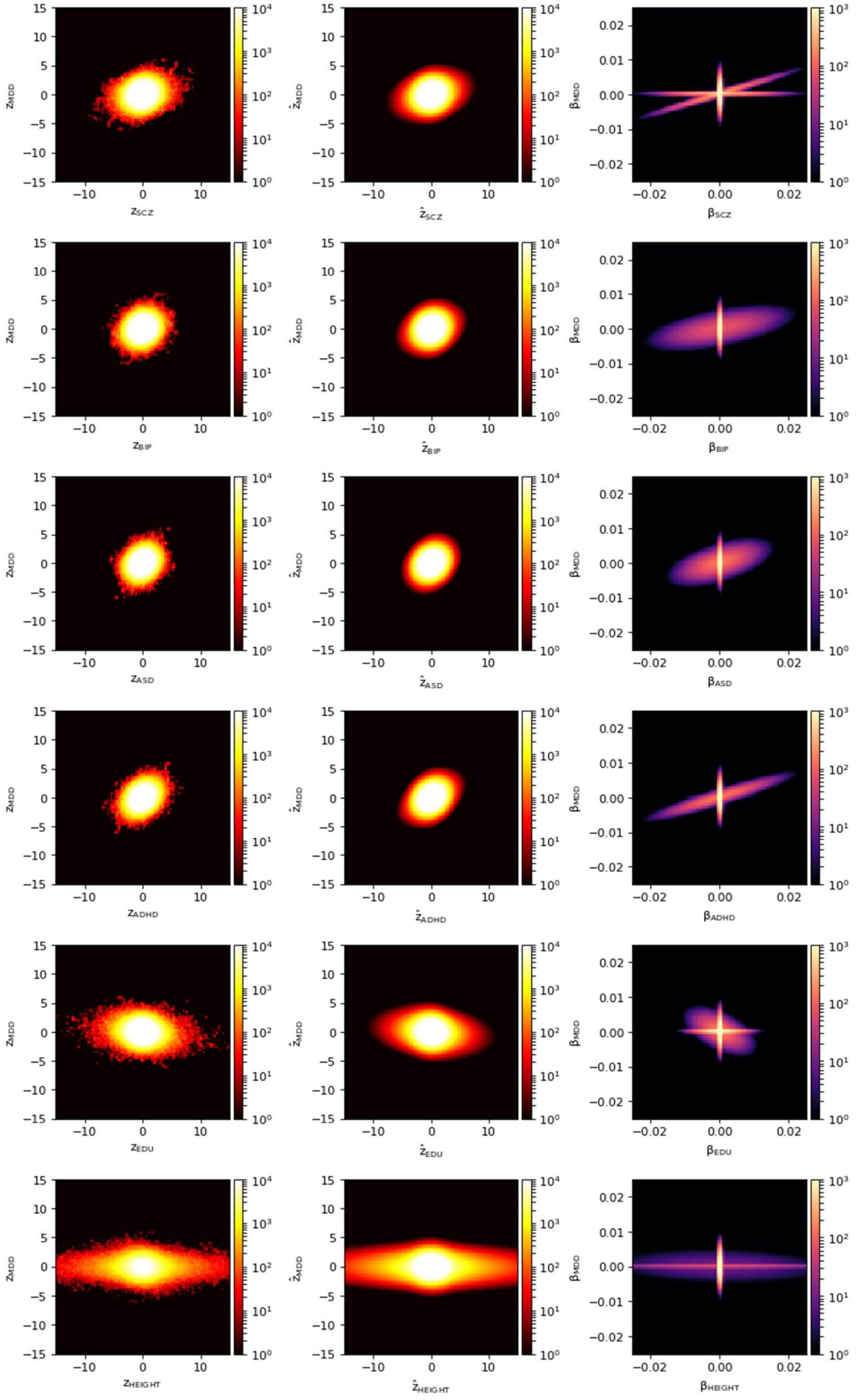

14d. Observed and predicted bivariate density of GWAS association statistics, autism spectrum disorder

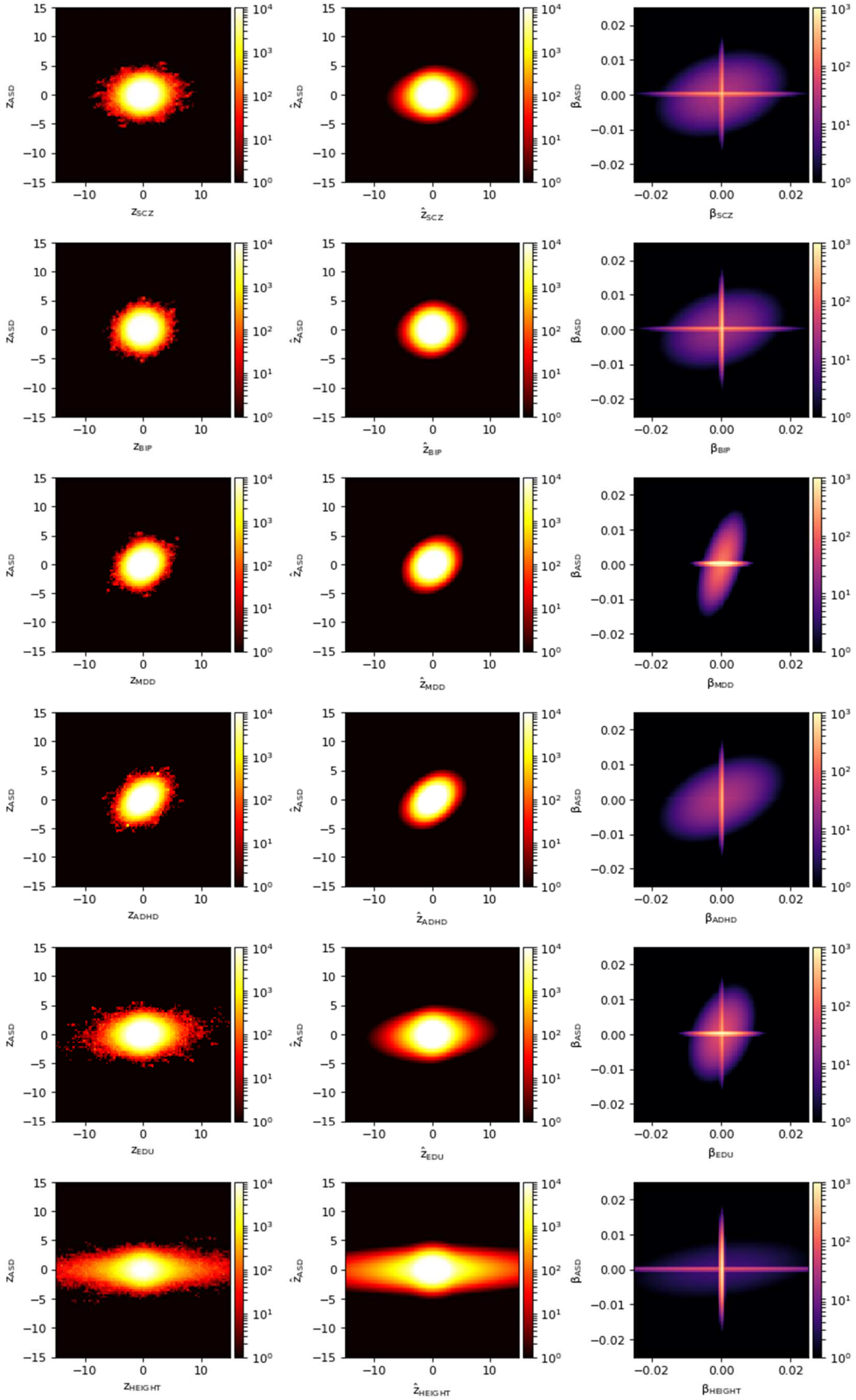

#### 14e. Observed and predicted bivariate density of GWAS association statistics, attention deficit / hyperactivity disorder

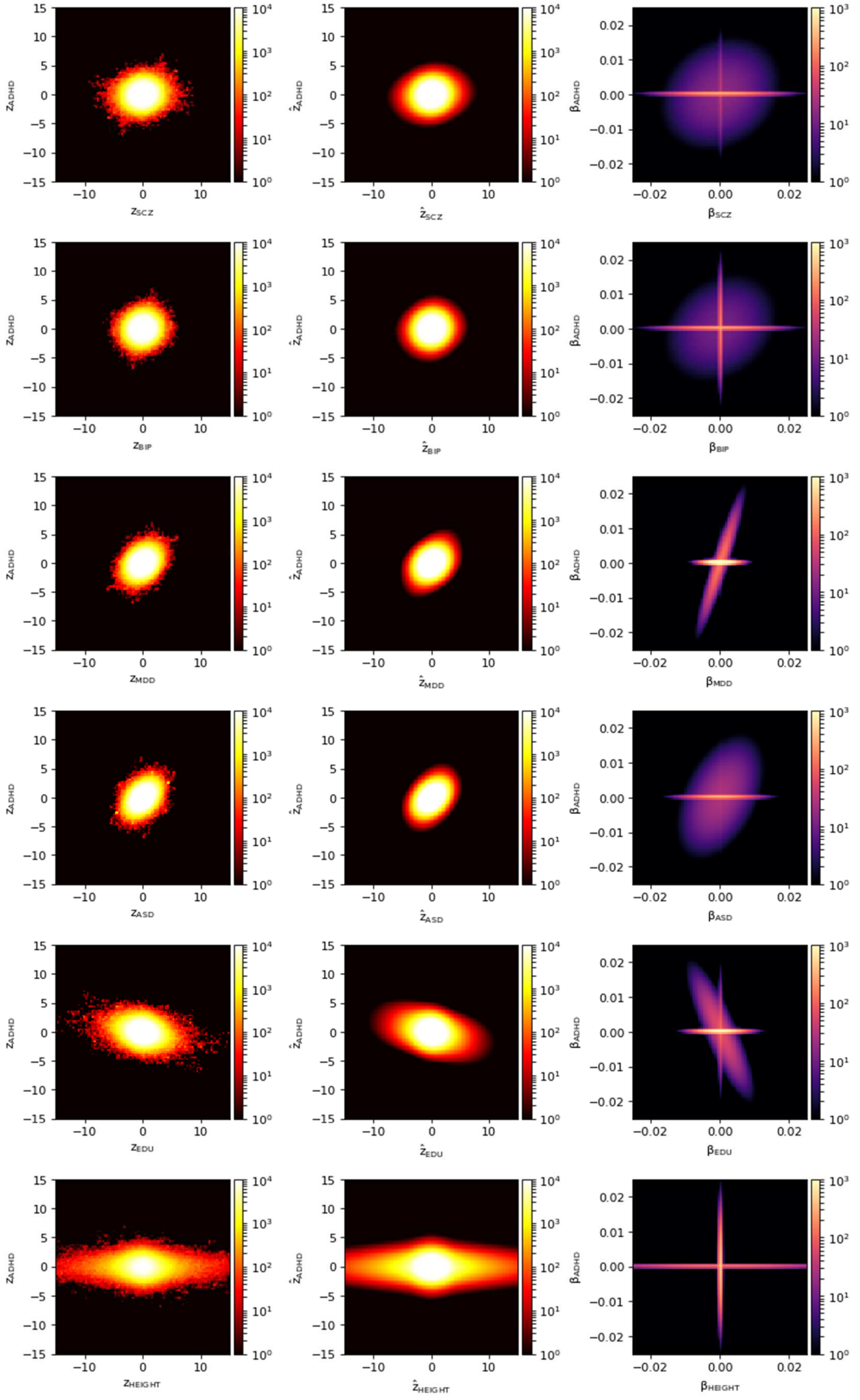

#### 14f. Observed and predicted bivariate density of GWAS association statistics, educational attainment

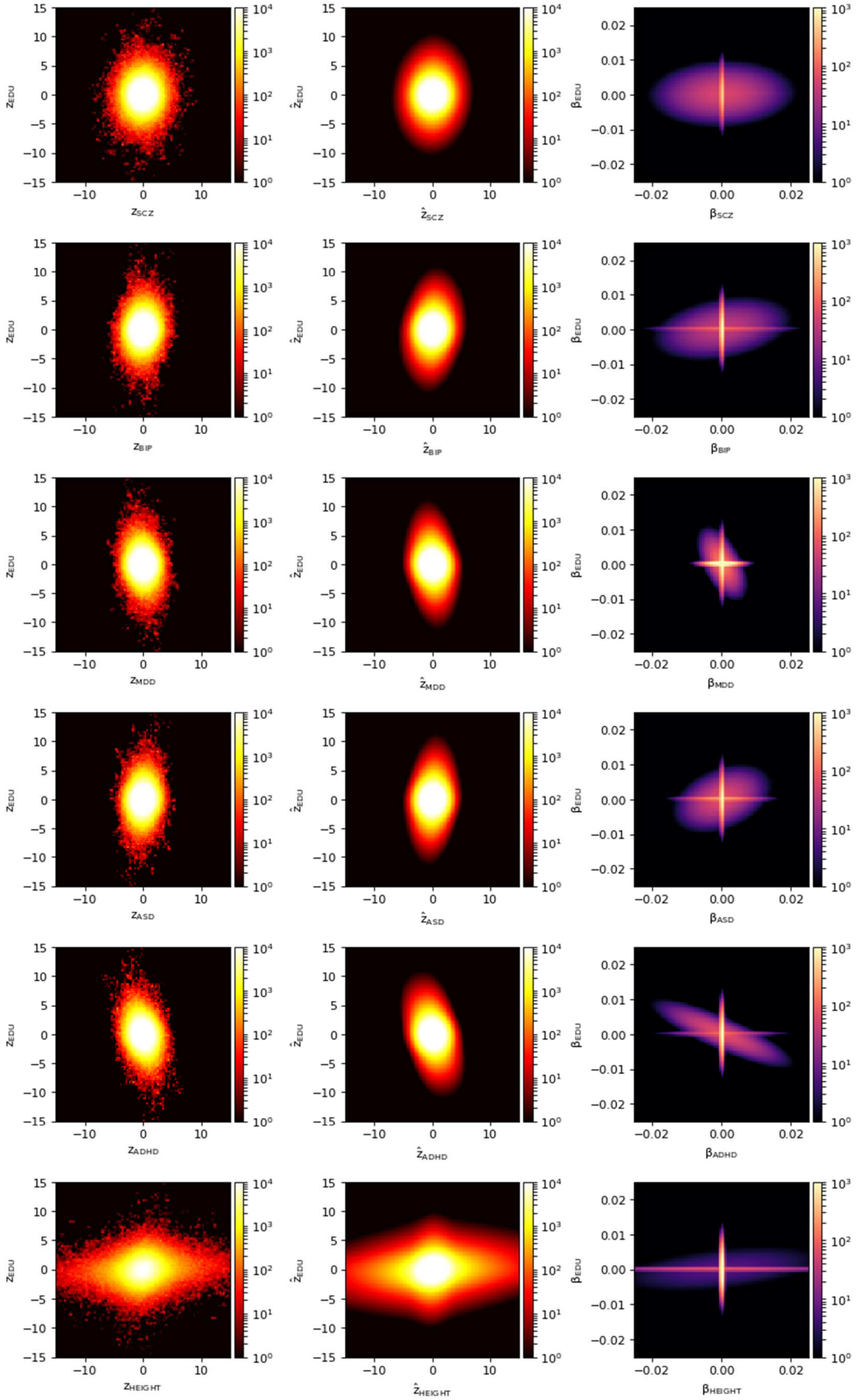

#### 14g. Observed and predicted bivariate density of GWAS association statistics, height

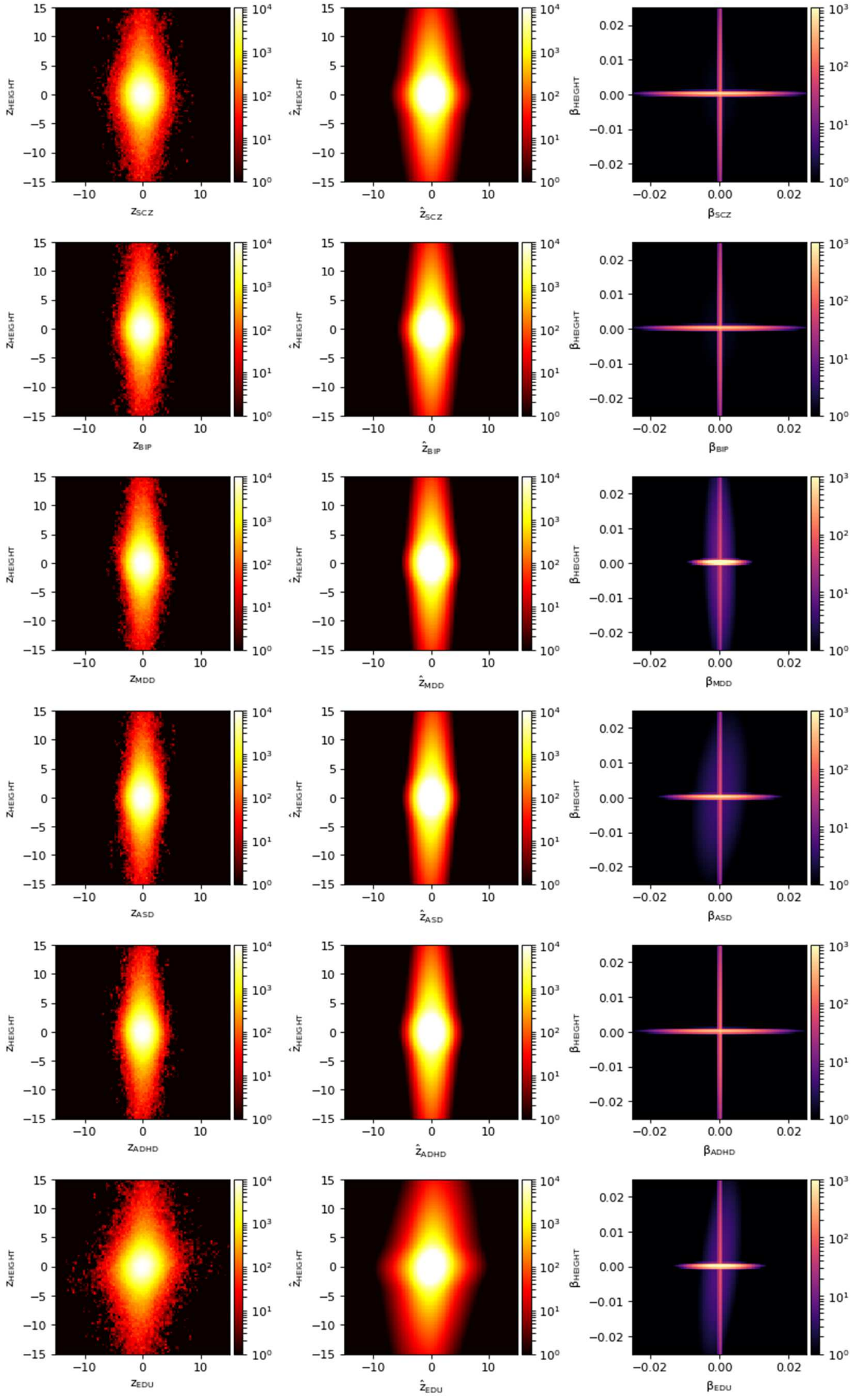

#### 15. Projected power plot for future GWAS sample sizes

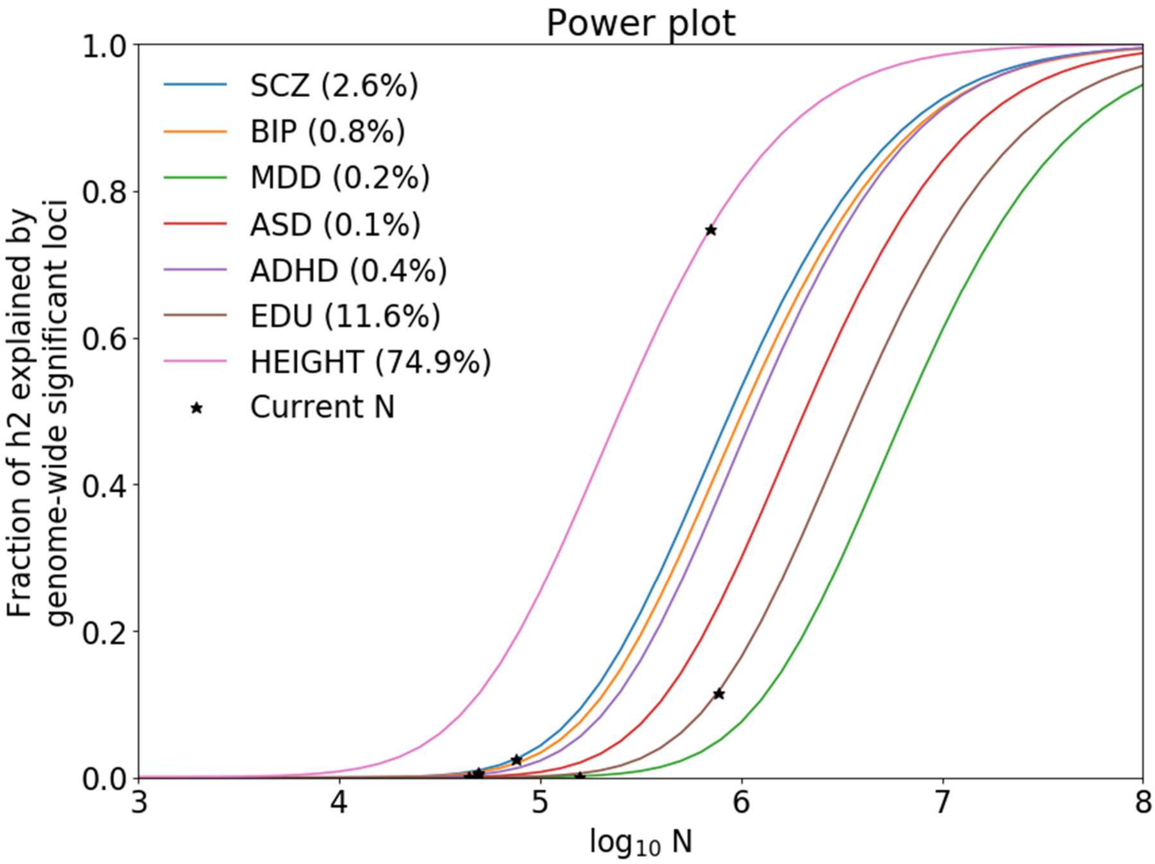


Proportion of SNP-heritability, captured by genome-wide significant SNPs, projected to the future GWAS sample size, N. Values for current GWAS sample sizes are shown in parentheses.

#### 16. Univariate Q-Q plots

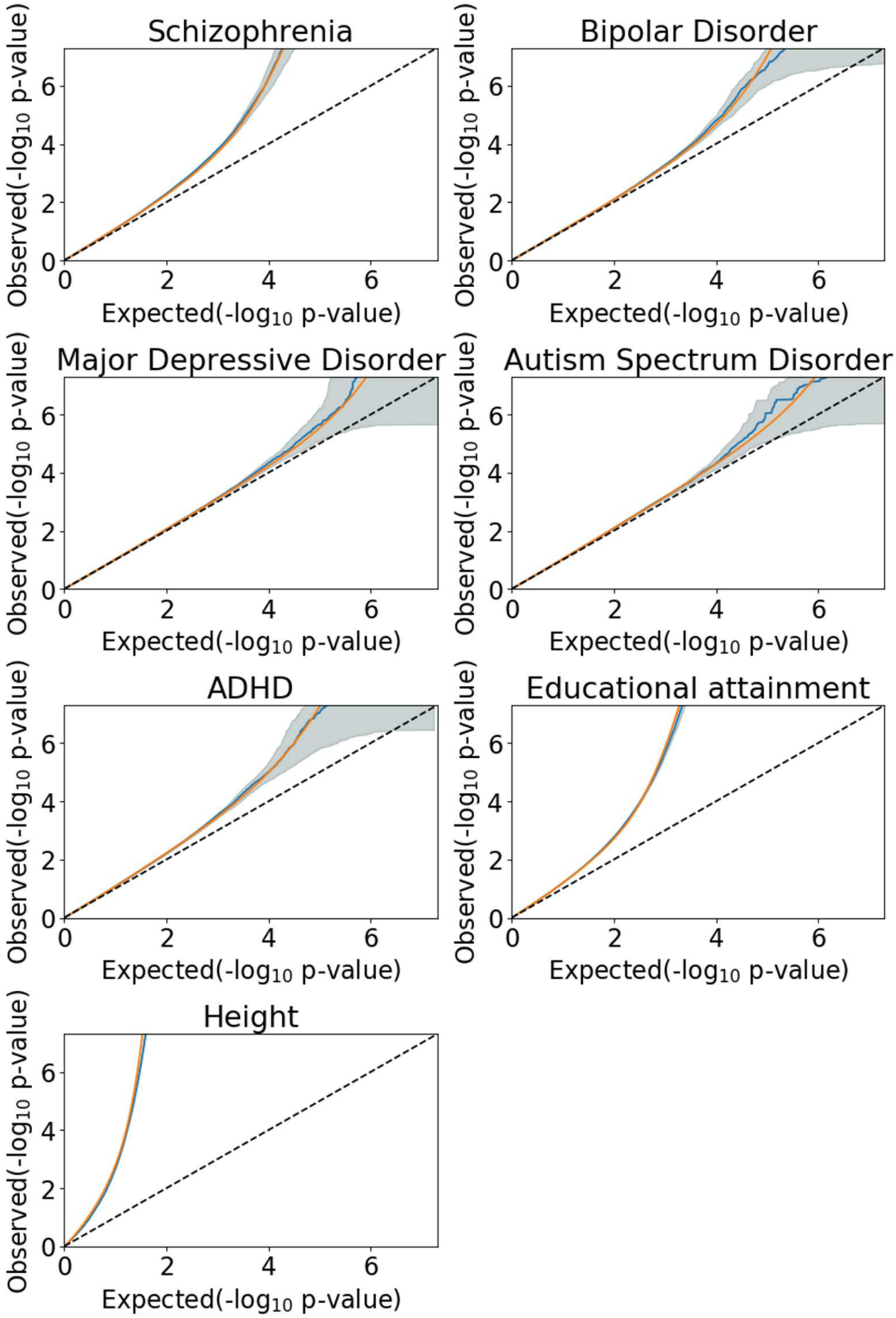


QQ plots for observed GWAS p-values (in blue) and model prediction (in orange). The dashed line is the expected QQ plot under null (no SNPs associated with the phenotype). The vertical axis is limited to the standard GWAS threshold of $p<5\times10^{-8}$, to highlight behavior of polygenic component. Points on the QQ plot are weighted according to LD structure, using n=64 iterations of random pruning at LD threshold r2=0.1. QQ plots are calculated on the entire set SNPs available in GWAS, constrained to LD Score Regression reference panel of 9.997.231 SNPs.

#### 17a. Q-Q plots of SNPs partitioned by MAF and LD score, schizophrenia

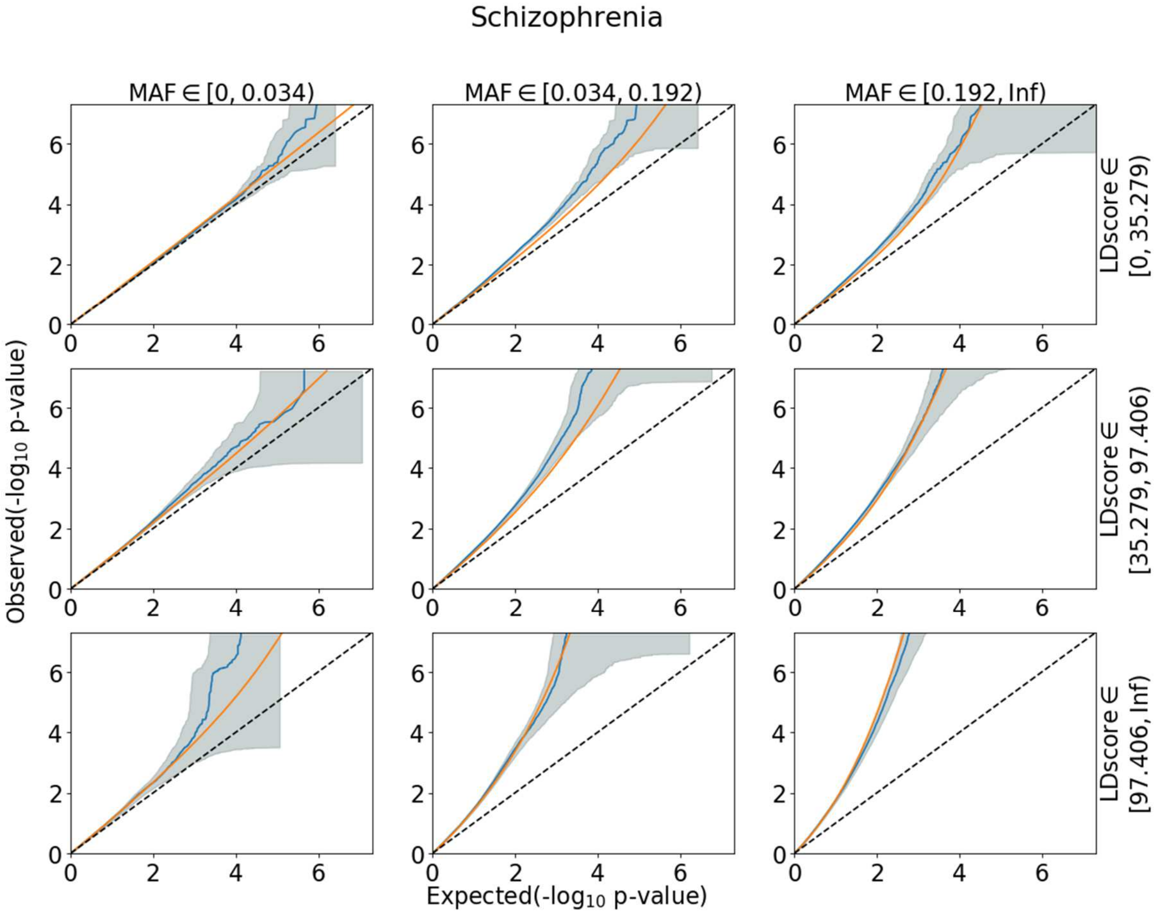


QQ plots for subsets of SNPs, showing observed GWAS p-values (in blue) and model prediction (in orange). All SNPs were partitioned into 9 groups according to minor allele frequency (MAF) and total LD score. The model was fit only once, so that all model predictions are based on the same set of parameters. Observed QQ plots show a stronger GWAS signal for SNPs with to higher MAF and higher LD score. Model's prediction follows the same pattern, indicating that model correctly captures dependency of GWAS association statistics on MAF and Total LD score.

#### 17b. Q-Q plots of SNPs partitioned by MAF and LD score, bipolar disorder

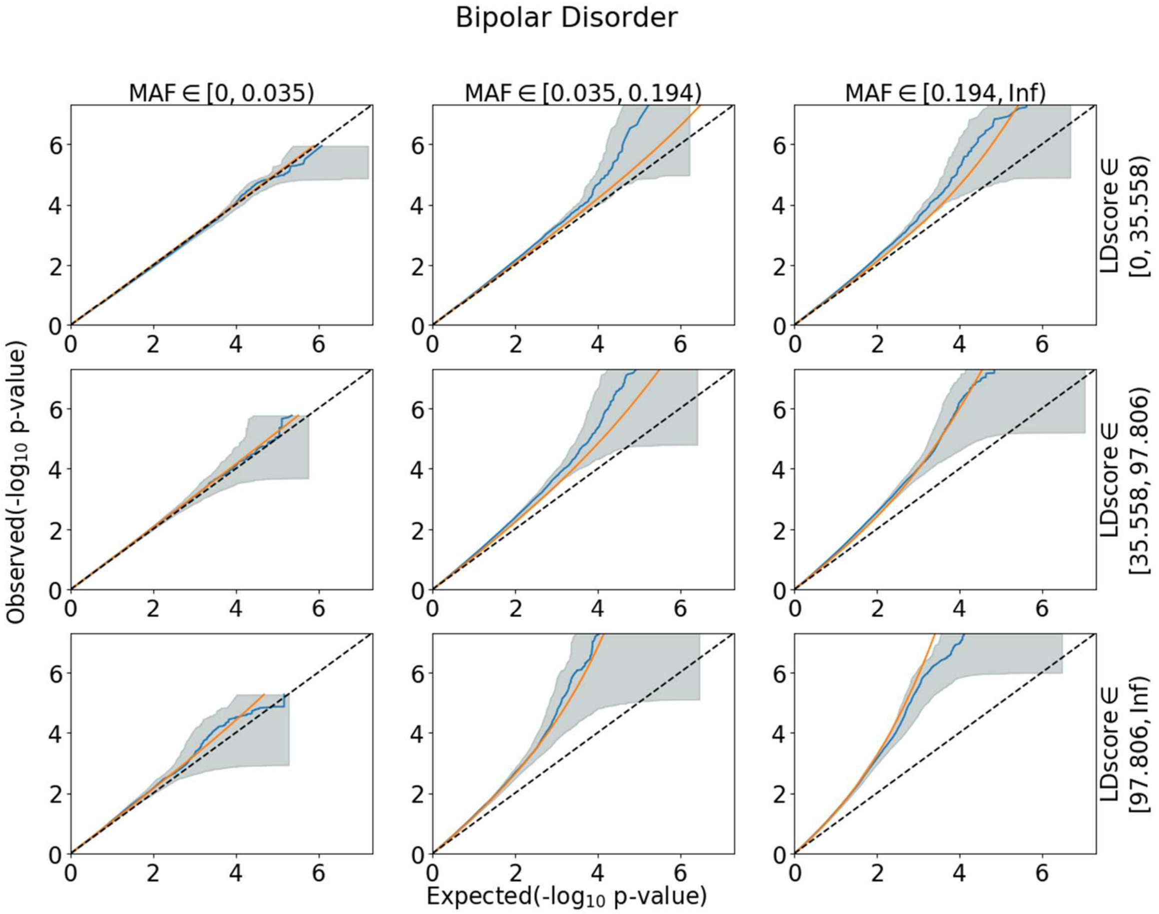


QQ plots of SNPs partitioned by MAF and LD score. Appearance of the Q-Q plot is as described on the previous figure.

#### 17c. Q-Q plots of SNPs partitioned by MAF and LD score, major depressive disorder

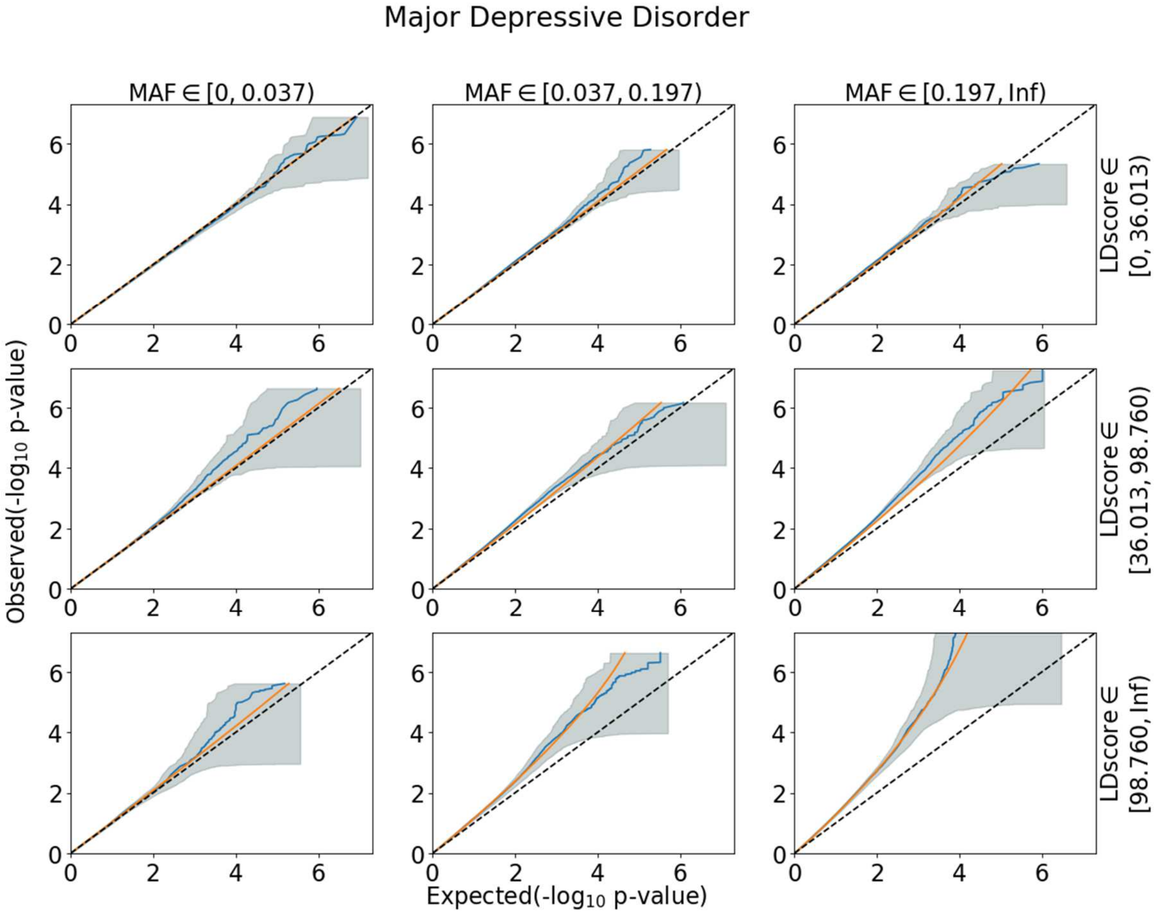


QQ plots of SNPs partitioned by MAF and LD score. Appearance of the Q-Q plot is as described on the previous figure.

#### 17d. Q-Q plots of SNPs partitioned by MAF and LD score, autism spectrum disorder

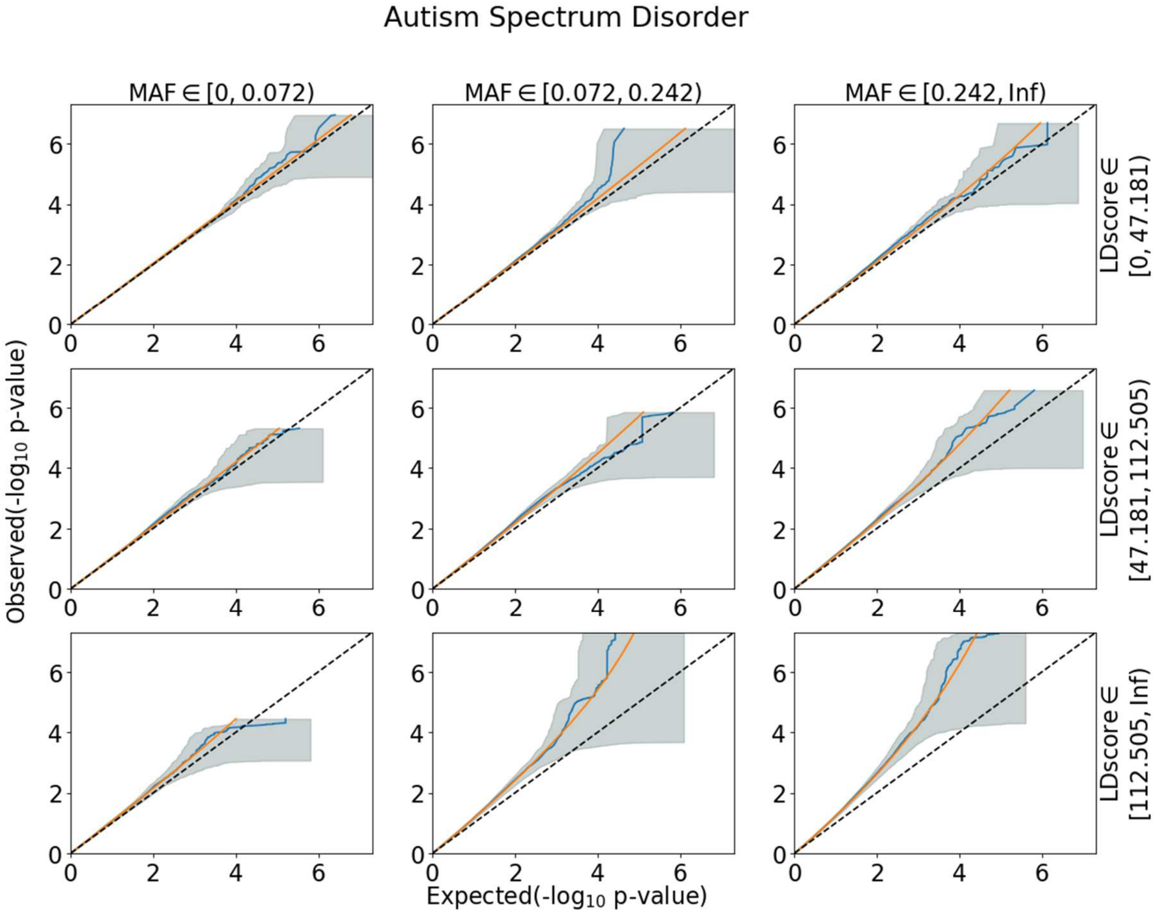


QQ plots of SNPs partitioned by MAF and LD score. Appearance of the Q-Q plot is as described on the previous figure.

#### 17e. Q-Q plots of SNPs partitioned by MAF and LD score, ADHD

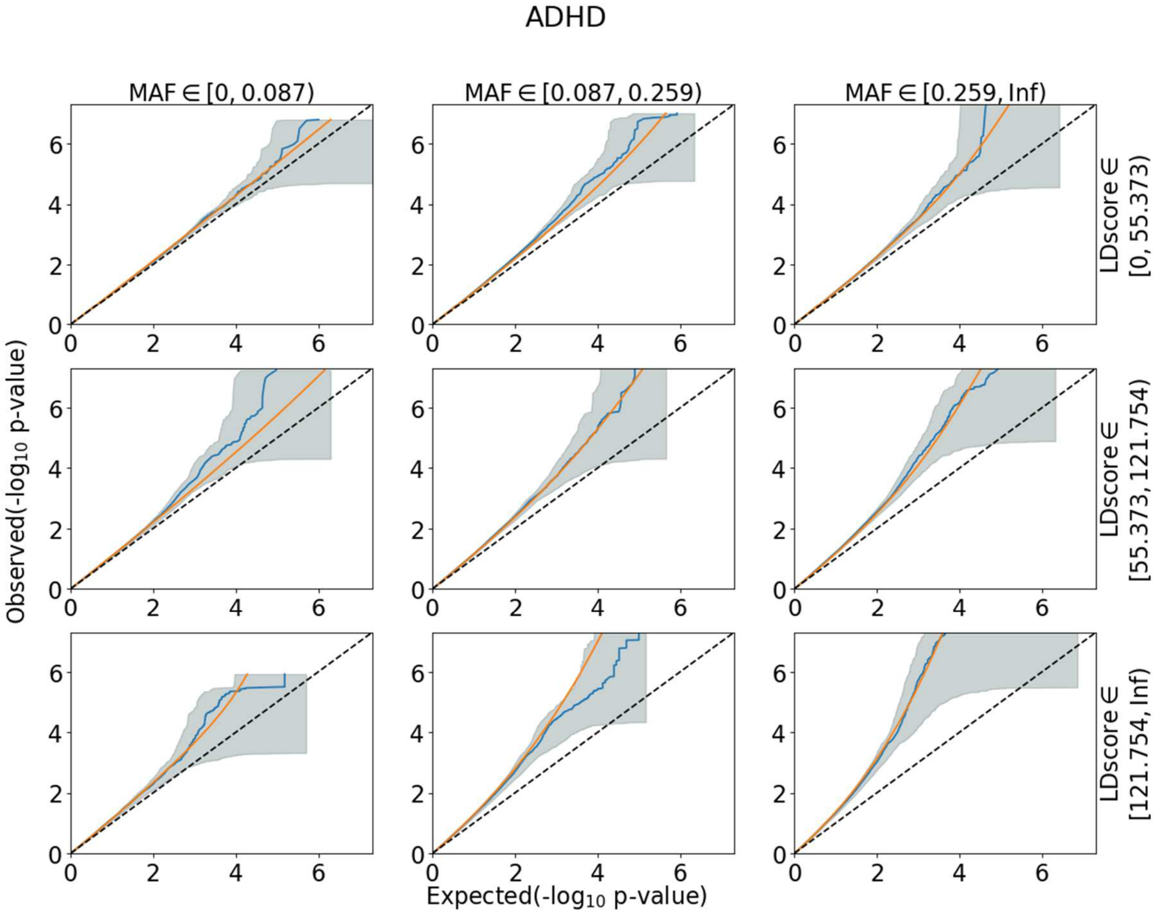


QQ plots of SNPs partitioned by MAF and LD score. Appearance of the Q-Q plot is as described on the previous figure.

#### 17f. Q-Q plots of SNPs partitioned by MAF and LD score, educational attainment

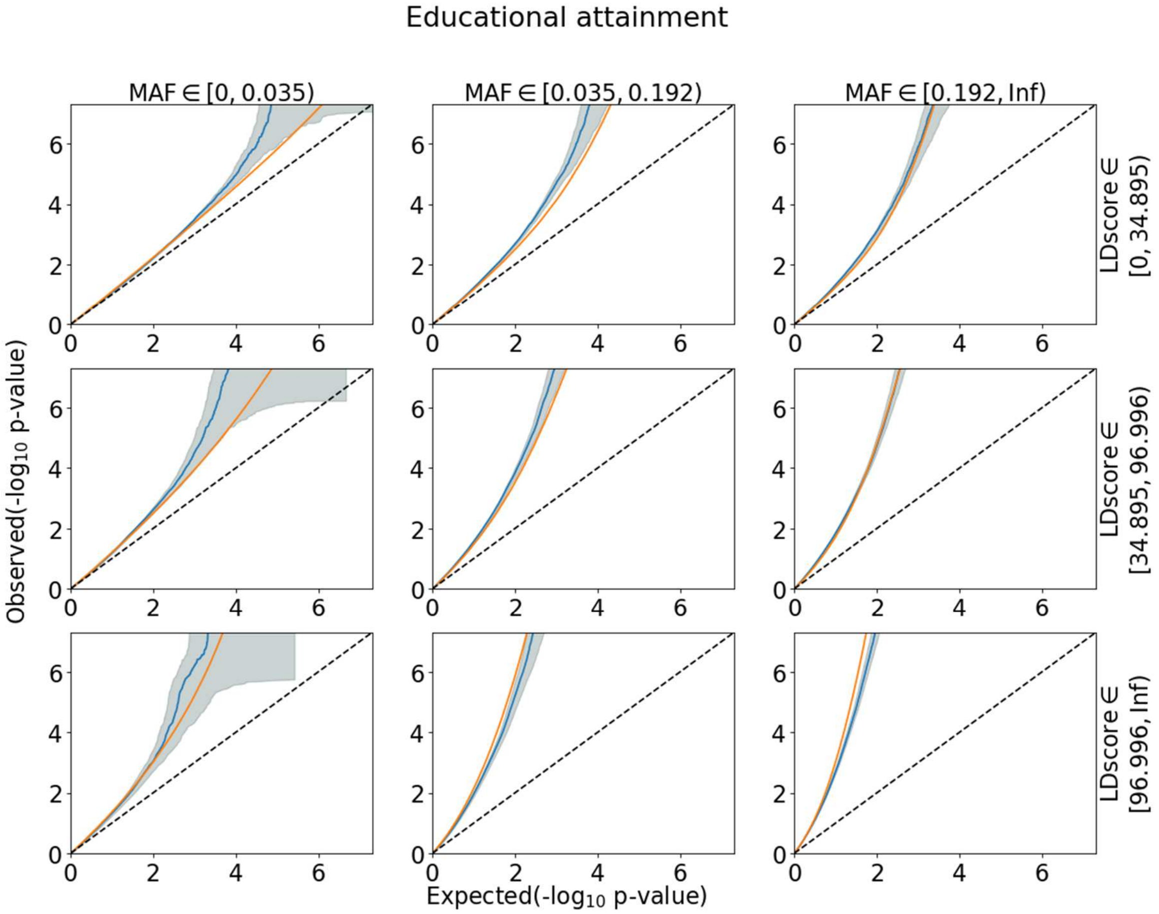


QQ plots of SNPs partitioned by MAF and LD score. Appearance of the Q-Q plot is as described on the previous figure.

#### 17g. Q-Q plots of SNPs partitioned by MAF and LD score, height

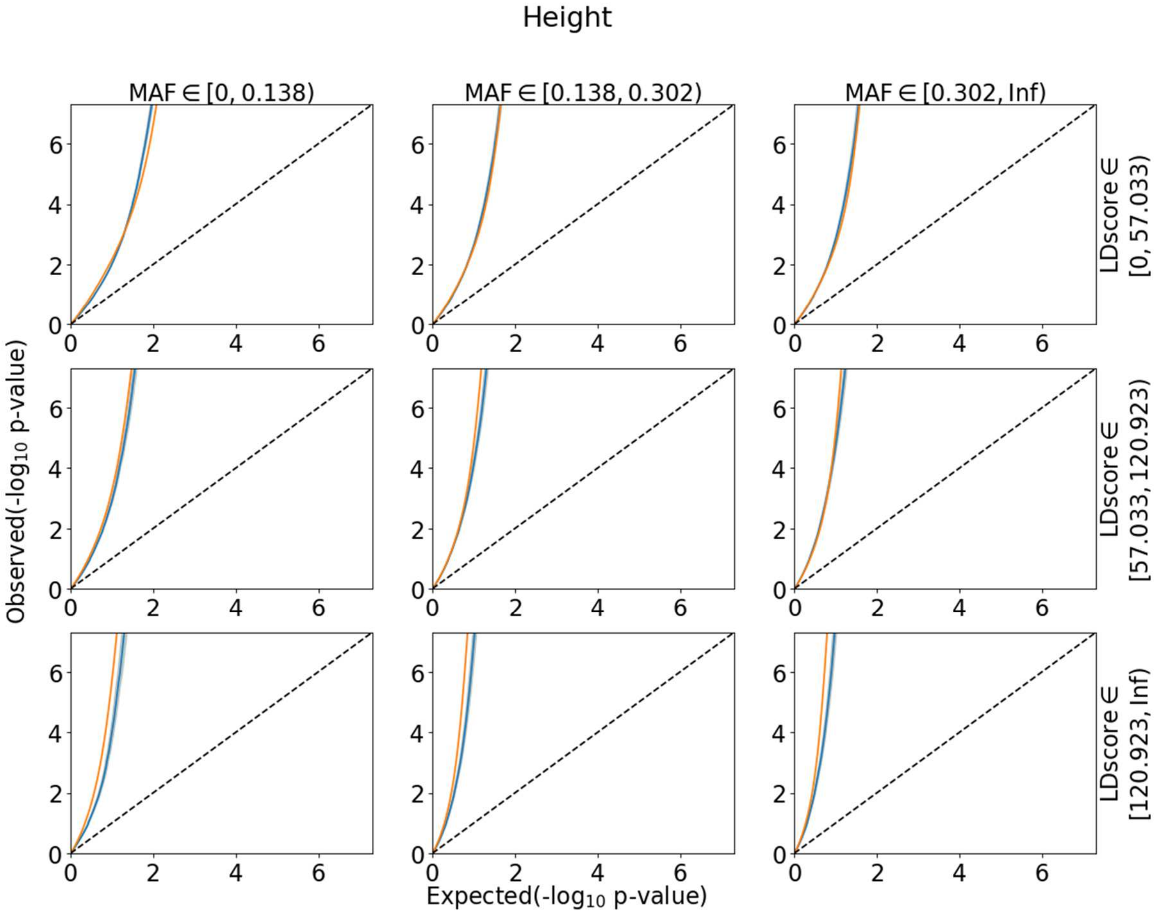


QQ plots of SNPs partitioned by MAF and LD score. Appearance of the Q-Q plot is as described on the previous figure.

### 18. Univariate likelihood as a function of polygenicity parameter

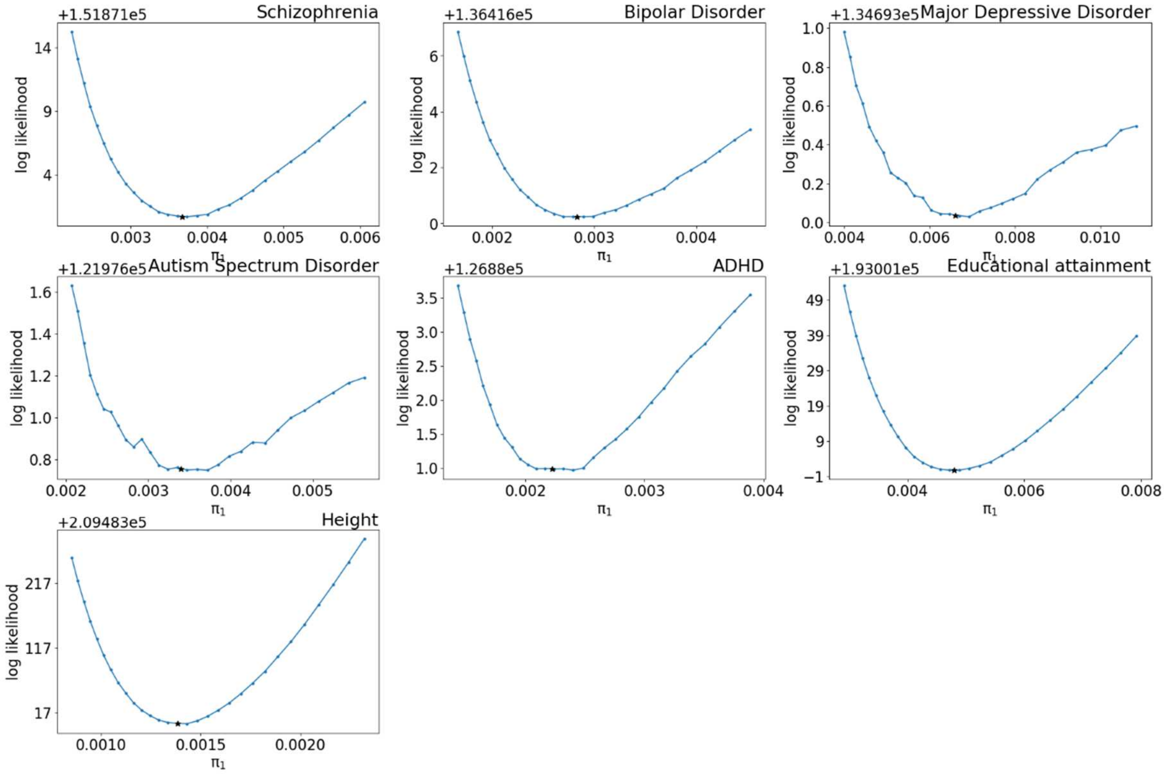


Log-likelihood of the univariate fit as a function of $\pi_1$ parameter. The remaining parameters of the model were constrained to the fitted values of heritability (h2) and variance distortion ($\sigma_0^2$). Asterix indicates fitted value of $\pi_1$ parameter.

#### 19. Bivariate likelihood as a function of polygenic overlap parameter

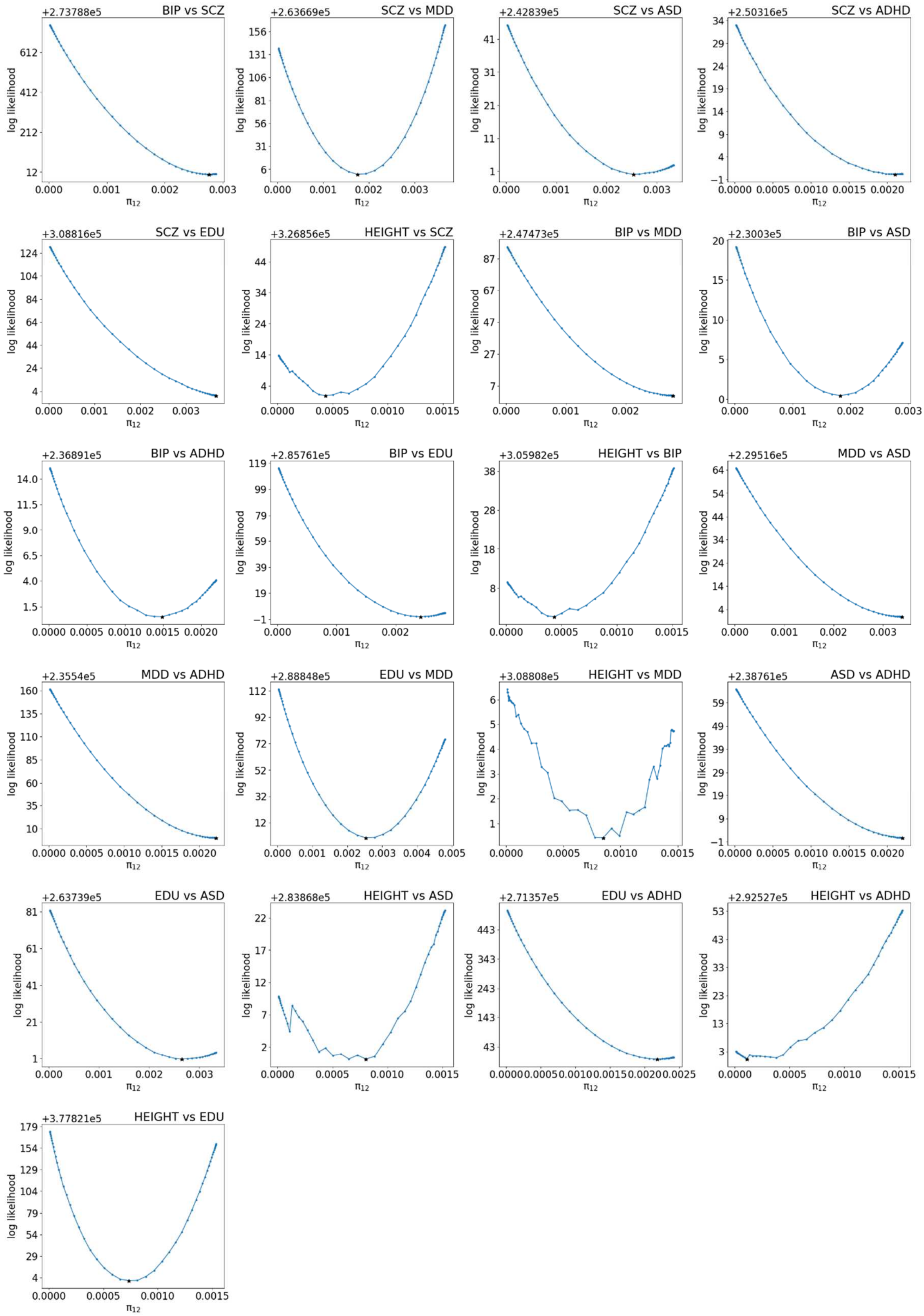


Log-likelihood of the bivariate fit as a function of $\pi_{12}$ parameter. The remaining parameters of the model were constrained to their fitted values. Asterix indicates fitted value of $\pi_{12}$ parameter.
