## Supplementary material for "Bivariate causal mixture model quantifies polygenic overlap between complex traits beyond genetic correlation"

### SUPPLEMENTARY NOTE

**Theory.** Consider a bi-allelic genetic variant $j$, and let $\beta_j$ be the effect size of allele substitution of that variant on a given quantitative trait. Variants with non-zero effect $\beta_j \neq 0$ are said to be *causal* for the trait. We assume a simple additive generative model

$$y_k = \sum_{j=1}^{M} g_{kj}\beta_j + e_k, \quad (1)$$

where $y_k$ is a quantitative trait measured on $k$-th individual ($k = 1, \ldots, N$), $g_{kj}$ is an additively coded number of reference alleles for $j$-th variant ($j = 1, \ldots, M$) on $k$-th individual, $\beta_j$ is the effect size of variant $j$, and $e_k$ contains additive environmental and measurement error effects. In a vector form, $\mathbf{y} = \mathbf{G}\beta + \mathbf{e}$, and we denote rows of the genotype matrix $\mathbf{G}$ as $\mathbf{x_k^T}$ (genotype vector for $k$-th individual) and columns as $\mathbf{v_j}$ (genotype vector for $j$-th variant).

The scales of $\beta_j$ and $e_k$ are chosen so that phenotype vector $\mathbf{y}$ has unit variance, $\mathrm{Var}(\mathbf{y}) = 1$, and $\mathrm{Var}(\mathbf{e}) = 1 - h^2$ where $h^2$ is narrow-sense heritability of the trait. Genotypes $g_{kj}$ are assumed to be centered for each variant to have zero mean across individuals, but are not normalized, therefore $\mathrm{Var}(\mathbf{v_j}) = 2p_j(1-p_j)$, where $p_j$ is minor allele frequency of $j$-th variant.

**Theorem 1.** *Let $\hat{\beta}'_j$ be GWAS estimate of $j$-th effect size, assessed via univariate linear regression, and $z_j$ be corresponding z-score, $z_j = \hat{\beta}'_j/\hat{se}(\beta'_j)$. Then*

$$\begin{aligned} z_j &= \delta_j + \epsilon_j, \\ \delta_j &= \sqrt{N_j}\sum_i \sqrt{H_i} r_{ij}\beta_i, \\ \epsilon_j &\sim \mathcal{N}(0, \sigma_0^2), \end{aligned} \quad (2)$$

*where $N_j$ is the number of subjects with non-missing genotype information on $j$-th variant; $H_i = 2p_i(1-p_i)$ is heterozygosity of $i$-th variant; $r_{ij} = corr(\mathbf{v_i}, \mathbf{v_j})$ is an allelic correlation coefficient (LD r2) between genotypes of variants $i$ and $j$; summation $\sum_i$ runs across all variants $i$ with non-zero $r_{ij}$ with $j$-th variant, and parameter $\sigma_0^2$ accounts for cryptic relatedness among individuals.*

In the absence of covariates, the least square estimates can be expressed as

$$\hat{\beta}'_j = \frac{\mathbf{v_j^T y}}{\mathbf{v_j^T v_j}} = \beta_j + \sum_{i \neq j} \hat{\xi}_{ij}\beta_i + \frac{\mathbf{v_j^T e}}{\mathbf{v_j^T v_j}},$$

where $\hat{\xi}_{ij} = \mathbf{v_i^T v_j}/\mathbf{v_j^T v_j} = \hat{\zeta}_{ij}/\hat{\zeta}_{jj}$, with $\hat{\zeta}_{ij} = \mathbf{v_i^T v_j}/N$ being an estimate of the covariance between $i$-th and $j$-th variants:

$$\hat{\zeta}_{ij} \simeq \sqrt{2p_i(1-p_i)}\sqrt{2p_j(1-p_j)}r_{ij}.$$

Here the symbol "$\simeq$" denotes asymptotic equality as $n \to \infty$. Then using $\hat{se}(\beta'_j) = \sqrt{H_j/N_j}$ result in (2). ■

**Univariate Causal Mixture model for GWAS.** In unvariate case we model $\beta_j$ as a mixture of null and non-null components:

$$\beta_j \sim (1-\pi_1)\mathcal{N}(0,0) + \pi_1\mathcal{N}(0,\sigma^2_\beta), \tag{3}$$

where $\pi_1$ is the proportion of causal variants, $\sigma^2_\beta$ is the "discoverability" (phenotypic variance explained per causal variant), and $\mathcal{N}(0,0)$ is a Dirac delta function with the entire probability mass concentrated as 0. All parameters are assumed to be equal across all variants. We are interested in finding parameters $\pi_1$, $\sigma^2_\beta$ and $\sigma^2_0$ that best describe the observed values of $z_j$ in GWAS summary statistics. For this we need likelihood term $pdf(z_j|\pi_1,\sigma^2_\beta,\sigma^2_0)$, where $z_j$ is given by (2) with $\beta_j$ distributed according to (3):

$$z_j|\pi_1,\sigma^2_\beta,\sigma^2_0 \sim \sum_i \Big[(1-\pi_1)\mathcal{N}(0,0) + \pi_1\mathcal{N}(0, N_jH_ir^2_{ij}\sigma^2_\beta)\Big] + \mathcal{N}(0,\sigma^2_0), \tag{4}$$

where the outer sumation $\sum_i$ denotes a sum of random variables, then each summand is a mixture of two distributions, finally the last plus sign again indicates sum of random variables (e.i. convolution of corresponding probability density functions).

Below we focus on a computationally efficient way of calculating likelihood from expression (4). This task is not straighforward because (4) involves multiple convolutions of a mixture distribution. First we simplify expression (4) by assuming that regardless of index $i$ all $r^2_{ij}$ values are equal to a certain parameter $\kappa^2_j$, and also assuming equal heterozigosity of all variants (for all $i$, $H_i = H_j$). Then we relax this assumption and derive general formulas for 2nd and 4th raw moments of $z_j|\pi_1,\sigma^2_\beta,\sigma^2_0$. Finally, we derive two computationally efficient approximations, one based on two-component Gaussian mixture, and another based on Poisson approximation to binomial distribution. In the Online methods, the two-component Gaussian mixture is referred to as "fast" model, and it is used to perform initial search in the space of models parameters. The Poisson approximation is not used directly, but it provides an important insight into mathematics of GWAS z scores in a context of the casal mixture model.

**Lemma 2.** *For a given variant $j$ assume all non-zero allelic correlations $r^2_{ij}$ are equal to a certain value $\kappa^2_j$, and assume equal heterozigosity of all variants (for all $i$, $H_i = H_j$). Then*

$$pdf(z_j|\pi_1,\sigma^2_\beta,\sigma^2_0) = \sum_{k=0}^{L_j}\binom{L_j}{k}\pi_1^k(1-\pi_1)^{L_j-k}\phi(z_j;0,\sigma^2_0 + k\kappa^2_jH_jN_j\sigma^2_\beta), \tag{5}$$

*where $\ell_j = \sum_i r^2_{ij}$ indicates total LD score of variant $j$, and $L_j = \ell_j/\kappa^2_j$ denote the number of variants in LD with variant $j$.*

*Proof.* Consider two-component mixture of random variables $A$ and $B$ with weights $p$ and $q$ (where $p+q=1$). $L$-times convolution of this mixture with itself is given by the following combinatorial expression:

$$\sum_{i=1}^{L}(pA+qB) = \sum_{k=0}^{L}\binom{L}{k}p^kq^{L-k}\Big(kA + (L-k)B\Big).$$

Applying this formula to (4) with $p=\pi_1$, $q=1-\pi_1$, $A=\mathcal{N}(0,H_jN_j\sigma^2_\beta)$ and $B=\mathcal{N}(0,0)$ concludes the proof. ∎

With $\pi_1 = 1$ expression (5) simplifies to $\mathcal{N}(0, \sigma_0^2 + \ell_j H_j N_j \sigma_\beta^2)$, which corresponds to the "infinitesimal" model underlying LD score regression.

Now we aim to relax the assumptions of $r_{ij}^2 = \kappa_j^2$ and $H_i = H_j$, and, for the general case, derive expressions for the 2nd and 4th raw moments $E(z_j^2)$ and $E(z_j^4)$. First we show how they are related to the raw moments of $\delta_j$.

**Lemma 3.**

$$E(z_j^2) = E(\delta_j^2) + \sigma_0^2,$$
$$E(z_j^4) - 3(E(z_j^2))^2 = \Big(E(\delta_j^4) - 3\big(E(\delta_j^2)\big)^2\Big).$$

*Proof.* Expression for $E(z_j^2)$ straighforwardly follows from (2). To prove the second formula we note that for two independent random variables $x$ and $y$ the expectation $E(x+y)^4 = Ex^4+3Ex^3Ey+6Ex^2Ey^2+3ExEy^3+Ey^4$, which simplifies to $E(x+y)^4 = Ex^4 + 6Ex^2Ey^2 + Ey^4$ for zero mean random variables. Thus

$$E(z_j^4) = E(\delta_j^4) + 6E(\delta_j^2)\sigma_0^2 + 3\sigma_0^4,$$

and hence $E(z_j^4) - 3(E(z_j^2))^2 = (N_j H_j)^2 \Big(E(\beta_j'^4) - 3\big(E(\beta_j'^2)\big)^2\Big)$. ∎

Last lemma allows us to simplify notation and derive all formulas in terms of $E(\delta_j^2)$ and $E(\delta_j^4)$, which depend only on $\pi_1$ and $\sigma_\beta^2$, but not on $\sigma_0^2$.

**Lemma 4.**

$$E(\delta_j^2) = N_j \ell_j \pi_1 \sigma_\beta^2,$$
$$E(\delta_j^4) - 3E(\delta_j^2)^2 = 3N_j^2 R_j \pi_1 \pi_0 \sigma_\beta^4, \tag{6}$$

*where $R_j = \sum_i H_i^2 r_{ij}^4$ is a sum of fourth power of allelic correlations (adjusted for heterozigosity), and $\ell_j = \sum_i H_i r_{ij}^2$ is total LD score (adjusted for heterozigosity).*

*Proof.* From (2), $\delta_j/N_j = \sum_i \sqrt{H_i} r_{ij} \beta_i$, where all $\beta_i$ are independent equally distributed random variables with $E(\beta_i^2) = \pi_1 \sigma_\beta^2$ and $E(\beta_i^4) = 3\pi_1 \sigma_\beta^4$.

$$E(\delta_j^2/N_j) = E\Big(\sum_i \sqrt{H_i} r_{ij} \beta_i\Big)^2 = \sum_i H_i r_{ij}^2 E(\beta_i^2) + \sum_{i<k} 2\sqrt{H_i H_k} r_{ij} r_{kj} E(\beta_i) E(\beta_k).$$

The second sum is equal to zero because $E(\beta_i) = 0$. Thus,

$$E(\delta_j^2/N_j) = \sum_i H_i r_{ij}^2 E(\beta_i^2) = \ell_j E(\beta^2) = \ell_j \pi_1 \sigma_\beta^2.$$

Similarly, for $E(\delta_j^4/N_j^2)$ we get

$$E(\delta_j^4/N_j^2) = E\Big(\sum_i \sqrt{H_i} r_{ij} \beta_i\Big)^4 = \sum_i H_i^2 r_{ij}^4 E(\beta_i^4) + \sum_{i<k} 6 H_i H_k r_{ij}^2 r_{kj}^2 E(\beta_i^2) E(\beta_k^2),$$

where all summands with zero mean are already omited. Note that

$$2\sum_{i<k} H_i H_k r_{ij}^2 r_{kj}^2 = \Big(\sum_i H_i r_{ij}^2\Big)^2 - \sum_i H_i^2 r_{ij}^4 = \ell_j^2 - R_j,$$

hence $E(\delta_j^4) = N_j^2 R_j E(\beta^4) + 3N_j^2(\ell_j^2 - R_j)E(\beta^2)^2$, and thus

$$E(\delta_j^4) - 3E(\delta_j^2)^2 = N_j^2 R_j \Big( E(\beta^4) - 3N_j^2 E(\beta^2)^2 \Big) = 3N_j^2 R_j (\pi_1 - \pi_1^2)\sigma_\beta^4.$$

■

**Corollary 4.1.** *An arbitrary LD structure of a $j$-th variant could be approximated via "spike"-like histogram of equal $r_{ij}$ values, with effective allelic correlation $\hat{\kappa}_j^2 = R_j/\ell_j$, and effective block size $\hat{L}_j = \ell_j^2/R_j$, and such approximation preserves 2nd and 4th moments of $z_j$.*

*Proof.* The statement directly follows from lemmas 3 and 4. ■

Last corolary also imply that the variance of $z_j$ depends only on the total LD score $\ell_j$, and does not depend on $\kappa_j^2$ parameter. On contrary, the excess kurtosis linearly depend on $\kappa_j^2$. Recall that the excess kurtosis is a measure of how heavy are the tails of the distribution. This means that in the context of a mixture model total LD score itself is not sufficient to describe the distribution of $z$ scores, because variants with higher $\kappa_j^2$ will tend to have larger $z$ scores despite having the same total LD score.

Last corollary alloes one to approximate likelihood $pdf(z|\pi_1, \sigma_\beta^2, \sigma_0^2)$ with binomial formula (5), with parameters $\kappa_j = R_j/\ell_j$, $H_j = 1$, and $L_j = \ell_j^2/R_j$. Such approximation is reasonable in a sense that it preserves 2nd and 4th moments of $z_j$.

Likelihood calculation from (5) still involves a potentially large sum $\sum_{k=0}^{L}$. To further speedup calculations we show that a two-component mixture of Gaussin distributions has enough flexibility to preserve 2nd and 4th moments of $z_j$.

**Theorem 5** (Gaussian approximation)**.** *Let*

$$z'_j \sim \pi'_{0j}\mathcal{N}(0, \sigma_0^2) + \pi'_{1j}\mathcal{N}(0, \sigma_0^2 + \sigma_{\delta_j}^2), \tag{7}$$

*where*

$$\begin{aligned} \pi'_{1j} &= \frac{\pi_1 \ell_j}{\pi_1 \ell_j + \pi_0 \kappa_j^2}, \\ \sigma_{\delta_j}^2 &= N_j \sigma_\beta^2 (\pi_0 \kappa_j^2 + \pi_1 \ell_j), \\ \ell_j &= \sum_i H_i r_{ij}^2, \quad R_j = \sum_i H_i^2 r_{ij}^4, \\ \kappa_j^2 &= R_j/\ell_j, \quad \pi_0 = 1 - \pi_1, \quad \pi'_{0j} = 1 - \pi'_{1j}. \end{aligned} \tag{8}$$

*Then 2nd and 4th moments of $z'_j$ and $z_j$ distributions are equal.*

*Proof.* Lemmas 3 and 4 derive expressions for 2nd and 4th moments of $z_j$. What's left is to find same moments for $z'_j$, defined by (7):

$$\begin{aligned} E(z_j'^2) &= \pi'_{1j}\sigma_{\delta_j}^2 + \sigma_0^2, \\ E(z_j'^4) - 3(E(z_j'^2))^2 &= \pi'_{1j}\pi'_{0j}\sigma_{\delta_j}^4, \end{aligned}$$

Using (8) and doing straighforward algebraic computations concludes the proof. ■

Gaussian approximation provides very efficient way of calculating likelihood $pdf(z_j|\pi_1, \sigma_\beta^2, \sigma_0^2)$. As a downside, such two-component gaussian mixture does not fully capture heavy tails of $z_j$ distribution — even though both 2nd and 4th moments are preserved. The following approximation, which we call *poisson-type approximation*, gives more accurate representation for the tails of z-score distribution. A key feature of poisson-type approximation is that it has the same underlying structure as the binomial formula (5), e.i. it is a mixture of gaussians with linearly increasing variance ($k\sigma_j^2$, $k = 0, 1, 2, \ldots,$).

**Theorem 6** (Poisson-type approximation)**.** *Let*

$$z''_j \sim \sum_{k=0}^{\infty} \frac{\lambda_j^k e^{-\lambda_j}}{k!} \mathcal{N}(0, \sigma_0^2 + k\sigma_j^2), \tag{9}$$

*where*

$$\begin{aligned} \lambda_j &= (\pi_1 \ell_j^2)/(\pi_0 R_j), \\ \sigma_j^2 &= N_j \sigma_\beta^2 \pi_0 R_j / \ell_j. \end{aligned} \tag{10}$$

*Then 2nd and 4th moments of $z''_j$ and $z_j$ distributions are equal.*

*Proof.* Lemmas 3 and 4 derive expressions for 2nd and 4th moments of $z_j$. What's left is to find same moments for $z''_j$. Let's denote $p_k = \frac{\lambda_j^k e^{-\lambda_j}}{k!}$. Then,

$$E(z''^2) = \sigma_0^2 + \sigma_j^2 \sum_{k=0}^{\infty} k p_k = \sigma_0^2 + \sigma_j^2 \lambda_i.$$

$$E(z''^4) - (E(z''^2))^2 = 3\sigma_0^4 + 3\sigma_j^4 \Big( \sum_{k=0}^{\infty} k^2 p_k - \Big( \sum_{k=0}^{\infty} k p_k \Big)^2 \Big) = 3\sigma_j^4 \lambda_j.$$

Using (10) and doing straighforward algebraic computations concludes the proof. ■

In practice the invfinite sum in (9) converges very quickly, typically $k_{max} = 5$ is sufficient. In addition, $\sigma_j^2$ can be adjusted by $poisscdf(k_{max-1}, \lambda)$ factor, where *poisscdf* gives cumulated distribution function of the discrete Poisson distribution. Such adjustement compensates for variance explained by $k > k_{max}$.

To get an improved fit of the model to the data one may split LD structure of $j$-th variant into regions with low $r_{ij}^2$ and high $r_{ij}^2$, each approximated with own $(\lambda_j, \sigma_j^2)$ values, This can be handled using the following analytical way of convolving two Poisson-type approximations:

$$z''_j \sim \sum_{k_1=0}^{\infty} \sum_{k_2=0}^{\infty} \frac{\lambda_{1j}^{k_1} e^{-\lambda_{1j}}}{k_1!} \frac{\lambda_{2j}^{k_2} e^{-\lambda_{2j}}}{k_2!} \mathcal{N}(0, \sigma_0^2 + (k_1 \sigma_{1j}^2 + k_2 \sigma_{2j}^2)).$$

This formulas simplify considerably if $\sigma_{1j}^2 = \sigma_{2j}^2 = \sigma_j^2$. If that's the case, we have

$$z''_j \sim \sum_k (f * g)_k \mathcal{N}(0, \sigma_0^2 + k\sigma_j^2), \tag{11}$$

where $f_k = \frac{\lambda_{1j}^{k_1} e^{-\lambda_{1j}}}{k_1!}$, $g_k = \frac{\lambda_{2j}^{k_2} e^{-\lambda_{2j}}}{k_2!}$, and $(f * g)_k = \sum_m f_m g_{k-m}$ denotes discrete confolution of coefficients $f_k$ and $g_k$.

**Bivariate causal mixture model for GWAS.** Bivariate model is defined as follows:

$$
\begin{aligned}
&(\beta_{1j}, \beta_{2j}) \sim \pi_0 \mathcal{N}(0,0) + \pi_1 \mathcal{N}(0,\Sigma_1) + \pi_2 \mathcal{N}(0,\Sigma_2) + \pi_{12}\mathcal{N}(0,\Sigma_{12}),\\
&\Sigma_1 = \begin{bmatrix} \sigma_1^2 & 0 \\ 0 & 0 \end{bmatrix}, \Sigma_2 = \begin{bmatrix} 0 & 0 \\ 0 & \sigma_2^2 \end{bmatrix}, \text{ and } \Sigma_{12} = \begin{bmatrix} \sigma_1^2 & \rho_{12}\sigma_1\sigma_2 \\ \rho_{12}\sigma_1\sigma_2 & \sigma_2^2 \end{bmatrix},\\
&(z_{1j}, z_{2j}) \sim (\delta_{1j}, \delta_{2j}) + \mathcal{N}\Big(0, \Sigma_0\Big), \;\; \delta_{\cdot j} = \sqrt{N_{\cdot j}} \sum_i \sqrt{H_i} r_{ij} \beta_{\cdot i},\\
&\Sigma_0 = \begin{bmatrix} \sigma_{01}^2 & \rho_0\sigma_{01}\sigma_{02} \\ \rho_0\sigma_{01}\sigma_{02} & \sigma_{02}^2 \end{bmatrix}.
\end{aligned} \tag{12}
$$

We denote the vector of nine parameters of the model by $\theta = (\pi_1, \pi_1, \pi_{12}, \sigma_1^2, \sigma_2^2, \rho_{12}, \sigma_{01}^2, \sigma_{02}^2, \rho_0)$. Below we derive a computationally tractable approximation for the likelihood term $pdf(z_{1j}, z_{2j}|\theta)$, which allows us to fit estimated parameters $\hat{\theta}$ from GWAS summary statistics $(z_{1j}, z_{2j})$ by maximizing log-likelihood, weighted by $w_j = 1/\ell_j$ or by weights based on random pruning to avoid over-counting in large LD blocks:

$$
F(\theta) = \sum_j w_j \log pdf(z_{1j}, z_{2j}|\theta) \to \max_\theta .
$$

First, we assume that parameters $\pi_1$, $\pi_2$ and $\pi_{12}$ are small ($\pi_1 \ll 1$, $\pi_2 \ll 1$, $\pi_{12} \ll 1$). Therefore $(\beta_{1j}, \beta_{2j})$ can be approximated as a convolution of three mixtures of null and non-null components:

$$
\begin{aligned}
(\beta_{1j}, \beta_{2j}) \approx & \Big[(1-\pi_1)\mathcal{N}(0,0) + \pi_1\mathcal{N}(0,\Sigma_1)\Big] + \\
& \Big[(1-\pi_2)\mathcal{N}(0,0) + \pi_2\mathcal{N}(0,\Sigma_2)\Big] + \\
& \Big[(1-\pi_{12})\mathcal{N}(0,0) + \pi_{12}\mathcal{N}(0,\Sigma_{12})\Big].
\end{aligned} \tag{13}
$$

Second, we assume that LD operator $\mathcal{L}_j = \sum_i \sqrt{H_i} r_{ij}$ is commutative with convolutions in (13). This assumption is accurate for the first two components, $\pi_1$ and $\pi_2$, and is approximate for the last component $\pi_{12}$:

$$
\begin{aligned}
\Big(\frac{\delta_{1j}}{\sqrt{N_{1j}}}, \frac{\delta_{2j}}{\sqrt{N_{2j}}}\Big) = \mathcal{L}_j\Big(\beta_{1j}, \beta_{2j}\Big) \approx & \\
& \mathcal{L}_j\Big[(1-\pi_1)\mathcal{N}(0,0) + \pi_1\mathcal{N}(0,\Sigma_1)\Big] + \\
& \mathcal{L}_j\Big[(1-\pi_2)\mathcal{N}(0,0) + \pi_2\mathcal{N}(0,\Sigma_2)\Big] + \\
& \mathcal{L}_j\Big[(1-\pi_{12})\mathcal{N}(0,0) + \pi_{12}\mathcal{N}(0,\Sigma_{12})\Big].
\end{aligned} \tag{14}
$$

Now we can use Gaussian-type approximation (Theorem 5) or Poisson-type approximation (Theorem 6) to apply LD operator $\mathcal{L}_j$ to each two-component mixture. For Gaussian-type approximation, the result is as follows:

$$
\begin{aligned}
\Big(\frac{\delta_{1j}}{\sqrt{N_{1j}}}, \frac{\delta_{2j}}{\sqrt{N_{2j}}}\Big) \approx & \Big[(1-\pi'_{1j})\mathcal{N}(0,0) + \pi'_{1j}\mathcal{N}(0,\Sigma'_{1j})\Big] + \\
& \Big[(1-\pi'_{2j})\mathcal{N}(0,0) + \pi'_{2j}\mathcal{N}(0,\Sigma'_{2j})\Big] + \\
& \Big[(1-\pi'_{12j})\mathcal{N}(0,0) + \pi'_{12j}\mathcal{N}(0,\Sigma'_{12j})\Big],
\end{aligned} \tag{15}
$$

where

$$\begin{aligned} \pi'_{cj} &= \frac{\pi_c \ell_j}{\pi_c \ell_j + (1-\pi_c)\kappa_j^2}, \quad c \in \{1, 2, 12\}, \\ \Sigma'_{cj} &= \Sigma_c\big((1-\pi_c)\kappa_j^2 + \pi_c \ell_j\big), \\ \ell_j &= \sum_i H_i r_{ij}^2, \quad R_j = \sum_i H_i^2 r_{ij}^4, \quad \kappa_j^2 = R_j/\ell_j. \end{aligned} \tag{16}$$

Ignoring interactions between non-zero components in each mixture, (15) can be approximated by

$$\begin{aligned} \Big(\frac{\delta_{1j}}{\sqrt{N_{1j}}}, \frac{\delta_{2j}}{\sqrt{N_{2j}}}\Big) &\approx \pi'_{0j}\mathcal{N}(0,0) + \pi'_{1j}\mathcal{N}(0,\Sigma'_{1j}) + \pi'_{2j}\mathcal{N}(0,\Sigma'_{2j}) + \pi'_{12j}\mathcal{N}(0,\Sigma'_{12j}), \\ \pi'_{0j} &= 1 - \pi'_{1j} - \pi'_{2j} - \pi'_{12j}. \end{aligned} \tag{17}$$

More accurately (15) can be written as a mixture of 8 components:

$$\begin{aligned} \Big(\frac{\delta_{1j}}{\sqrt{N_{1j}}}, \frac{\delta_{2j}}{\sqrt{N_{2j}}}\Big) \approx\ & \bar{\pi}'_{1j}\bar{\pi}'_{2j}\bar{\pi}'_{12j}\mathcal{N}(0,0) + \\ & \pi'_{1j}\bar{\pi}'_{2j}\bar{\pi}'_{12j}\mathcal{N}(0,\Sigma'_{1j}) + \\ & \bar{\pi}'_{1j}\pi'_{2j}\bar{\pi}'_{12j}\mathcal{N}(0,\Sigma'_{2j}) + \\ & \bar{\pi}'_{1j}\bar{\pi}'_{2j}\pi'_{12j}\mathcal{N}(0,\Sigma'_{12j}) + \\ & \pi'_{1j}\pi'_{2j}\bar{\pi}'_{12j}\mathcal{N}(0,\Sigma'_{1j}+\Sigma'_{2j}) + \\ & \pi'_{1j}\bar{\pi}'_{2j}\pi'_{12j}\mathcal{N}(0,\Sigma'_{1j}+\Sigma'_{12j}) + \\ & \bar{\pi}'_{1j}\pi'_{2j}\pi'_{12j}\mathcal{N}(0,\Sigma'_{2j}+\Sigma'_{12j}) + \\ & \pi'_{1j}\pi'_{2j}\pi'_{12j}\mathcal{N}(0,\Sigma'_{1j}+\Sigma'_{2j}+\Sigma'_{12j}), \end{aligned} \tag{18}$$

where $\bar{\pi}'_{cj} = 1 - \pi'_{cj}$.

To speedup computation of bivariate normal zero-mean denisty one may take advantage of closed-form expression for the matrix inverse. Let variance matrix be $\Sigma = \begin{bmatrix} a & b \\ b & c \end{bmatrix}$, then the precision matrix $\Sigma^{-1} = \frac{1}{ac-b^2}\begin{bmatrix} c & -b \\ -b & a \end{bmatrix}$, and bivariate normal density function

$$\phi(z_1, z_2; 0, \begin{bmatrix} a & b \\ b & c \end{bmatrix}) = \frac{1}{2\pi\sqrt{ac-b^2}} \exp\Big(-\tfrac{1}{2}\frac{cz_1^2 + az_2^2 - 2bz_1z_2}{ac-b^2}\Big). \tag{19}$$

**GWAS power curves.** We are interested in calculating the proportion $S(N)$ of SNP heritability captured by genome-wide significant hits, as a function of GWAS sample size $N$. The $S(N)$ is defined in Online Methods:

$$\begin{aligned} S(N) &= \frac{\sum_j \int_{z:\ |z| \geq z_t} C(z,N,j)dz}{\sum_j \int_z C(z,N,j)dz}, \\ C(z,N,j) &= \int \delta^2 P(z|\delta)P(\delta,j)d\delta. \end{aligned} \tag{20}$$

Whenever $\delta_j$ is modeled as mixture of normal distributions, it is possible to calculate the above integrals as an analytical expression. Let $\delta_j \sim \frac{1}{K}\sum_k N(0, S_{kj}^2)$, where $S_{kj}^2$ is typically a function of $N$, and $z_j \sim \frac{1}{K}\sum_k N(0, \sigma_0^2 + S_{kj}^2)$. Then

$$
\begin{aligned}
C(z, N, j) &= \frac{1}{K}\sum_{k=1}^{K}\int \delta^2 P(z|\delta)P(\delta, j)d\delta = \\
&= \frac{1}{\sqrt{2\pi}K}\sum_{k=1}^{K}\int \delta^2 \frac{1}{\sqrt{2\pi}S_{kj}\sigma_0} e^{-\frac{(z-\delta)^2}{2\sigma_0^2}-\frac{\delta^2}{2S_{kj}^2}} d\delta = \\
&= \frac{1}{\sqrt{2\pi}K}\sum_{k=1}^{K}\frac{S_{kj}^2(\sigma_0^4 + \sigma_0^2 S_{kj}^2 + z^2 S_{kj}^2)}{(\sigma_0^2 + S_{kj}^2)^{\frac{5}{2}}} e^{-\frac{z^2}{2(\sigma_0^2+S_{kj}^2)}},
\end{aligned}
\tag{21}
$$

and

$$
\int_{z:\ |z|\geq z_t} C(z, N, j)dz = \frac{1}{K}\sum_{k=1}^{K}\Big[\frac{\sqrt{\frac{2}{\pi}}S_{kj}^4 z_t}{(\sigma_0^2 + S_{kj}^2)^{\frac{3}{2}}} e^{-\frac{z_t^2}{2(\sigma_0^2+S_{kj}^2)}} + S_{kj}^2 \mathrm{erfc}\Big(\frac{z_t}{\sqrt{2(\sigma_0^2 + S_{kj}^2)}}\Big)\Big],
\tag{22}
$$

where $\mathrm{erfc}(z) = \frac{2}{\sqrt{\pi}}\int_z^\infty e^{-t^2}dt$ is the complementary error function.
